# Exchange, promiscuity, and orthogonality in *de novo* designed coiled-coil peptide assemblies

**DOI:** 10.1101/2024.09.01.610678

**Authors:** Kathleen W. Kurgan, Freddie J. O. Martin, William M. Dawson, Tom Brunnock, Andrew J. Orr-Ewing, Derek N. Woolfson

**Author notes:** Contributed equally to this work (Kathleen W. Kurgan, Freddie J. O. Martin). **Author contributions:** KWK, FJOM, WMD, and DNW conceived the study and designed the peptides and experiments. KWK, FJOM, and TB synthesized the peptides, conducted the experiments, determined the peptide X-ray crystal structures, and analyzed the data with advice from AJO. KWK, FJOM, and DNW wrote the paper. All authors have read and contributed to the preparation of the manuscript.

## Abstract

*De novo* protein design is delivering new peptide and protein structures at a rapid pace. Many of these synthetic polypeptides form well-defined and hyperthermal-stable structures. Generally, however, less is known about the dynamic properties of the *de novo* designed structures. Here, we explore one aspect of dynamics in a series of *de novo* coiled-coil peptide assemblies: namely, peptide exchange within and between different oligomers from dimers through to heptamers. First, we develop a fluorescence-based reporter assay for peptide exchange that is straightforward to implement, and, thus, would be useful to others examining similar systems. We apply this assay to explore both homotypic exchange within single species, and heterotypic exchange between coiled coils of different coiled-coil oligomer states. For the former, we provide detailed study for the dimeric coiled coil CC-Di finding a half-life for exchange of 4.2 ± 0.3 minutes when the concentration of CC-Di is 200 µM. Interestingly, more broadly when assessing exchange across all of the oligomeric states, we find that some of the designs are faithful and only undergo homotypic strand exchange, whereas others are promiscuous and exchange to form unexpected hetero-oligomers. Finally, we develop two design strategies to improve the orthogonality of the different oligomers: (i) using alternate positioning of salt bridge interactions; and (ii) incorporating of non-canonical repeats into the designed sequences. In so doing, we reconcile the promiscuity and deliver a set of faithful homo-oligomeric *de novo* coiled-coil peptides. Our findings have implications for the application of these and other coiled coils as modules in chemical and synthetic biology.

## INTRODUCTION

*De novo* protein design, defined here as designing proteins “from scratch” without starting from a natural protein structure, has become a wide-spread pursuit.^1-4^ Indeed, *de novo* peptides and proteins are impacting areas of cell biology, synthetic biology, nanotechnology, and materials science.^5-10^ Now, computational protein design is accelerating success rates for *de novo* proteins designed with atomic-level accuracy, and the application of AI methods is beginning to democratize protein design making it accessible to experts and non-experts alike.^11-13^ Nonetheless, many challenges remain and harnessing *de novo* protein design to achieve new and effective functionalities is far from routine. A major limitation to computational methods lies in designing proteins with complex energy landscapes similar to those of native proteins.^1-4^ For instance, understanding and controlling the dynamics of *de novo* peptide assemblies and proteins would greatly aid designing functionality and expanding the current boundaries of the field.

While protein design is increasingly relying on computational and AI-based methods,^1-4,11-13^ rational peptide and protein design has also contributed considerable advances in the field.^2,4,10^ *De novo* coiled coils (CCs) are particularly appealing modules for designing self-assembling systems as they are short, mutable, and the principles governing their assembly are well understood.^14-16^ Many groups have delivered robust *de novo* coiled-coil systems, which have been characterized through to atomic structures and used in a wide variety of applications in cell and synthetic biology and in biotechnology.^10,14-16^ Recent in-cell studies have highlighted the potential use of *de novo* coiled-coil design for targeting natural protein-protein interactions^17^ and effecting allosteric activation of these.^18,19^ CC dynamics and specificity are important factors to consider when applying these systems for in-cell purposes, where there is an abundance of native coiled-coil structures and assemblies.^20,21^ The application of peptide-PAINT (points accumulation for imaging in nanoscale topography)^22,23^ and peptide nucleic acid (PNA)^24^ technology to facilitate fluorescence-based imaging of proteins in cells further highlights the potential of dynamic CCs in cell biology. Orthogonal dimeric peptides allow specific labeling or transient association of fluorophores with potentially multiple proteins of interest, and optimization of these platforms has benefited from the ability to tune association of the peptides by altering peptide length and sequence.^22-24^ Yet beyond some examples of studies on dimers^25-27^ and trimers,^28,29^ the dynamics of *de novo* CCs have not been examined in detail.

To date, our lab has elucidated sequence-to-structure rules for defining the oligomeric state of a fleet of *de novo* CCs from dimer to nonomer (Table 1).^30-33^ And along with others,^10,18,34-45^ we have developed rules for controlling topology: parallel vs antiparallel assembly,^17,46,47^ and hetero-peptide association.^17,46,48-50^ Whilst sequence-to-structure relationships governing the oligomeric state, topology and partnering of discrete CC assemblies are well established, the dynamics of these and CC systems in general are less-well explored and understood. The association of parallel heterodimer pairs has been rigorously characterized.^25-27^ However, prior to the work presented here, we have not systematically evaluated the dynamics of exchange and specificity of our *de novo* designs across the whole range of oligomeric states. Therefore, we chose to assess strand exchange using fluorescence as a read out. Wendt et al. have described using a fluorescence-based method to follow the exchange kinetics of peptides based on a designed ⍺-helical leucine zippers.^51,52^ In this method, peptides are appended with an *N*-terminal carboxyfluorescein (FAM) moiety. In solution, the peptides dimerize and FAM self-quenches resulting in low fluorescence emission. When treated with denaturing reagents or unlabeled variants of the peptides, self-quenching is reduced and fluorescence emission increases.^51^ This method allows the observation of kinetics of CC unfolding and strand exchange.

**Table 1.**
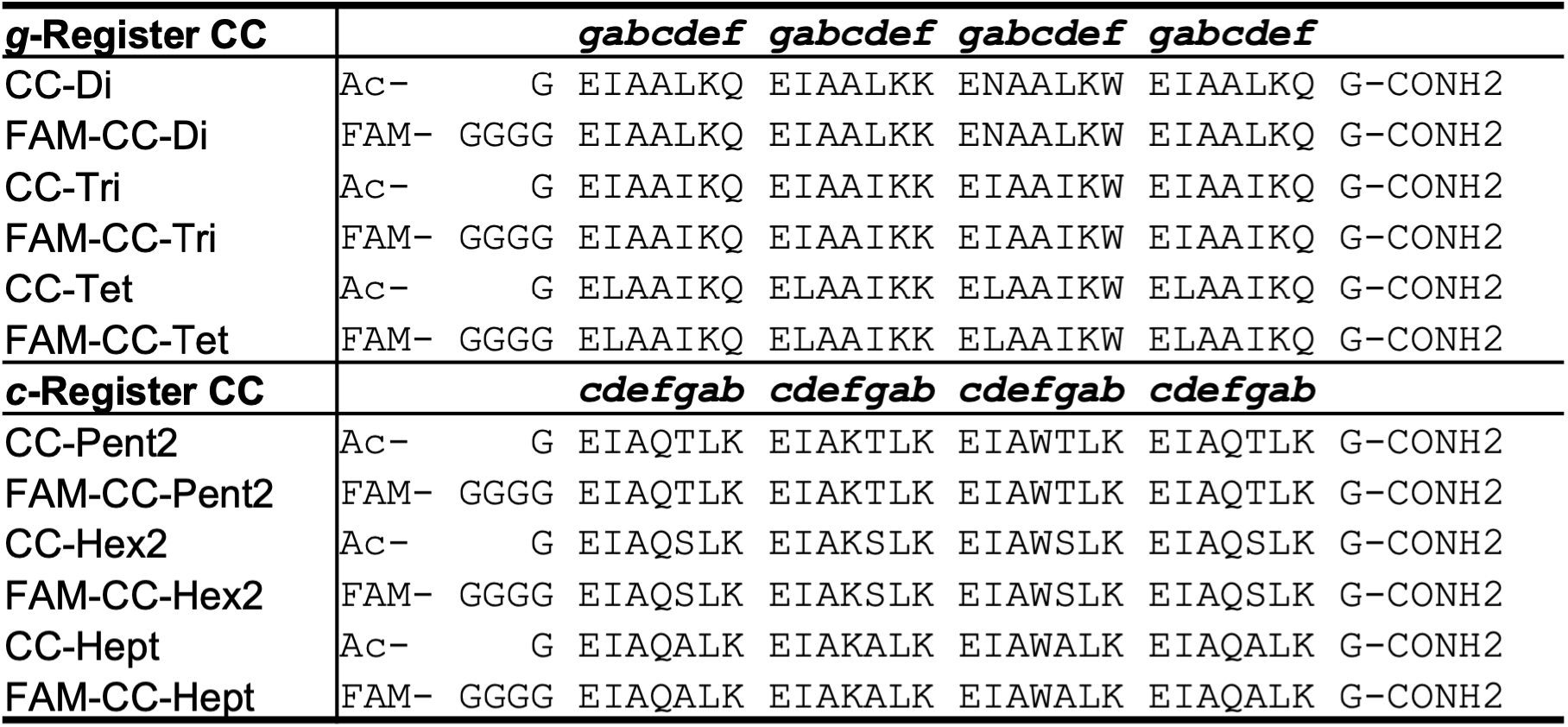
CC Basis Set Peptide Sequences.

Here, we set out to understand the dynamics and specificity of homo- and heterotypic strand exchange in a broader set of *de novo* designed CC peptides. First, we adapt the aforementioned method of Wendt *et al*. to develop a fluorescence-based reporter assay for our own systems. Then, we apply this to study the kinetics and orthogonality of exchange in our published “Basis Set” of *de novo* CC peptides (Table 1).^30-33^ Surprisingly, we find several instances of promiscuous exchange between peptides of different oligomeric state. With this new knowledge, next we generate a set of orthogonal CCs ranging from dimer to heptamer. We use two strategies to increase selectivity in the original CC Basis Set: (i) strategic placement of salt-bridge interactions; and (ii) incorporation of non-canonical, hendecad repeats, in the designed sequences. In these ways, we identify a set of CCs that show little to no heterotypic strand exchange. On this basis, we call these peptides an “Orthogonal CC Basis Set”. The intention is that these can be used in concert with each other (mixed and matched) to drive complex specific protein assemblies whilst minimizing off-target interactions for applications in chemical and synthetic biology.^10,16,18^

## RESULTS AND DISCUSSION

### A reporter system for coiled-coil strand exchange

To explore the dynamics of strand exchange in and between *de novo* coiled-coil peptide assemblies, we sought a minimally invasive assay that could be performed in medium-to-high throughput. We chose fluorescence measurements, and the strategy depicted in Figure 1A. Using a similar experimental design to Wendt *et al*.,^51^ CC peptides were synthesized in two forms: one with an *N*-terminal 5(6)-carboxyfluorescein (FAM) moiety (the ‘labeled’ peptide) and the other unlabeled. FAM self-quenches at concentrations above 10 mM.^53^ We reasoned that parallel homo-oligomerization of labeled peptides would bring two or more FAM moieties to within ≈1 nm, increasing the FAM effective local concentration to ≈1 M and thus quenching the fluorescence. However, if peptides can freely exchange between assembled coiled coils, the addition of an unlabeled peptide in excess should result in hetero-complexes of the labeled and unlabeled peptides and, so, reduce self-quenching leading to a fluorescence signal measured over time (Figure 1A).^51^

**Figure 1.**
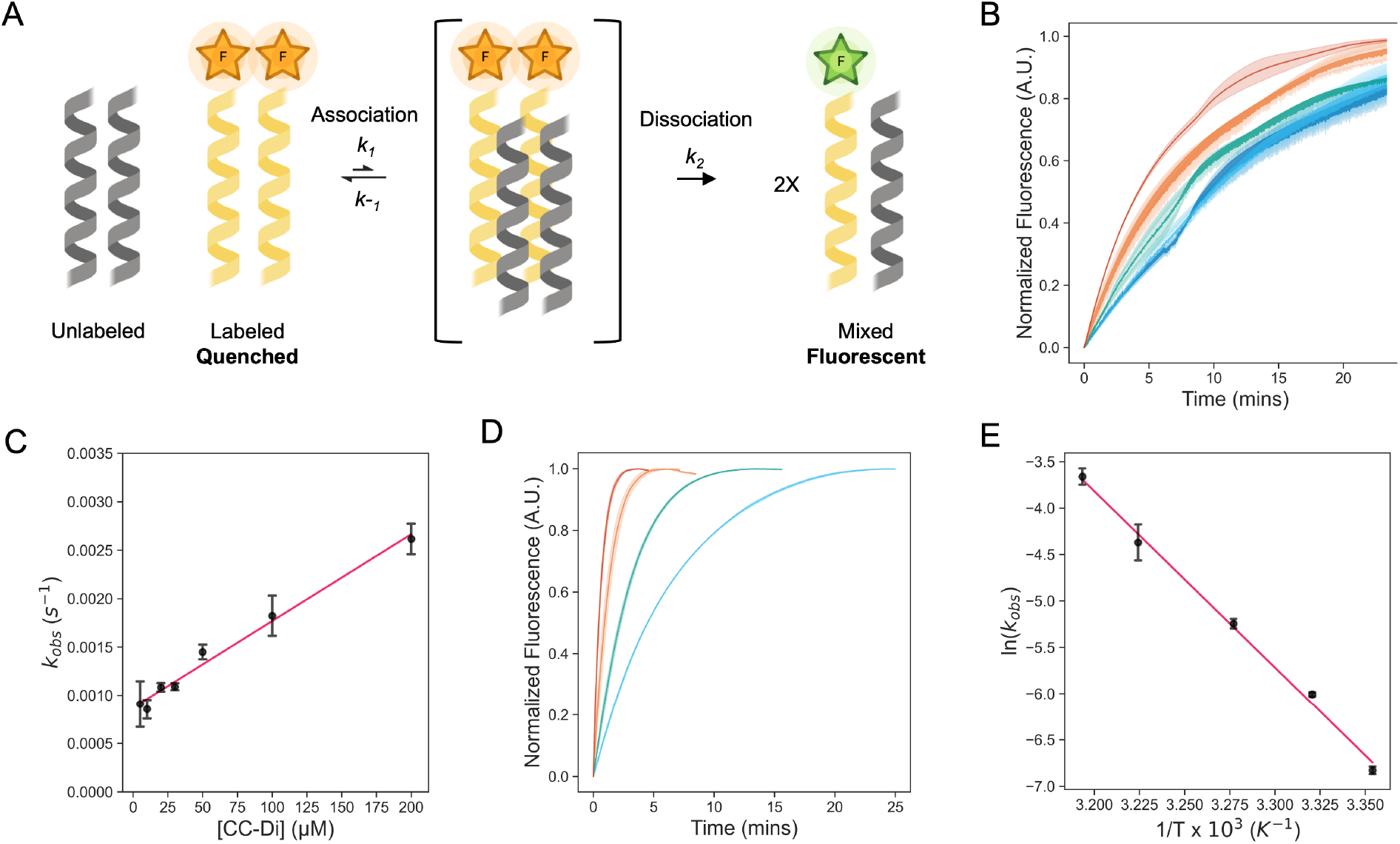
Assessing strand exchange of CC-Di using fluorescence-based measurements. (**A**) One possible scheme of the exchange between a quenched labeled and an unlabeled CC dimer, via a tetrameric steady-state intermediate, to form the fluorescent mixed species. The scheme was created with Biorender.com. (**B**) Normalized fluorescence time course plots for the exchange of CC-Di at different concentrations of unlabeled CC-Di. The experiments were carried out at 2 µM of labeled CC-Di. The plots are colored by the concentration of CC-Di: blue, 20 µM; cyan, 30 µM; green, 50 µM; orange, 100 µM; and red, 200 µM. (**C**) Plots of the observed pseudo-first-order rate constant (k_obs_) for exchange at different concentrations of unlabeled CC-Di (2 – 200 µM). Data points are shown as the average of 3 independent replicates, error bars are for 1 standard deviation, and the line of best fit is shown. (**D**) Normalized fluorescence time course plots for the exchange of CC-Di at different temperatures (28, 32, 37, and 40 °C). The experiments were carried out at 2 µM of labeled CC-Di and 20 µM of unlabeled CC-Di. The plots are colored by the temperature: blue, 28 °C; green, 32 °C; orange, 37 °C; and red, 40 °C. (**E**) Arrhenius plot for the temperature dependence of the rate constants for exchange of CC-Di. Values determined from fits to data shown in Figures S13-16 and Table S1. Errors are shown to one standard deviation of independent triplicate measurements. All experiments were carried out at 25 °C in phosphate buffered saline (PBS) at pH 7.4 unless otherwise stated. Observed rate constants (k_obs_) were determined by fitting the normalized fluorescence time course profiles to an exponential rise (Supplemental Information Equation 2).

To test this approach, we selected our simplest *de novo* CC, the parallel homodimer, CC-Di,^31^ synthesizing it in the two forms (Table 1). Previously, Wendt *et al*. have determined dissociation constants (*K*_d_) of similarly labeled and unlabeled CC dimers using fluorescence, chemical denaturation, and circular dichroism measurements. The resulting *K*_d_ values are consistent across the different methods, suggesting that association/dissociation are not perturbed significantly by labeling with FAM.^51^ Nonetheless, fluorescein has been observed to promote aggregation or enhance the stability of coiled-coil assemblies.^54^ Therefore, we tested for any influence of the fluorophore on CC association in our system by assessing different linkers between the FAM and the CC sequence.^55^ We found four Gly residues to be the simplest linker that gave consistent results in the following kinetic experiments.^55^ Next, we explored various ratios of labeled and unlabeled peptides to achieve high fluorescence values upon exchange, settling on a 1:10 ratio of the assembled species, as opposed to the previously used 1:1 ratio.^51^ For instance, for CC-Di, this was 2 µM of labeled and 20 µM of unlabeled peptide, or 1 µM:10 µM of the dimeric assemblies. Under these conditions, we observed a rapid increase in fluorescence upon mixing the two peptides, Figure 1B, indicating rapid strand exchange between the folded labeled and unlabeled species.

Subsequently, we performed kinetic experiments varying labeled to unlabeled peptide ratios to probe the mechanism of exchange for CC-Di. First, we kept the concentration of unlabeled peptide constant (200 µM) and varied the labeled peptide concentration (1 – 20 µM), Figure S17. The observed rate constants changed little. Under these conditions, with the unlabeled peptide in excess, we expected and observed no correlation between the change in the rate constant and change in concentration of labeled peptide (see SI for details). Second, we kept the labeled peptide constant (2 µM) and varied the unlabeled peptide (2 – 200 µM), Figure 1B&C. In this case, the rate constant increased linearly with concentration of the unlabeled peptide. This is consistent with the pseudo-first order kinetics expected with an excess of unlabeled peptide over labeled peptide. While the mechanism of exchange of FAM-CC-Di and CC-Di cannot be elucidated from these data, we propose some possible mechanisms of exchange (Figure 1A, Scheme S3,S4). For instance, Figure 1A shows one mechanism where: (1) association of labeled and unlabeled dimers form a tetrameric steady-state intermediate; which is followed by (2) dissociation of the intermediate to form fluorescent dimers composed of one copy of FAM-CC-Di and one copy of CC-Di. We posit that this is more likely to occur than the other proposed mechanisms, which are initiated by dimer dissociation, as the folded FAM-CC-Di and CC-Di dimers will be in great excess compared to dissociated monomeric variants (Tables S1 and S2). This is because the experiments are conducted at µM peptide concentrations, but the dissociation constant of CC-Di is sub-nM.^31^ Based on conditions of pseudo-first-order reaction kinetics, we fitted the kinetic transients to single exponential curves to obtain rate constants and calculate half-lives for the strand exchange (*t*_*1/2*_ = ln2/*k*_*obs*_). The calculated half-life for strand exchange in 25 °C conditions where the CC-Di concentration is 200 µM was *t*_*1/2*_ = 4.2 ± 0.3 minutes. This is longer than for recently examined heterodimeric *de novo* CCs, with *t*_*1/2*_ ≈ 7 – 70 seconds,^27^ though these have a range of *K*_*D*_ values (8.1 x 10^−9^ – 3.8 x 10^−5^ M) that are weaker than CC-Di. Finally, as expected, the exchange rate constant increased with temperature, Figure 1D, and an Arrhenius analysis returned an activation enthalpy of 37.9 ±1.9 kcal (mol of dimer)^-1^, Figure 1E. This is comparable to the activation energy determined for the unfolding of the GCN4-p1 leucine-zipper dimer, 30.8 kcal (mol of dimer)^-1^.^56^ We conducted similar kinetic strand-exchange experiments for the other *de novo* designed CC Basis Set peptides, *i.e*., CC-Tri through CC-Hept.^16,30-33^ Although these all showed increases in fluorescence consistent with strand exchange to produce mixed labeled/unlabeled species, the kinetic mechanisms for exchange in these higher-order oligomers are more complicated than those shown in Figure 1 and Schemes S3&S4 for CC-Di.^55^ As such, these data could not be fitted using simple rate equations. This is because, rather than one dominant transient species and a single mixed fluorescent equilibrium species, many species with different numbers of labeled (l) and unlabeled (u) peptides are possible, and the number of combinations increases with increasing oligomer state. For instance, for CC-Tri there are 4 assembled parallel species alone: u-u-u, l-l-l, u-u-l, and u-l-l. This led us to conclude that, for the larger oligomer states, rather than following and quantifying the kinetic traces directly, we needed to focus on the endpoints in the exchange experiments.

### Homotypic and heterotypic strand exchange for the CC Basis Set

With the above in mind, we adopted the approach of monitoring the extent of fluorescence (again, as a proxy for exchange) at three points as shown in Figure 2A. This involved mixing the labeled and unlabeled peptides (with the assumed assemblies at 1 µM and 10 µM, respectively) under folded conditions incubated at 25 °C. Aliquots were taken for fluorescence measurements at 1 hr and 24 hr time points. Then, the remaining samples were heated to 95 °C for 5 minutes and cooled to 25 °C over 2 hours. Regardless of differences in kinetics, this final annealing step aimed to facilitate any further possible strand exchange prior to recording the final fluorescence signal.

**Figure 2.**
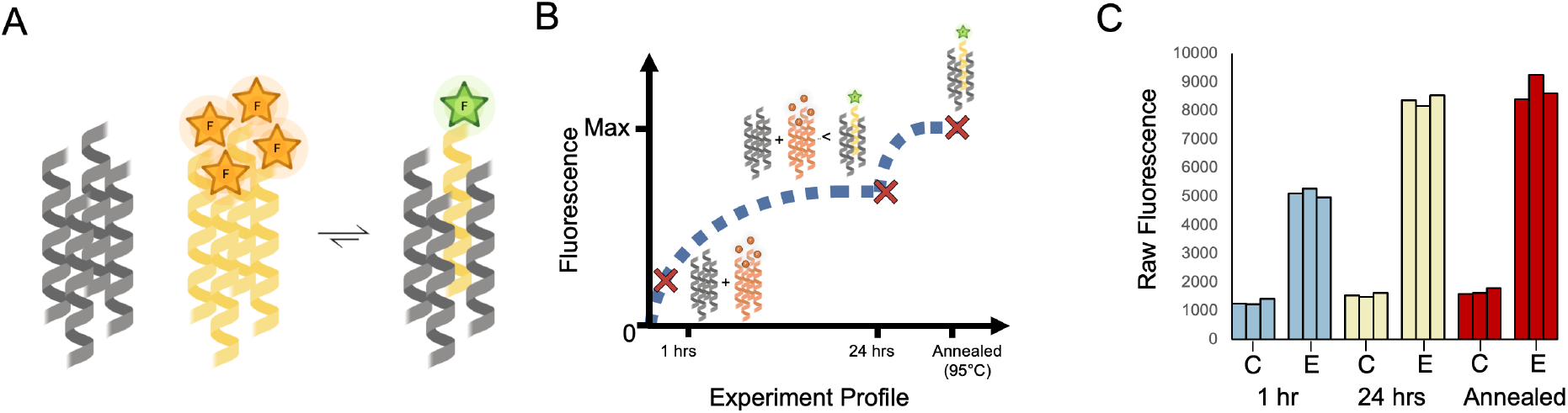
Homotypic exchange of CC-Tet. (**A**) Cartoon for the assumed equilibrium reached when labeled and unlabeled variants of CC-Tet are mixed in a 1:10 ratio. For simplicity, we have omitted the excess of the unlabeled peptide and any intermediate species. This cartoon was made with Biorender. **(B)** Cartoon for the anticipated changes in population of the quenched and free FAM-CC-Tet over time where the major species are represented at different time points. (**C**) Raw data from experiments following the protocol outlined in the text and depicted in panel B. (See SI for experimental details.) The data were collected for two different samples: (1) a control sample, represented as “C,” which corresponds to 2 µM FAM-CC-Tet in PBS and (2) an exchange sample, represented as “E,” which corresponds to 2 µM FAM-CC-Tet + 20 µM CC-Tet in PBS. The fluorescence measurements were collected at three different time points (1 hr, 24 hr, and after annealing) and repeated 3 times. All fluorescence data of FAM-CC-Tet mixtures were subsequently normalized against the data collected for the annealed samples where the averaged observed fluorescence of FAM-CC-Tet in buffer was set to zero and the averaged observed fluorescence of 1:10 ratio of FAM-CC-Tet to CC-Tet was set to one.

Taking the CC-Tet homomeric exchange as an example, we anticipated exchange between labeled and unlabeled variants of CC-Tet and, with an excess of unlabeled peptide, the equilibrium should shift toward a population of unquenched FAM (Figure 2A). Depending on the rate of exchange, it would take time for the mixture to reach this equilibrium. Earlier time points would still have a high population of unmixed FAM-labeled peptide, corresponding to lower fluorescence values. Indeed equilibrium, and consequently high fluorescence values, may only be reached after annealing (Figure 2B). This anticipated behavior was apparent in the fluorescence data (Figure 2C). For the FAM-labeled peptide alone in buffer, the fluorescence changed little over time (Figure 2C). However, the mixture of labeled and unlabeled CC-Tet gave comparatively high fluorescence values at the 1 hr and 24 hr time points, with the latter comparable to the signal after annealing. These data illustrate the utility of the endpoint method for assessing strand exchange in CC systems.

With this method in hand, we assessed all combinations of labeled and unlabeled peptides for the whole CC Basis Set (Table 1). We applied min-max scaling to the raw fluorescence data using the averaged fluorescence value of the pure annealed labeled peptide as the minimum value, and that for the annealed 1:10 mixture of the homotypic exchange that corresponds to the labeled peptide as the maximum, Figure 3. High fluorescence values, *i.e*. high amounts of exchange, are shown in red, and low fluorescence/exchange is in blue. In comparison to homotypic exchange, it is less clear what the composition of the peptide mixtures, the oligomeric state and the stoichiometry of potential mixed assemblies, will be at equilibrium. The mechanism of exchange Is also less obvious. For these reasons, we cannot expect to see the same trends (increasing fluorescence over time followed by maximum fluorescence after annealing), that we observe in the homotypic exchange experiments. As annealing allows the systems to reach a thermodynamic minimum, these vales best represent the possible full level of exchange.

**Figure 3.**
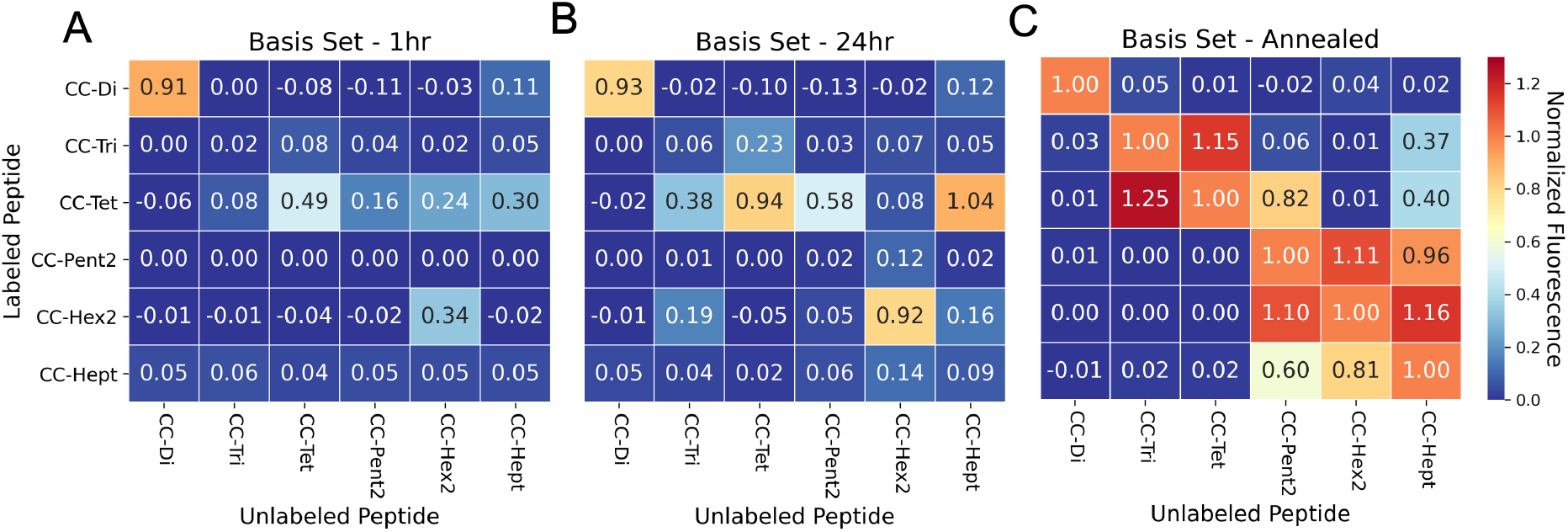
Summary of exchange across the whole CC Basis Set. These heat maps show the normalized fluorescence values for all peptide mixtures. Measurements were taken at 1 hr (**A**) and 24 hrs (**B**) after mixing, and after a subsequent annealing step (**C**). With a few exceptions, the post-annealed values are much higher than those observed after 1 hr and 24 hrs. The post-annealed data set shows exchange within the homotypic mixtures (diagonal values), and exchange between CC-Tri and CC-Tet and between Type II coiled coils (CC-Pent 2, CC-Hex 2, and CC-Hept).

Focusing on the diagonal in Figure 3A, after a 1-hour incubation at 25°C CC-Di had undergone significant homotypic exchange and CC-Tet and CC-Hex2 had partly exchanged with their labeled variants, whereas, CC-Tri, CC-Pent2, and CC-Hept had exchanged little. These data are consistent with the thermal stabilities of the peptides assemblies: CC-Di has the lowest reported T_M_ (78°C) of the CC Basis Set, whereas the others are hyper-thermostable and do not unfold completely upon heating to 95°C. ^30-33^ Moreover, from the off-diagonal cells, there were signs of heterotypic exchange; for instance, between labeled CC-Tet and the unlabeled higher-order CC-Pent2, CC-Hex2, and CC-Hept. After 24 hours at 25 °C (Figure 3B), these trends were accentuated, although for some peptides (*e.g*. CC-Tri, CC-Pent2, and CC-Hept) little homotypic exchange had occurred. As expected, the heating-and-cooling step increased exchange across the whole set of combinations, and further highlighted this potential for ‘promiscuous’ heterotypic exchange (Figure 3C). For example, CC-Tri and CC-Tet exchanged with each other, as did the α-helical barrels, CC-Pent2, CC-Hex2, and CC-Hept.

The key results and our interpretations from these experiments on mixing the original CC Basis Set peptides are as follows. (1) CC-Di only undergoes homotypic exchange; *i.e*., it is a faithful design that is orthogonal to the other *de novo* CCs (top left cell of Figure 3C). This is likely because it is the only design with a buried polar residue—an Asn at the central ***a*** site, Table 1—incorporated to specify the parallel dimer.^31^ Thus, exchange with any other CC, which have exclusively hydrophobic cores, would be energetically unfavorable. (2) CC-Tri and CC-Tet exchange with each other, but less so with the higher-order CCs, (although labeled CC-Tri and CC-Tet do exchange with unlabeled CC-Pent2 and CC-Hept, and with CC-Hept, respectively, to some extent (Figure 3)). We propose that this is because CC-Tri and CC-Tet have similar heptad repeats, `E`***a***`AAIKX` in ***g****⟶****f*** register, with only the residues at ***a*** being different (***a*** = Ile in CC Tri, and Leu in CC Tet).^31^ This opens possible CC-Tri:CC-Tet heterotypic interactions that we had not considered before conducting these fluorescence-based exchange experiments. Finally, (3) the higher-order CCs, CC-Pent2, CC-Hex2, and CC-Hept, also interact with each other. However, this is to differing degrees and is only significant after heating and cooling (compare the bottom-right quadrants of the three panels of Figure 3). Again, the peptide sequences are key to understanding this. These peptides are Type-II CCs in which residues at ***g, a, d***, and ***e*** sites engage in helix-helix interactions.^16,30,32,33,57^ Moreover, the ***g****⟶****f*** heptad repeats are similar, *i.e*. ***g***LKEIAX with ***g*** = Thr for CC-Pent2, Ser for CC-Hex2, and Ala for CC-Hept. Overall, whilst these subtle changes demonstrably give oligomer-state specificity to the homomers,^30,32,33^ there is unforeseen potential for heterotypic interactions.

### Establishing orthogonal trimeric and tetrameric coiled coils

Given these new insights into the CC Basis Set from the fluorescence-exchange experiments and the need for orthogonal components for applications in chemical and synthetic biology, we sought to improve the orthogonality of the more-promiscuous CCs using rational redesign. Our aim was to make minimal mutations that would not compromise the CC oligomer state or stability. We started with the CC-Tri and CC-Tet sequences, reasoning that salt-bridge interactions could be exploited to achieve orthogonality between these trimeric and tetrameric CCs. This was prompted by a variant of the latter, CC-Tet*,^58^ where interhelical salt-bridging lysine residues and glutamates are moved to the ***b*** and ***c*** positions, as compared with in the ***e*** and ***g*** positions in the original CC-Tri and CC-Tet sequences (Figures 4A&B). CC-Tet* is more robust as a parallel tetramer than the original CC-Tet design; minor variations to the latter can result in trimers.^31,58^ Therefore, we tested the exchange between CC-Tri and CC-Tet* (Figure 4C). We observed lower fluorescence values for the mixed heterotypic exchange in comparison to high fluorescence values of the homotypic exchange and the CC-Tri:CC-Tet experiments (compare Figures 4C and 3). These results suggest that orthogonality can be driven by strategic positioning of salt-bridge interactions.

**Figure 4.**
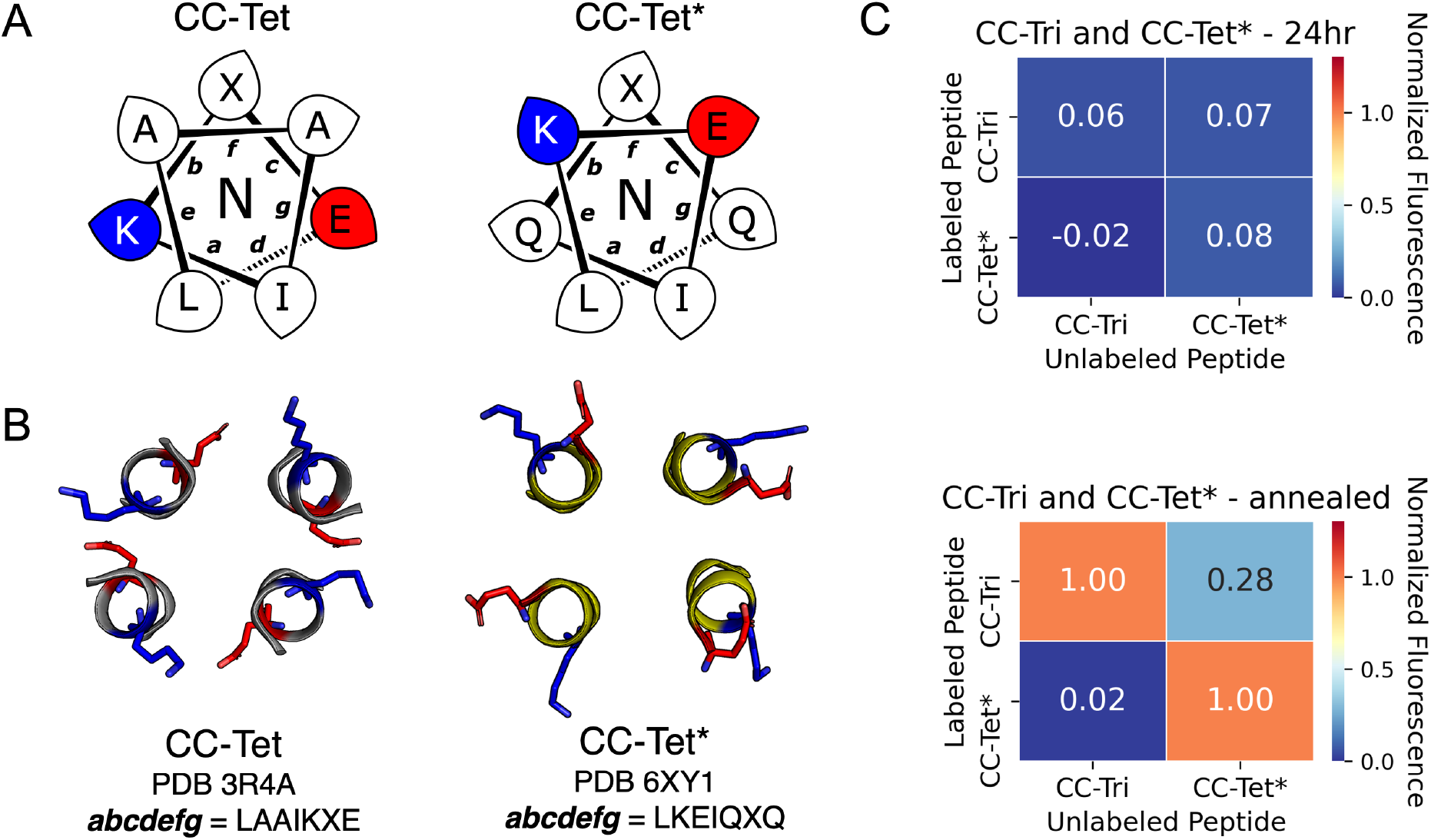
Improving orthogonality between CC-Tri and CC-Tet. (**A**) Helical wheel representations of the heptad-repeat sequences of CC-Tet and CC-Tet*. In CC-Tet, residues that promote interhelical salt-bridge interactions (lysine and glutamic acid) are positioned at the ***e*** and ***g*** positions, whereas in CC-Tet* these are at ***b*** and ***c***. (**B**) Structural consequences of the different placements of these residues. (**C**) Normalized fluorescence values for exchange between CC-Tri and CC-Tet*. After annealing, values for the hetero-mixtures (off diagonal) are much lower than those for the homomeric samples, indicating orthogonality over the CC-Tri:CC-Tet combination (Figure 3C).

### Designing orthogonal higher-order coiled coils using alternative sequence repeats

For the ⍺-helical barrels, CC-Pent2, CC-Hex2, and CC-Hept, designing orthogonal variants was more challenging. Shifting the salt-bridge interactions was not an option as these were already at ***b*** and ***c*** in all sequences (Table 1). Additionally, apart from pentamers, where glutamine can be introduced into the ***a***/***d*** cores,^55,59-61^ there are few examples to guide the rational design of polar layers in these higher-order CCs. Therefore, we took a different approach of incorporating non-heptad sequence repeats into these peptide sequences. Canonical heptad repeats space core-facing side chains 3 and 4 residues apart at the ***a*** and ***d*** sites (Figure S38). Variations of this 3-4 spacing can lead to alternative CC sequences and structures. For instance, 11-amino acid, hendecads have core residues spaced 3,4,4 apart at the ***a, d***, and ***h*** sites of ***a****⟶****k*** repeats (Figure S38).^16,62-66^ A consequence of this is that while heptad repeats drive association of left-handed CCs, hendecads produce right-handed structures (Figure S38). ^16,62-66^ We reasoned that replacing heptad with hendecad repeats at different locations of the pentamer to heptamer sequences would alter the hydrophobic seam, which, in turn, might confer homo-specificity in the resulting sequences.

Initially, we made several variants of CC-Pent2, CC-Hex2, and CC-Hept that extended the second heptads into hendecad repeats (Table 2). To do this, the original ***a*** – ***g*** sequences were maintained, and the new ***i*** – ***k*** sites were all made Ala. What to place at the core-forming ***h*** sites was less clear, so we tested Ala, Ile, and Leu in each of the three designs. These were tested in the endpoint fluorescence assay against their FAM-labeled parent, heptad-based peptides (Figures 5A – C). We reasoned that any orthogonality in these experiments would be an indicator of potential orthogonality with the other oligomeric states. Generally, for the CC-Pent2 and CC-Hept designs, these experiments revealed less exchange between the heptad and hendecad variants compared to the homotypic exchange of the parents, indicating that the strategy had indeed increased orthogonality. However, the experiments with CC-Hex2 showed considerable exchange and therefore promiscuity between all pairings. To investigate this further, we compared the structural data of CC-Pent2-hen2, CC-Hex2-hen2, and CC-Hept-hen2 all with Ala at the ***h*** position to see if our hypothesis, that the hydrophobic seam would be altered in comparison to the parent CC assemblies, was correct. The AlphaFold2^67^ model of CC-Pent-hen2 with Ala at ***h*** (Figure 5D) displayed a marked kink in the monomeric helix and the crystal structure of CC-Hept-hen2 with Ala at ***h*** (Figure 5F) showed a more subtle kink. This feature is consistent with combining left- and right-handed CC repeats. Yet the crystal structure of CC-Hex2-hen2 with Ala at ***h*** revealed a similar supercoiling to its parent despite the incorporation of the hendecad repeat (Figures 5E), possibly explaining the extended promiscuity observed between the CC-Hex2 variants.

**Table 2.**
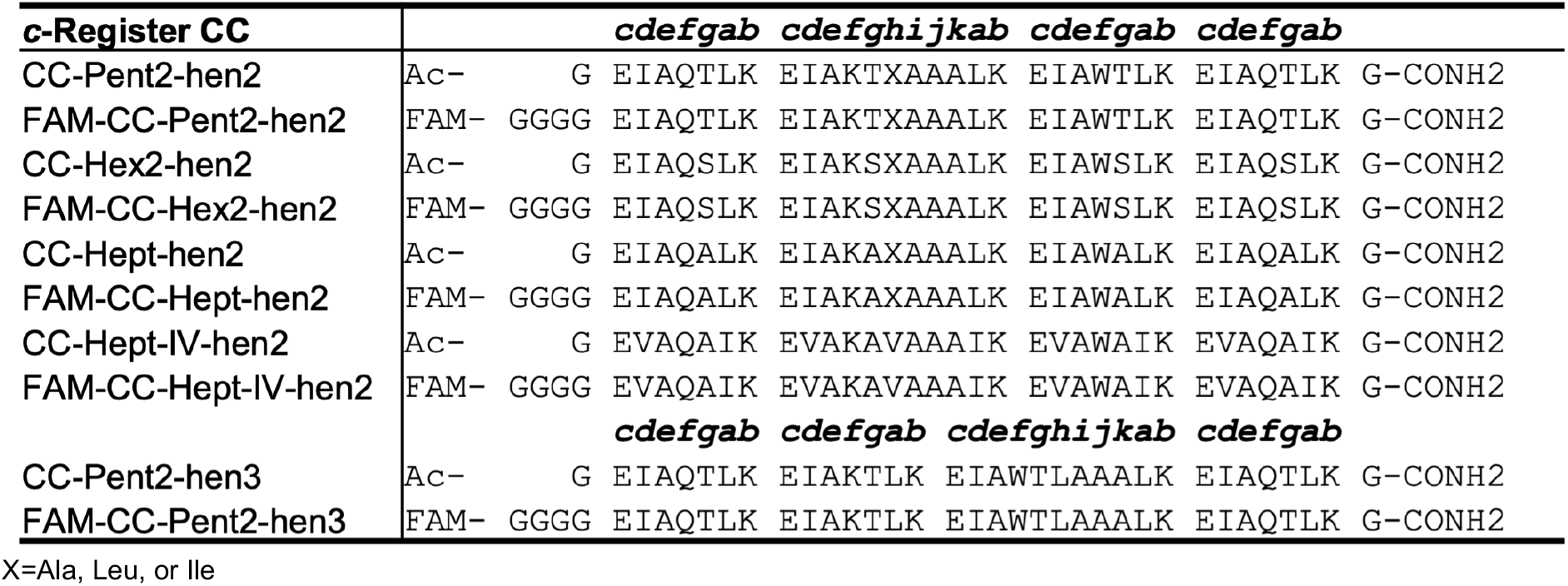
Hendecad incorporated variants of the ⍺-helical barrels, CC-Pent2, CCHex2, and CC-Hept. X=Ala, Leu, or Ile.

**Figure 5.**
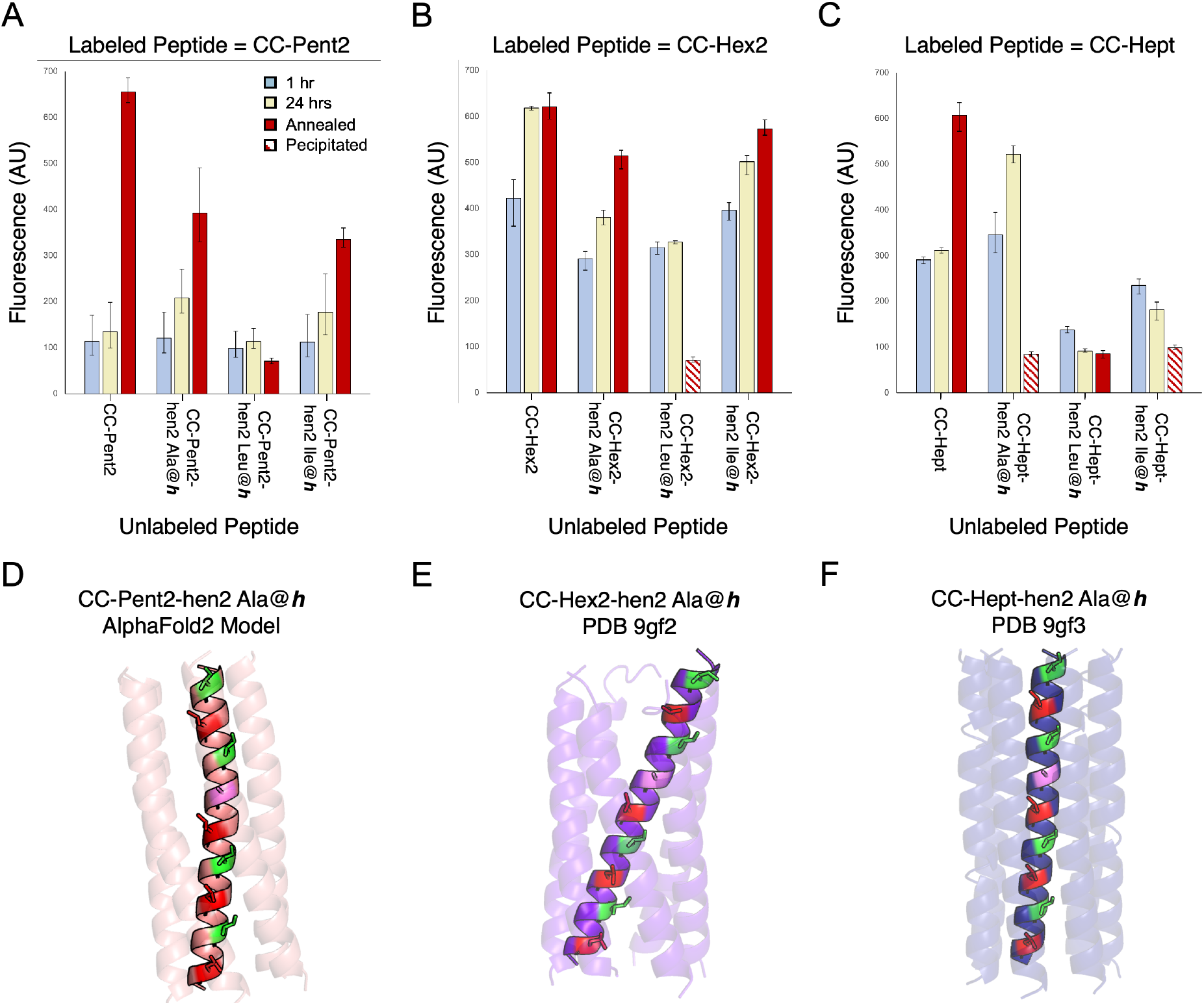
Mixing heptad and hendecad repeats to improve orthogonality. (**A** – **C**) Fluorescence-based orthogonality screens of ⍺-helical-barrel variants incorporating hendecad repeats, CC-Pent2-hen2, CC-Hex2-hen2, and CC-Hept-hen2, where the h positions were Ala, Leu, or Ile. This initial screen was performed against FAM-labeled variants of the parent, heptad-based peptides. Compared with homotypic controls (*i.e*. FAM-CC-Pent2 + CC-Pent2, FAM-CC-Hept + CC-Hept), the CC-Pent2-hen2 and CC-Hept-hen2 variants showed marked decreases in fluorescence indicating less exchange and thus improved orthogonality (D & F). However, the CC-Hex2 variants still showed some cross-exchange and promiscuity (E). Note: Some of the hen2 variants were not stable up to 95°C and precipitated from solution after annealing as indicated by the striped data columns. (**D**-**F**) An AlphaFold2^67^ model and X-ray crystal structures of CC-Pent2-hen2, CC-Hex2-hen2, and CC-Hept-hen2 (all with Ala at ***h***) respectively are shown with the ***a*** positions colored red, ***d*** green, and ***h*** in lilac.

### An orthogonal Coiled-coil Basis Set

Our final goal was to achieve as much orthogonality across a revised CC Basis Set as possible. Based on the above experiments collectively, we reasoned that the original CC-Di, CC-Tri, and CC-Hex2 sequences could be kept, and that CC-Tet* could be substituted for CC-Tet. For CC-Pent2 and CC-Hept, initially, we chose to include hendecad repeats in place of the third and second heptads, respectively, to give CC-Pent2-hen3 and CC-Hept-hen2 taking the Leu-at-***h*** variants for both (Table 2). Unfortunately, FAM-CC-Hept-hen2 was not soluble in PBS buffer, so we switched to a heptamer sequence with Ile in the ***a*** sites and Val at ***d*** and ***h*** (CC-Hept-IV-hen2, Table 2). This proved more soluble and retained orthogonality. CC-Pent-hen3 was confirmed as a pentamer by analytical ultracentrifugation (AUC) experiments (Table S9, Figure S48). CC-Hept-IV-hen2 sedimented as a hexamer in AUC (Table S9, Figure S49), but an X-ray crystal structure revealed a heptamer (Figure 6A, Table S8). Finally, we tested all the proposed revised Basis Set, denoted Orthogonal CC Basis Set, in the endpoint fluorescence assay (Figures 6B – D). Although, some promiscuity remained between the tetra-, penta- and hexameric CCs, overall, this showed significant improvements in orthogonality across the whole set compared with the original Basis Set going into this study (compare with Figure 3).

**Figure 6.**
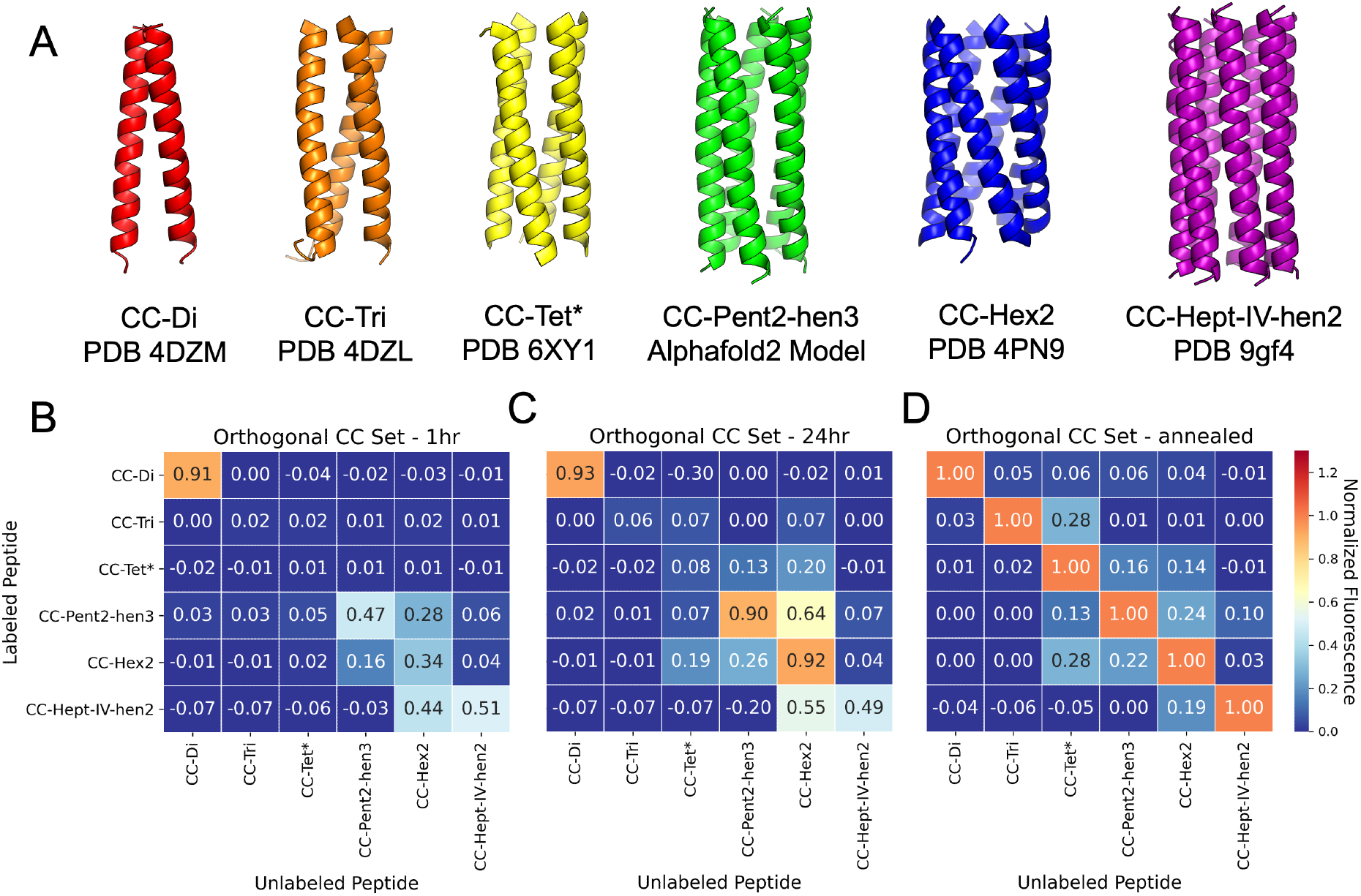
An Orthogonal CC Basis Set. (**A**) X-ray crystal structures and an AlphaFold2^67^ model for the assembled peptides in this set. The pentamer and heptamer were designed in this study, the other peptides have been published elsewhere.^30-33,58^ (**B – D**) Heat maps from the fluorescence exchange data. The normalized fluorescence values of the mixed peptides show some evidence of association between non-homomeric couples at the 1 hr (**B**) and 24 hr (**C**) time points. However, the post-annealing values (**D**) indicate considerable orthogonality across the new set: the homotypic exchange (diagonal) values are all higher than those for any of the heterotypic exchange experiments (off-diagonal). All mixtures containing CC-Hept-IV-hen2 and/or FAM-CCHept-IV-hen2 were annealed to 75 °C instead of 95 °C as these peptides were not stable up to 95 °C (see Figure S30).

## CONCLUSION

Here, we develop a method for assessing association between coiled-coil peptides using a fluorophore reporter strategy. This is applied to assess homo- and heterotypic strand exchange between a published set of *de novo* coiled coils ranging in oligomeric state from dimer to heptamer.^30-33^ We observe that while the dimer, CC-Di, is completely orthogonal, coiled-coil designs with similar helix-helix interface but different oligomeric states can exchange promiscuously. For example, the trimer, CC-Tri, and the tetramer, CC-Tet, which have Type I interfaces (meaning that the ***g, a*** and ***d*** residues contribute to the helix-helix interactions), exchange with one another. Similarly, peptides with Type II interfaces (where the ***g, a, d***, and ***e*** residues are all engaged in helix-helix interactions), namely the pen tamer, hexamer, and heptamer, CC-Pent2, CC-Hex2 and CC-Hept, also exchange with each other. For applications of the original CC Basis Set, whilst this may not be an issue when using designs on their own, or paired with orthogonal designs (*e.g*, CC-Di with CC-Tri), our aim was to achieve a set that is as orthogonal as possible and can be used in many combinations. To address this, and using the insight from the data and analysis presented here, we test two different strategies for increasing specificity of coiled coils: (i) alternate placement of salt bridge-promoting residues, lysine and glutamate to discriminate the tetramer from the trimer; and (ii) incorporation of hendecad repeats to separate the higher-order Type II assemblies. The first strategy was successful as demonstrated by the low heterotypic exchange between CC-Tri (with lysine and glutamate at the ***e*** and ***g***, respectively) and CC-Tet* (where lysine and glutamate are moved to ***b*** and ***c***, respectively).^58^ The second strategy differentiates the ⍺-helical barrels, CC-Pent2, CC-Hex2, and CC-Hept, by introducing hendecad repeats in a pentamer and heptamer to give CC-Pent2-hen3 and CC-Hept-IV-hen2. A search for hendecad repeats in structurally validated CC assemblies of the CC+ Database gave only one example of a pentameric assembly and no examples of heptameric assemblies.^68^ However, sequence analysis and model predictions suggest that higher oligomeric CC assemblies with hendecad repeats do occur in nature.^66^ Fluorescence measurements showed minimal exchange between CC-Di, CC-Tri, CC-Tet*, CC-Pent2-hen3, CC-Hex2, and CC-Hept-IV-hen2. We have denoted this group of peptides as the Orthogonal Coiled-coil Basis Set. Orthogonal CC dimers have been used to design macromolecular assemblies,^69-72^ protein-protein interactions in cells,^17-19^ and scaffolds for specific fluorophore labeling of proteins of interest.^22-24^ In a similar vein, we envision this wider set as building blocks for protein design for applications in chemical and synthetic biology.

## ACKNOWLEDGMENTS

KWK and DNW were funded by a BBSRC-NSF grant (BB/V004220/1). FJOM and DNW were funded by a BBSRC grant (BB/R00661X/1) to DNW. FJOM was also supported by the Bristol Chemical Synthesis Centre for Doctoral Training funded through the EPSRC (EP/G036764). DNW was also supported by BrisEngBio, a BBSRC-funded Engineering Biology Research Centre (BB/L01386X/1), a Royal Society Wolfson Research Merit Award (WM140008), and a Biotechnology and Biological Sciences Research Council (BBSRC) grant (BB/S002820/1). AJOE thanks ESPRC for support through grant EP/V026690/1. We thank the University of Bristol, School of Chemistry, Mass Spectrometry Facility for access to the EPSRC-funded Bruker Ultraflex MALDI-TOF instrument (EP/K03927X/1). We would like to thank Diamond Light Source for access to beamlines I04 and I24 (Proposal mx23269). We thank all members of the Woolfson laboratory for helpful discussions.

## Notes

### Competing Interest Statement

The authors have declared no competing interest.

