## Supplemental Information for "Exchange, promiscuity, and orthogonality in *de novo* designed coiled-coil peptide assemblies"

### Supporting Information for: Exchange, promiscuity, and orthogonality in *de novo* designed coiled-coil peptide assemblies

Kathleen W. Kurgan,[1,†] Freddie J. O. Martin,[1,†] William M. Dawson,[1] Thomas Brunnock,[1] Andrew J. Orr-Ewing,[1] and Derek N. Woolfson[1,2,3,4*]

[1]School of Chemistry, University of Bristol, Cantock's Close, Bristol BS8 1TS, UK

[2]School of Biochemistry, University of Bristol, Medical Sciences Building, University Walk, Bristol BS8 1TD, UK

[3]Bristol BioDesign Institute, School of Chemistry, University of Bristol, Cantock's Close, Bristol BS8 1TS, UK

[4]Max Planck-Bristol Centre for Minimal Biology, University of Bristol, Cantock's Close, Bristol BS8 1TS, UK

[†]Contributed equally to this work.

**Methods**……S2-S6
- **General**……S2
- **Peptide Synthesis and Purification**……S2-S3
- **Biophysical Characterization**……S3-S4
- **Fluorescence Measurements**……S4-S6
- **X-Ray Structure Determination**……S6

**CC-Di Kinetics Data**……S7-S22

**CC Basis Set Data**……S23-S35
- **Sequences**……S23
- **MALDI and AHPLC**……S23-S27
- **CD Characterization**……S27-S29
- **Fluorescence Measurements**……S30-S35

**CC-Tri and CCTet* Fluorescence Measurements**……S36

**Orthogonal CC Set Data**……S36-S49
- **Sequences**……S36
- **MALDI and AHPLC**……S36-S38
- **CD Characterization**……S38-S39
- **X-Ray Structures**……S40-S41
- **Analytical Ultracentrifugation Data**……S42
- **Fluorescence Measurements**……S43-S49

**References**……S50-S51

#### Methods

##### General

All solvents, chemicals, and reagents were purchased from commercial sources and used without further purification. Fluorenylmethoxycarbonyl(Fmoc)-α-L-amino acids, Rink amide MBHA resin for solid-phase peptide synthesis (SPPS) and N,N-dimethylformamide (DMF) were purchased from Sigma-Aldrich and Cambridge Reagents. Coupling reagents Oxyma Pure and diisopropylcarbodiimide (DIC) were purchased from Fluorochem. Morpholine, and trifluoroacetic acid (TFA) were purchased from Sigma-Aldrich; pyridine was from Thermo Fisher; triisopropylsilane (TIPS) was from Acros Organics. All other chemicals were reagent grade and purchased from Sigma-Aldrich. Peptide biophysical characterization data were recorded in phosphate buffered saline (PBS; 8.2 mM sodium phosphate dibasic, 1.8 mM potassium phosphate monobasic, 137 mM NaCl, 2.4 mM KCl), pH 7.4. Peptide characterisation data for CC-Di, CC-Tri, CC-Tet, CC-Tet*, CC-Pent2, CC-Hex2, and CC-Hept have been published previously.[1-3]

##### Peptide Synthesis

Peptides were prepared by standard Fmoc microwave-assisted solid-phase peptide synthesis using a Liberty Blue™ automatic synthesizer (CEM) with inline UV monitoring. Pre-swelled Rink amide MBHA resin (100 µmol) was added to the sample loader. Fmoc-protected amino acids were coupled to the resin via addition of 2.5 mL of 0.2 M Fmoc-amino acid (5 equiv.) in DMF, 1.0 mL of 1 M *N,N'*-diisopropylcarbodiimide in DMF (10 equiv.) and 1.0 mL of 0.5 M ethyl cyano(hydroxyamino)acetate (5 equiv.) in DMF, as recommended by CEM, to the reaction vessel. Deprotection of the Fmoc group was performed by addition of 20% (v/v) morpholine in DMF into the reaction vessel and heating the reaction vessel to 90 ºC for 1 min (125 W 30 s, 32 W 60 s).

Following the deprotection of the N-terminal Gly residue, the resin was transferred to a fritted syringe. Unlabeled peptides were acetyl capped by addition of 0.5 mL acetic anhydride and 0.25 mL of pyridine in excess DMF. The resulting mixture was rocked at RT for 20 min. Carboxy-fluorescein (5 equiv.) was coupled onto the N-terminus to generate labeled peptides using the coupling conditions described above and rocked at RT for 2 hrs. Acetyl and FAM-capped peptides were washed 4X with DMF and 4X with DCM prior to cleavage. Peptides were then cleaved from the resin by addition of 8 mL of a cleavage cocktail consisting of 2.5 % $H_2O$, 2.5% triisopropylsilane and 95% trifluoroacetic acid (TFA). The solution was rocked for 2 hours and subsequently filtered through the fritted syringe. The resin was washed two more times with TFA, and the filtered TFA solutions were combined. The TFA in the combined solution was evaporated under a stream of nitrogen. To the resulting brown oil was added chilled diethyl ether, which caused precipitation. Crude peptide was isolated using centrifugation, re-dissolved in 1:1 MeCN:$H_2O$ and lyophilized to yield a white powder.

**Peptide Purification**

Crude products of peptide syntheses were purified via RP-HPLC using a Jasco system consisting of a UV-4075 UV/Vis detector, PU-4180 HPLC Pump, LC-Net II/ADC computer-instrument interface, and co-2060 Plus HPLC column thermostat. The purification was conducted with Luna® C18 (Phenomenex) column (150 mm x 10 mm, 5 μm particle size, 100 Å pore size) using a gradient elution of 20-80%, 30-90%, or 40-100% B solvent over 30 minutes. Solvent A is 0.1% TFA in MilliQ $H_2O$, and solvent B is 0.1% TFA in HPLC-grade acetonitrile. When required, the column was heated to 50 ºC to assist peptide elution. Chromatograms were monitored at 220 and 280 nm wavelengths. The presence of the desired peptide was established via matrix-assisted laser desorption ionization-time of flight (MALDI-TOF) mass spectroscopy attained via use of a Bruker ULTRAFLEX™ instrument in positive-ion reflector mode. Peptides were spotted on a ground steel target plate using α-cyano-4-hydroxycinnamic acid dissolved in 1:1 MeCN/$H_2O$ as the matrix. Expected monoisotopic masses of the singly protonated species of each peptide are listed in Tables S2 and S5. Purity was quantified utilizing a Jasco 2000 HPLC and a Phenomenex Kinetex C18 (100 x 4.6 mm, 5 μm particle size, 100 Å pore size) column. Chromatograms were monitored at 220 and 280 nm wavelengths. The linear gradient was 20 – 100% MeCN in water (each containing 0.1% TFA) over 25 min at a flow rate of 1 mL/min.

**Determining Concentration of FAM-labeled Peptides**

The concentration of purified lyophilized peptide dissolved in MilliQ water was determined by measuring the absorbance of the sample at 280 nm using a ThermoScientific 2000 UV-Vis spectrometer. For all non-labeled peptides, the $\varepsilon_{280nm}$ = 5500 $cm^{-1}M^{-1}$ as each contains a single tryptophan. The protocol for determining the concentration of the FAM-labeled peptides follows the one described in "Exploring the Dynamic and Conformational Landscape of α-Helical Peptide Assemblies."[4] Peptide samples were diluted in 25 mM Tris pH 8, 1% SDS buffer and their absorbance measured at 495 nm. The extension coefficient used for FAM-labeled peptides in 25 mM Tris pH 8, 100 mM NaCl, 1% SDS buffer was previously measured as $\varepsilon_{495nm}$ = 89000 $cm^{-1}M^{-1}$.

**Circular Dichroism**

Circular dichroism (CD) data were collected on a JASCO J-810 spectropolarimeter fitted with a Peltier temperature controller. Peptide samples were made up as 50 or 25 μM peptide solutions in phosphate buffered saline (PBS; 8.2 mM sodium phosphate dibasic, 1.8 mM potassium phosphate monobasic, 137 mM NaCl, 2.4 mM KCl), pH 7.4. Data were collected in a 1 mm quartz cuvette between 190 and 600 nm with the instrument set as follows: band width 1 nm, data pitch 1 nm, scanning speed 100 nm/min, 1 s response time at 5 °C. Each CD spectrum was obtained by averaging 8 scans and subtracting the background signal of buffer and cuvette. For thermal response experiments, the CD signal at 222 nm wavelength was monitored over the temperature range 5 – 95 °C at a ramp rate of 60 °C per hour, with the same settings and peptide concentrations given above. The spectra were converted from ellipticities (mdeg) to mean residue ellipticities (MRE, ($deg \times cm^2 \times dmol^{-1} \times res^{-1}$)) by normalizing for the concentration of peptide bonds and the cell path length using equation 1:

$$MRE = \frac{\theta x 10^6}{cxlxn}$$

**Equation 1:** where the variable $\theta$ is the measured difference in absorbed circularly polarized light in millidegrees, $c$ is the µM concentration of the compound, $l$ is the path length of the cuvette in mm, and $n$ is the number of amide bonds in the polypeptide.

**Analytical Ultracentrifugation**

Analytical ultracentrifugation (AUC) was performed on a Beckman Optima X-LA or X-LI analytical ultracentrifuge with an An-50-Ti or An-60-Ti rotor (Beckman-Coulter). Buffer densities, viscosities and peptide and protein partial specific volumes ($\overline{v}$) were calculated using SEDNTERP (http://rasmb.org/sednterp/). For sedimentation velocity (SV) experiments, peptide samples were prepared in PBS at 75 µM peptide concentration and placed in a sedimentation velocity cell with 2-channel centerpiece and quartz windows. The samples were centrifuged at 50 krpm at 20 °C, with absorbance scans taken over a radial range of 5.8 – 7.3 cm at 5 min intervals to a total of 120 scans. Data from a single run were fitted to a continuous c(s) distribution model using SEDFIT[5] at a 95% confidence level. Residuals for sedimentation velocity experiments are shown as a bitmap in which the grayscale shade indicates the difference between the fit and raw data (residuals < -0.05 black, > 0.05 white). Good fits are uniformly grey without major dark or light streaks. Sedimentation equilibrium (SE) experiments were performed at 50 µM peptide concentration in 110 µL at 20 °C. The experiment was run in triplicate in a six-channel centerpiece. The samples were centrifuged at speeds in the range of 20 – 45 krpm and scans at each recorded speed were duplicated after equilibration for 8 hours. Data were fitted using SEDPHAT[6] to a single species model. Monte Carlo analysis was performed to give 95% confidence limits.

**Fluorescence Measurements – CC-Di Kinetics**

Kinetic traces of CC-Di exchange were measured using a Jasco FP-6500 spectrofluorometer with a Julabo F12 temperature controller set at the desired temperature. Fluorescence was measured at an excitation wavelength of 495 nm (± 2 nm) and an emission wavelength of 550 nm (± 2 nm). Time-course measurements were carried out for up to 10,000 seconds or until an end to the rise in fluorescence had been reached where possible. Stocks of labeled and unlabeled peptides were maintained at the experiment temperature in a Grant-bio thermo-shaker. 100 µL of the stock of labeled peptide was added to a quartz cuvette and allowed to equilibrate; at this point, the fluorescence measurement was started. Once the fluorescence of the labeled stock had equilibrated, an equal volume of the unlabeled peptide stock was added to the solution and mixed thoroughly in the cuvette; after this, the lid was closed, and the fluorescence time course measurement resumed. The curves were fit to either a single or second-order exponent (equation 2).

$$f(t)=A(1-\exp(k_{obs}\cdot t))$$

**Equation 2**: where t is time, A is the pre-exponential factor and $k_{obs}$ is the observed rate. Activation energy of CC-Di was calculated using Equation 3.

$$k=A\cdot exp(^{-E_A}/_{RT})$$

**Equation 2**: The Arrhenius equation, where *k* is the rate constant, A is the pre-exponential factor, $E_A$(J $mol^{-1}$) is the activation energy, R is the molar gas constant (J $mol^{-1}K^{-1}$), and T is the temperature (K).

All measurements were replicated 3 times.

Data were processed and fit to equation 2 using the code in a Jupyter notebook. This code is available on the Woolfson lab github (http://github.com/woolfson-group/CC_exchange)

**Fluorescence Measurements – Screen of Hendecad-Incorporated Peptides**

Samples of 1:10 FAM-labeled peptide assembly:unlabeled peptide assembly were prepared in PBS buffer. The mixtures were kept in the dark and incubated at 25 °C. At 1hr, 24hr, and post-annealing time points, 200 µL of sample was transferred to 1 cm quartz cuvettes and fluorescence measurements were conducted using a Jasco FP-6500 spectrofluorometer with a Julabo F12 temperature controller set to 25 °C. Samples were excited at 495 nm (bandwith = 1 nm) and emission was measured from 510-700 nm (bandwith = 3 nm) with a data pitch of 1 nm. Response time was set to 1 sec, PMT Voltage was set to 500 V, and scanning speed was set to 200 nm/min. Three 510-700 nm scans were acquired for each sample and averaged. Each measurement was performed three times.

**Fluorescence Measurements – Plate Reader**

Samples of 1:10 FAM-labeled peptide assembly:unlabeled peptide assembly were prepared in PBS buffer. The mixtures were kept in the dark and incubated at 25 °C. At 1hr, 24hr, and post-annealing time points, 100 µL of sample was transferred to black 96-well plates suitable for fluorescence measurements. Fluorescence was measured using a BMG Labtech (Aylesbury, UK) Clariostar plate reader set to 25 °C. Samples were excited at 483 (+/- 14 nm) and emission measured at 530 (+/- 30 nm) with dichroic filter 502.5. For all measurements the gain was set to 500 and the focal height to 5.7 mm. All experiments were performed 3 times.

All data were normalized to the values corresponding to the FAM-labeled peptide where the annealed FAM-labeled peptide in buffer is set to zero and the annealed homotypic exchange of the FAM-labeled peptide set to one. For example, all data acquired of samples prepared with FAM-CC-Di and unlabeled peptide were normalized against the averaged fluorescence values of annealed FAM-CC-Di in buffer, set to zero, and the averaged fluorescence values of annealed FAM-CC-Di+CC-Di, set to one.

**Peptide Crystallization**

Diffraction-quality peptide crystals were grown using a sitting- drop vapor-diffusion method. Commercially available sparse matrix screens were used (Morpheus®, JCSG-plus™, Structure Screen 1 and 2, Pact Premier™, ProPlex™; Molecular Dimensions), and the drops were dispensed using a robot (Oryx8; Douglas Instruments). For each well of an MRC 2 drop plate, 0.3 µL of peptide (8 mg/mL) and 0.3 µL of reservoir solution in parallel with 0.4 µL of the peptide and 0.2 µL of reservoir solution were mixed and the plate was incubated at 20 °C. Crystals of CC-Hex2-hen2, Ala at ***h***, grown in 0.3 M magnesium formate dihydrate and 0.1 M BIS_TRIS pH 5.5, CC-Hept-hen2, Ala at ***h***, grown in 1 M sodium citrate, pH 5.5 and 20 % w/v PEG 3000, and CC-Hept-IV-hen2 grown in 1.5 M ammonium sulfate and 0.1 M sodium acetate, pH 5.0 were looped and soaked in a reservoir containing 25% (v/v) glycerol as a cryoprotectant.

**X-Ray Data Collection and Structure determination.**

Diffraction data for the crystals were obtained at the Diamond Light Source (Didcot, UK) on beamlines I04 and I24. Data were processed using the automated pipelines: Xia2 pipelines[7], which ports data through DIALS[8] or MOSFLM[9] to POINTLESS and AIMLESS[10] as implemented in the CCP4 suite[11], or XDS to XSCALE[12]; or the AUTOPROC pipelines, which use the same integrating and data reduction software in addition to STARANISO.[13] The data acquired for the CC-Hex2-hen2, Ala at ***h***, were phased using either *ab initio* phasing using ARCIMBOLDO_LITE[14,15] to generate a model for molecular replacement using PHASER.[16] A phenylalanine version of the AlphaFold2[17] model of CC-Hept-IV-hen2 was used as the search model to phase the CC-Hept-IV-hen2 data using molecular replacement using PHASER.[16] Initial models were built from the initial solution using BUCCANEER.[18] Final structures were obtained after iterative rounds of model building with COOT[19] and refinement with REFMAC5.[20]

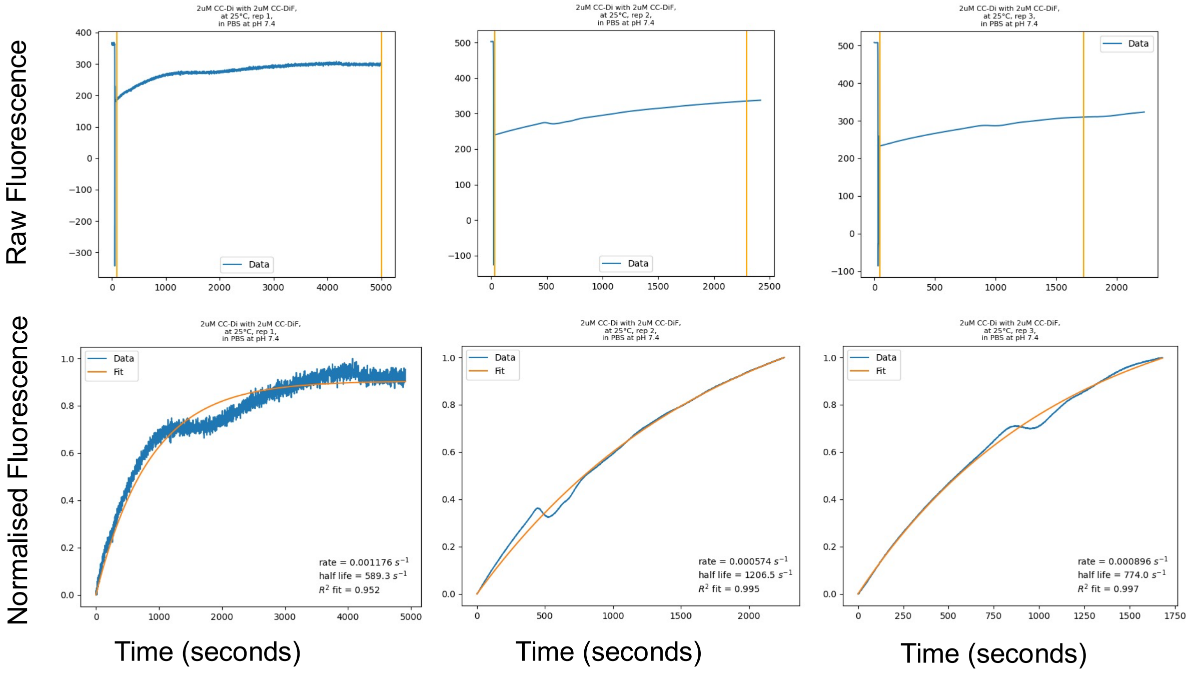


**Figure S1. Time course data and fits for the exchange of CC-Di (2 μM) and labeled CC-Di (2 μM), replicates 1-3.** (Top) Raw fluorescence (A.U.) time course data for the exchange with orange lines showing where the raw time course was trimmed for the fit to the single exponent. (Bottom) Normalized fluorescence data (A.U.) (blue) and monoexponential fits (orange).

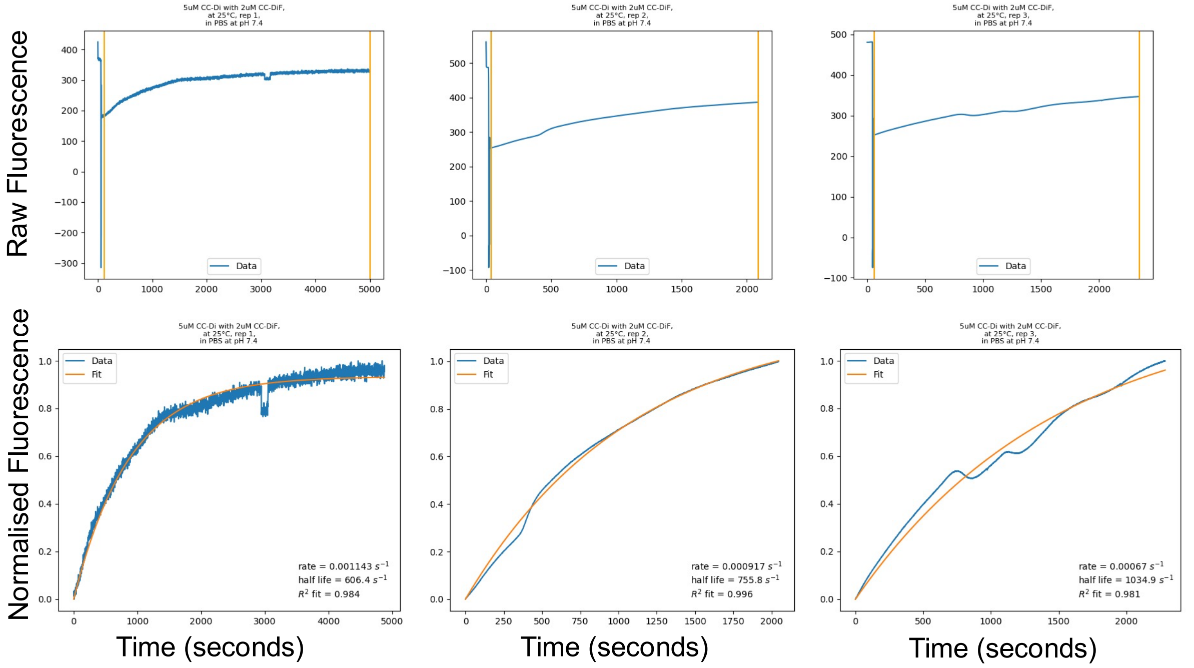


**Figure S2. Time course data and fits for the exchange of CC-Di (5 μM) and labeled CC-Di (2 μM), replicates 1-3.** (Top) Raw fluorescence (A.U.) time course data for the exchange with orange lines showing where the raw time course was trimmed for the fit to the single exponent. (Bottom) Normalized fluorescence data (A.U.) (blue) and monoexponential fits

(orange).

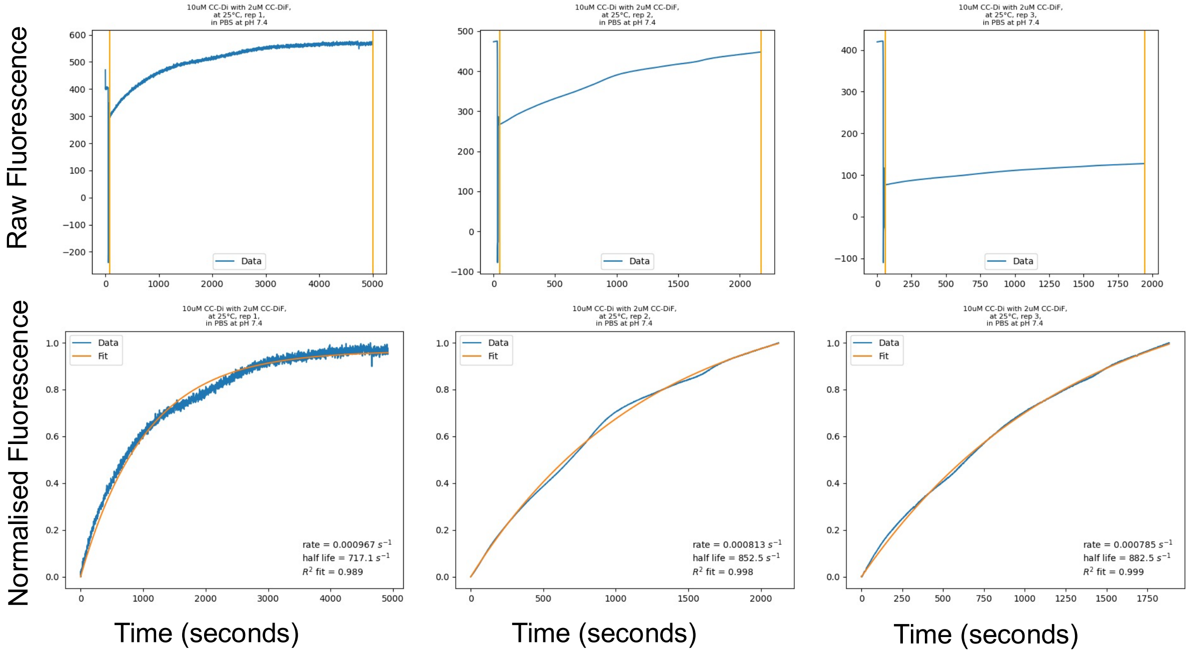


**Figure S3. Time course data and fits for the exchange of CC-Di (10 μM) and labeled CC-Di (2 μM), replicates 1-3.** (Top) Raw fluorescence (A.U.) time course data for the exchange with orange lines showing where the raw time course was trimmed for the fit to the single exponent. (Bottom) Normalized fluorescence data (A.U.) (blue) and monoexponential fits (orange).

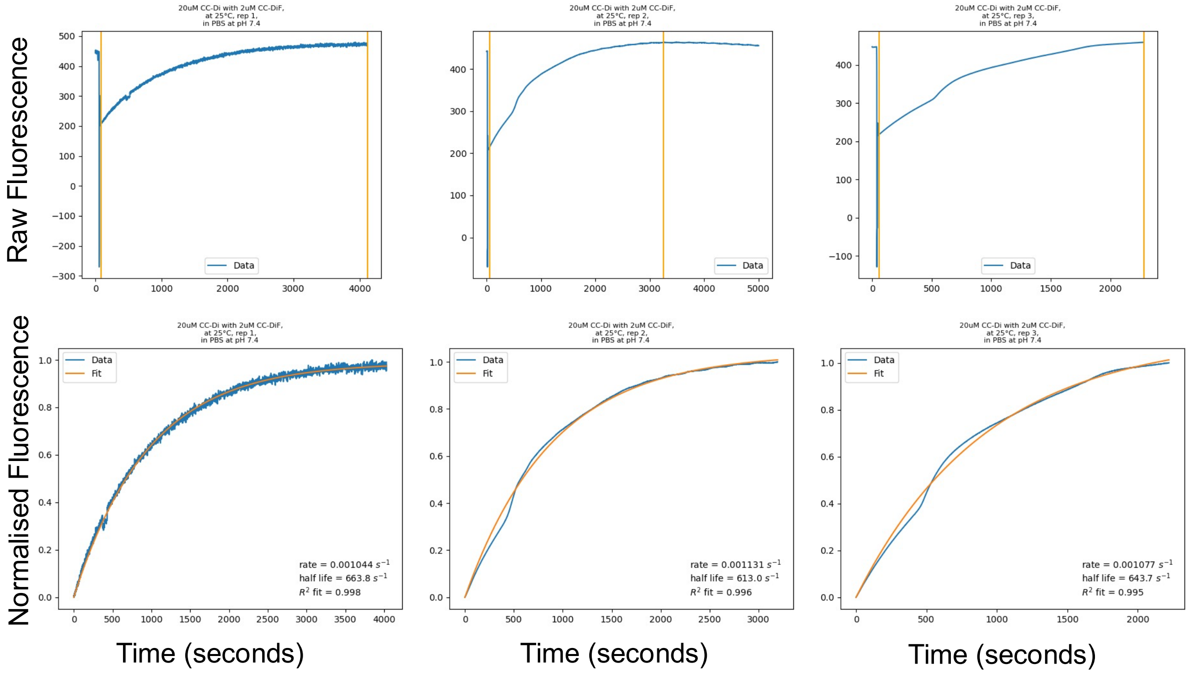


**Figure S4. Time course data and fits for the exchange of CC-Di (20 μM) and labeled CC-Di (2 μM), replicates 1-3.** (Top) Raw fluorescence (A.U.) time course data for the exchange with orange lines showing where the raw time course was trimmed for the fit to the single exponent. (Bottom) Normalized fluorescence data (A.U.) (blue) and monoexponential fits (orange).

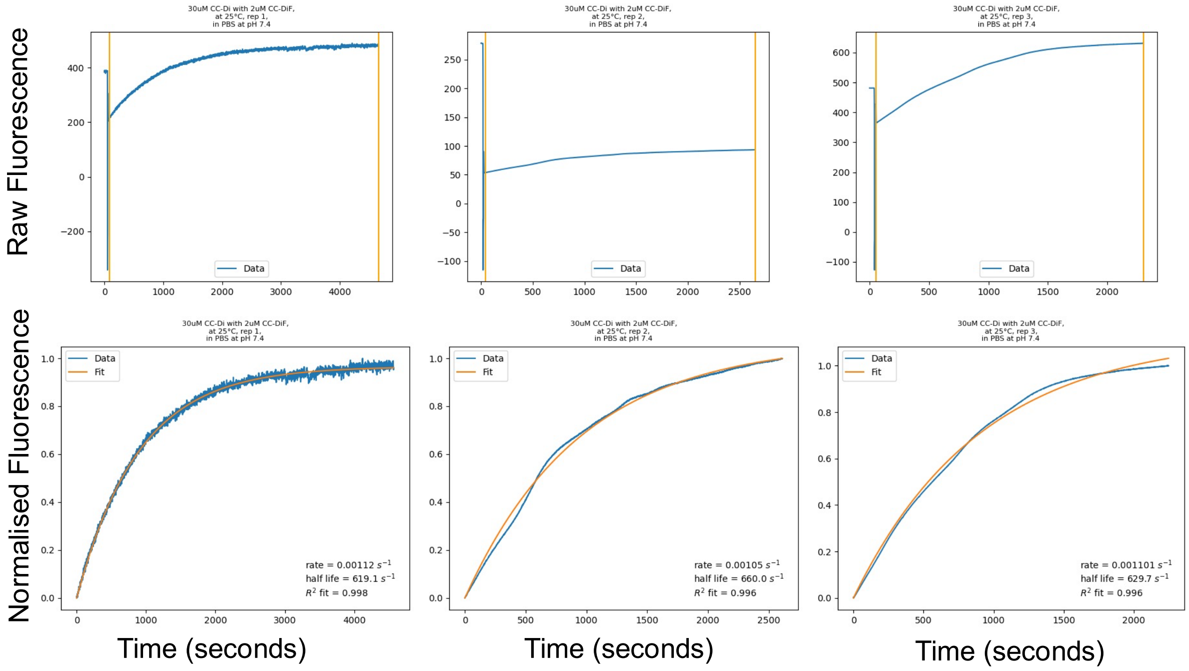


**Figure S5. Time course data and fits for the exchange of CC-Di (30 μM) and labeled CC-Di (2 μM), replicates 1-3.** (Top) Raw fluorescence (A.U.) time course data for the exchange with orange lines showing where the raw time course was trimmed for the fit to the single exponent. (Bottom) Normalized fluorescence data (A.U.) (blue) and monoexponential fits (orange).

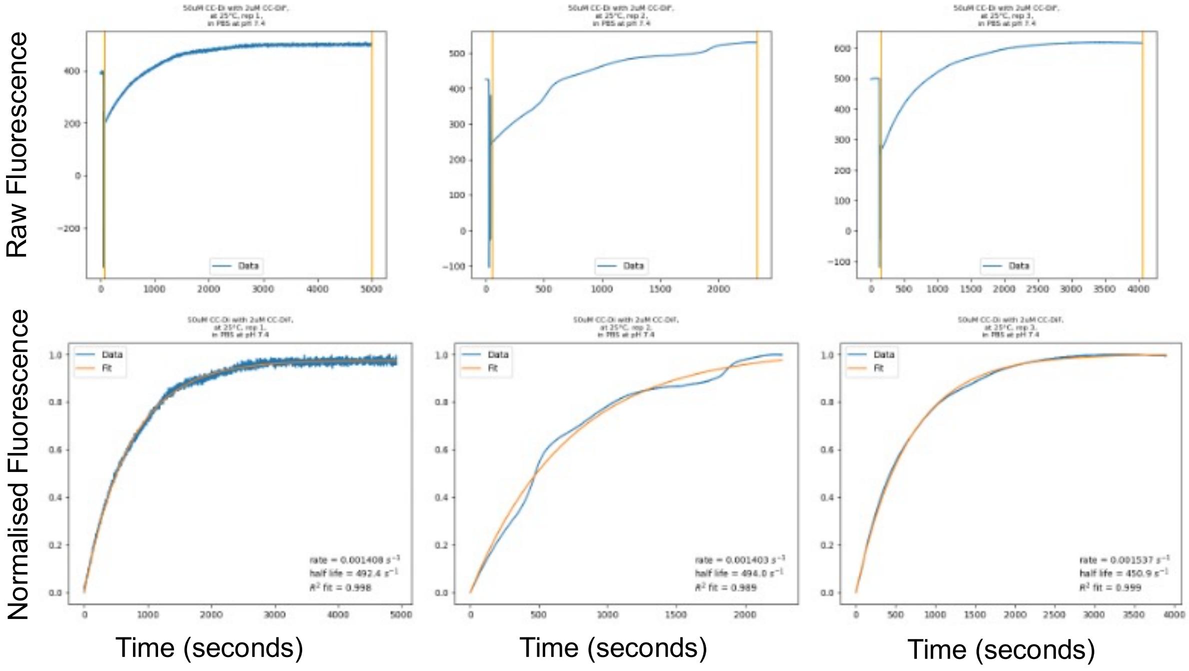


**Figure S6. Time course data and fits for the exchange of CC-Di (50 μM) and labeled CC-Di (2 μM), replicates 1-3.** (Top) Raw fluorescence (A.U.) time course data for the exchange with orange lines showing where the raw time course was trimmed for the fit to the single exponent. (Bottom) Normalized fluorescence data (A.U.) (blue) and monoexponential fits

(orange).

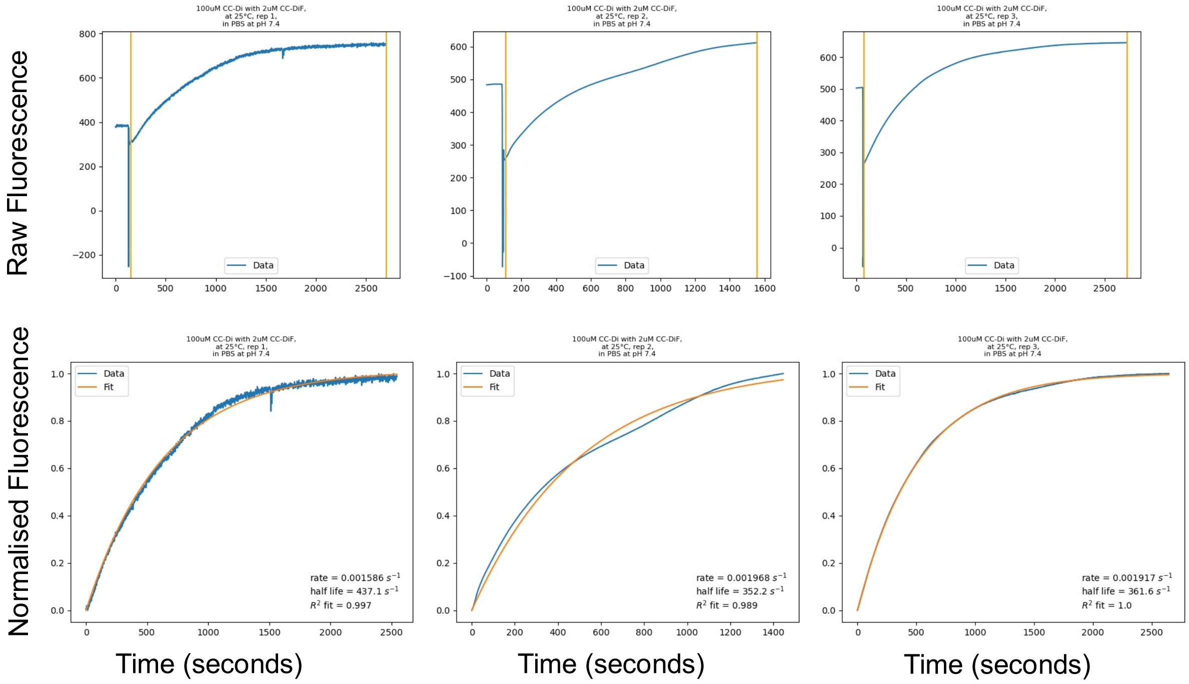


**Figure S7. Time course data and fits for the exchange of CC-Di (100 μM) and labeled CC-Di (2 μM), replicates 1-3.** (Top) Raw fluorescence (A.U.) time course data for the exchange with orange lines showing where the raw time course was trimmed for the fit to the single exponent. (Bottom) Normalized fluorescence data (A.U.) (blue) and monoexponential fits (orange).

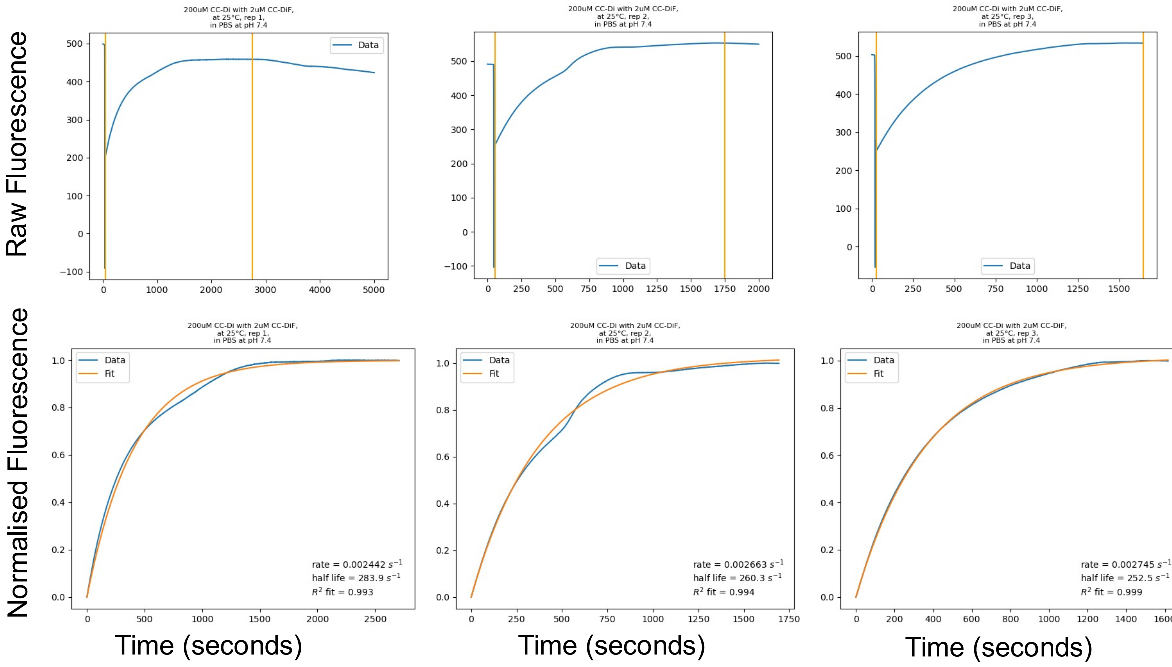


**Figure S8. Time course data and fits for the exchange of CC-Di (200 μM) and labeled CC-Di (2 μM), replicates 1-3.** (Top) Raw fluorescence (A.U.) time course data for the exchange with orange lines showing where the raw time course was trimmed for the fit to the single exponent. (Bottom) Normalized fluorescence data (A.U.) (blue) and monoexponential fits (orange).

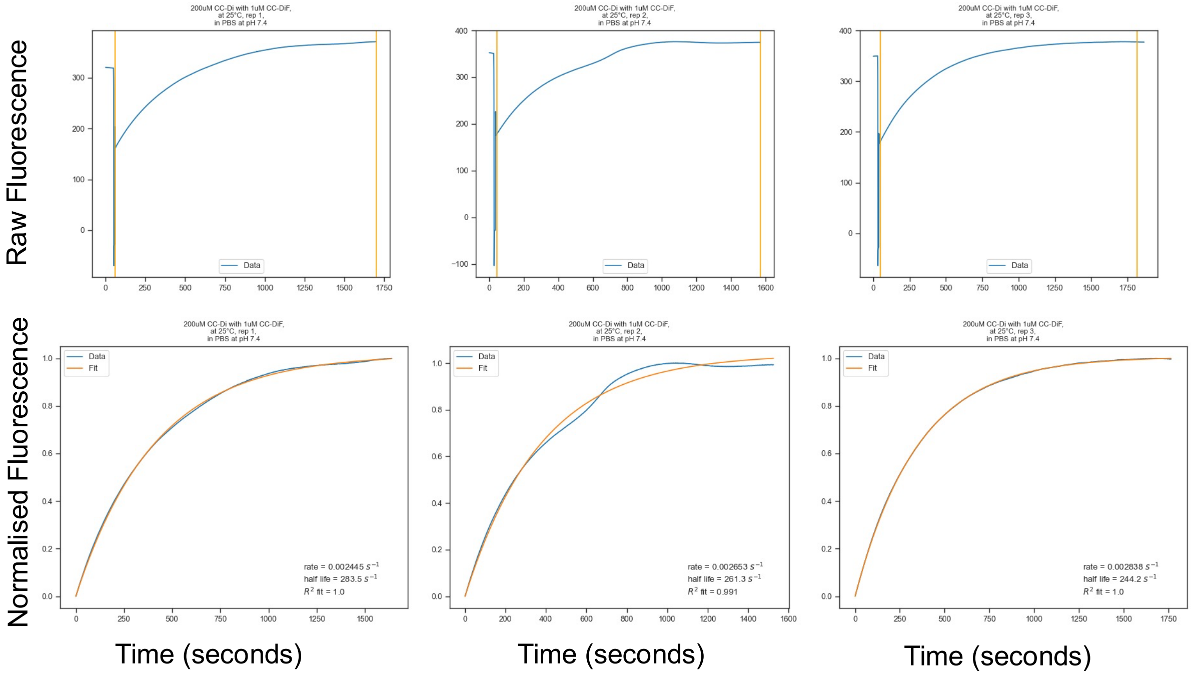


**Figure S9. Time course data and fits for the exchange of CC-Di (200 µM) and labeled CC-Di (1 µM), replicates 1-3.** (Top) Raw fluorescence (A.U.) time course data for the exchange with orange lines showing where the raw time course was trimmed for the fit to the single exponent. (Bottom) Normalized fluorescence data (A.U.) (blue) monoexponential fits (orange).

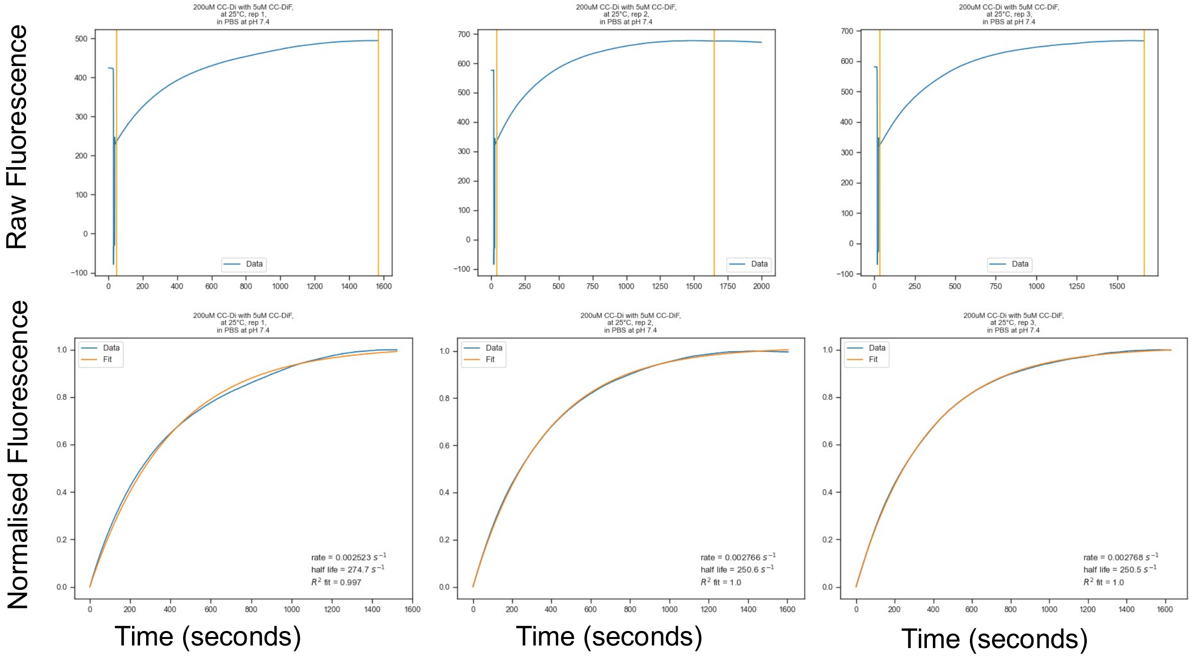


**Figure S10. Time course data and fits for the exchange of CC-Di (200 µM) and labeled CC-Di (5 µM), replicates 1-3.** (Top) Raw fluorescence (A.U.) time course data for the exchange with orange lines showing where the raw time course was trimmed for the fit to the single exponent. (Bottom) Normalized fluorescence data (A.U.) (blue) and monoexponential fits (orange).

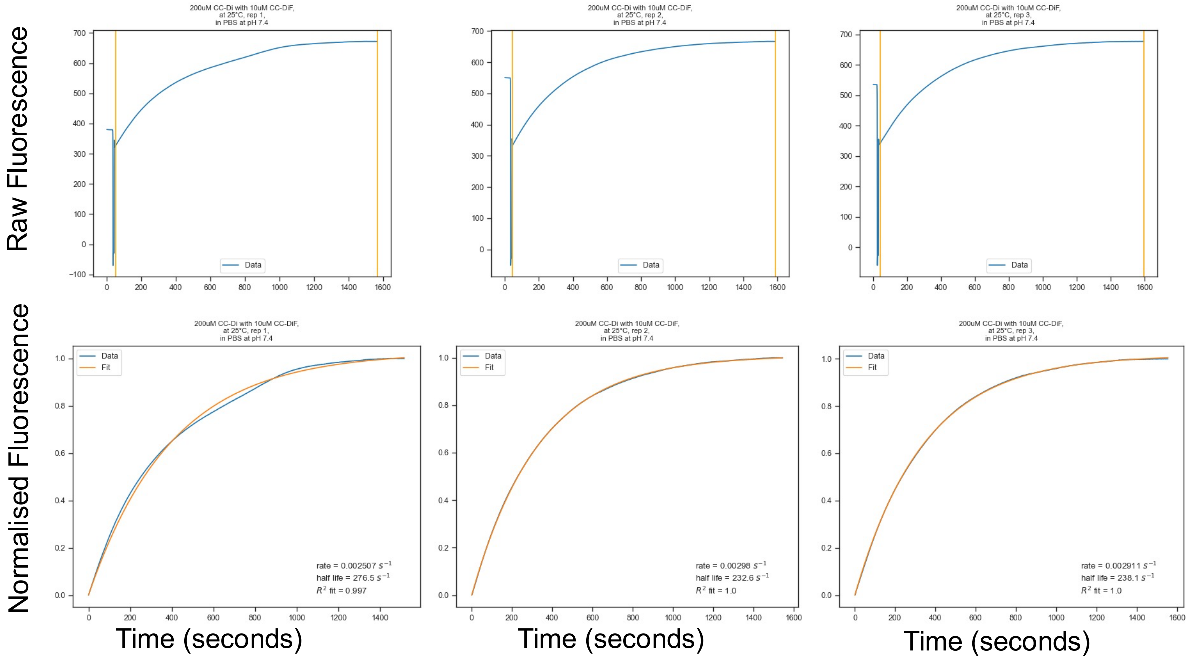


**Figure S11. Time course data and fits for the exchange of CC-Di (200 μM) and labeled CC-Di (10 μM), replicates 1-3.** (Top) Raw fluorescence (A.U.) time course data for the exchange with orange lines showing where the raw time course was trimmed for the fit to the single exponent. (Bottom) Normalized fluorescence data (A.U.) (blue) and monoexponential fits (orange).

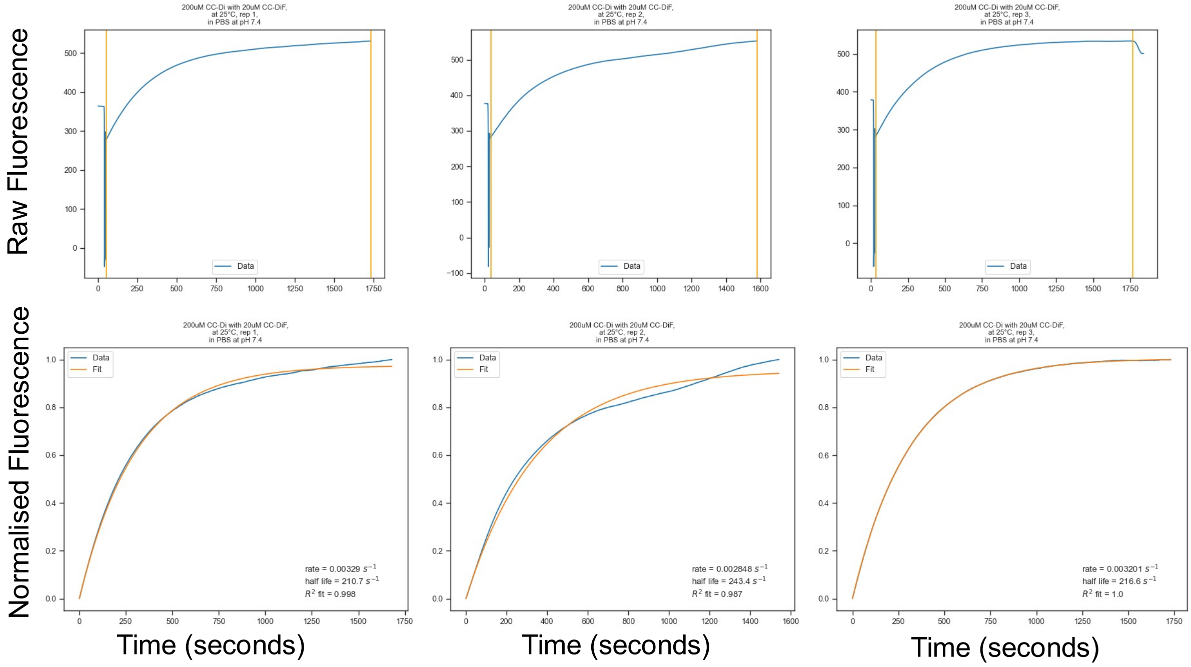


**Figure S12. Time course data and fits for the exchange of CC-Di (200 μM) and labeled CC-Di (20 μM), replicates 1-3.** (Top) Raw fluorescence (A.U.) time course data for the exchange with orange lines showing where the raw time course was trimmed for the fit to the single exponent. (Bottom) Normalized fluorescence data (A.U.) (blue) monoexponential fits

(orange).

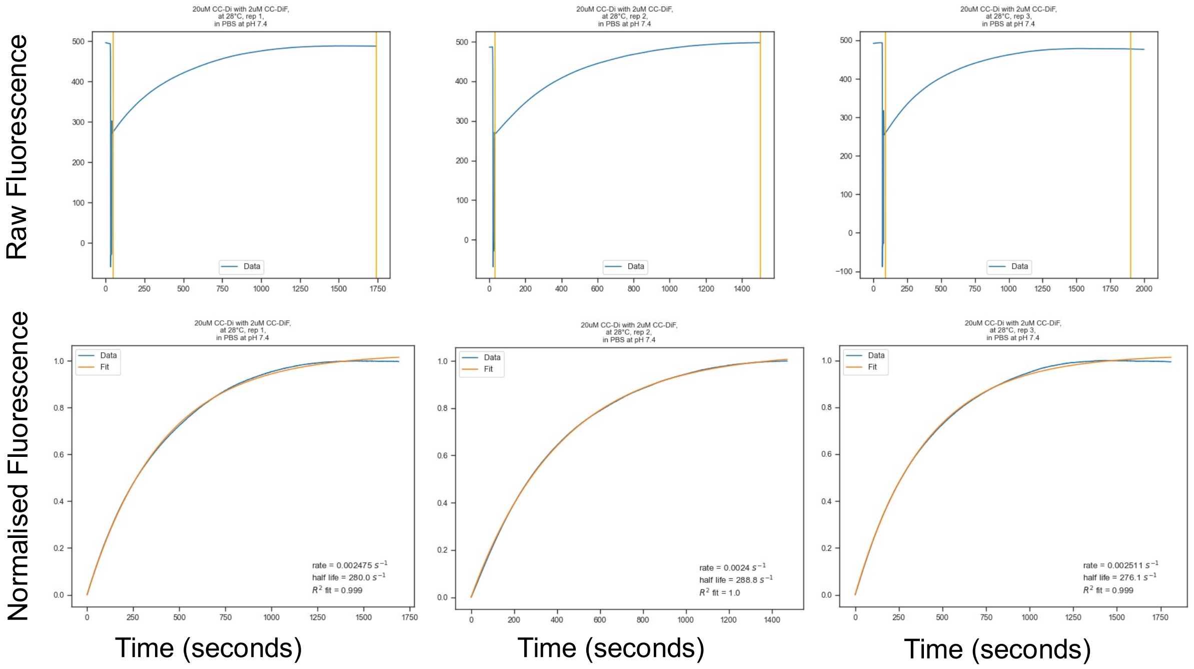


**Figure S13. Time course data and fits for the exchange of CC-Di (20 μM) and labeled CC-Di (2 μM), replicates 1-3 at 28 °C.** (Top) Raw fluorescence (A.U.) time course data for the exchange with orange lines showing where the raw time course was trimmed for the fit to the single exponent. (Bottom) Normalized fluorescence data (A.U.) (blue) and monoexponential fits (orange).

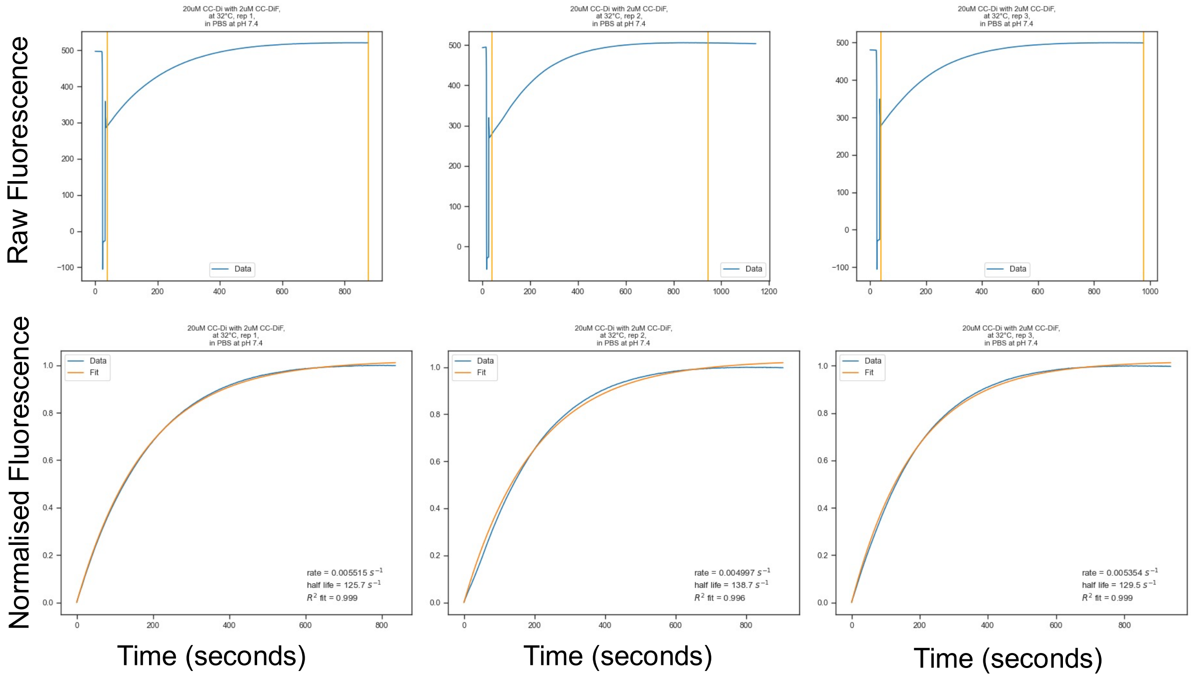


**Figure S14. Time course data and fits for the exchange of CC-Di (20 μM) and labeled CC-Di (2 μM), replicates 1-3 at 32 °C.** (Top) Raw fluorescence (A.U.) time course data for the exchange with orange lines showing where the raw time course was trimmed for the fit to the single exponent. (Bottom) Normalized fluorescence data (A.U.) (blue) and monoexponential fits (orange).

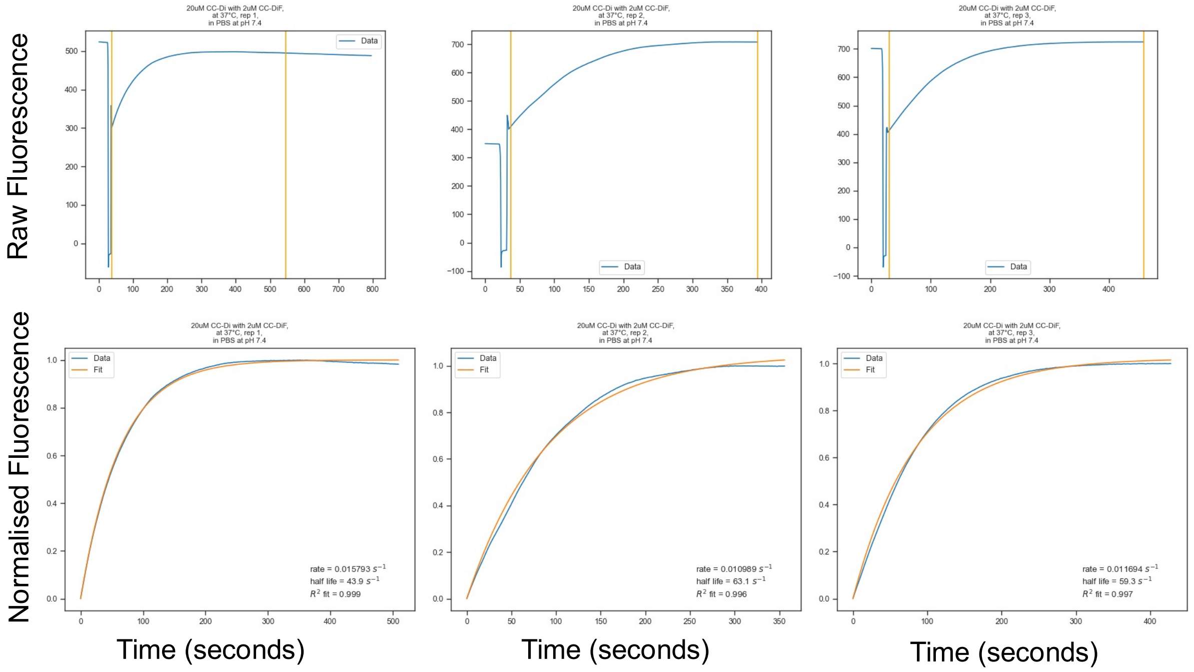


**Figure S15. Time course data and fits for the exchange of CC-Di (20 µM) and labeled CC-Di (2 µM), replicates 1-3 at 37 °C.** (Top) Raw fluorescence (A.U.) time course data for the exchange with orange lines showing where the raw time course was trimmed for the fit to the single exponent. (Bottom) Normalized fluorescence data (A.U.) (blue) and monoexponential fits (orange).

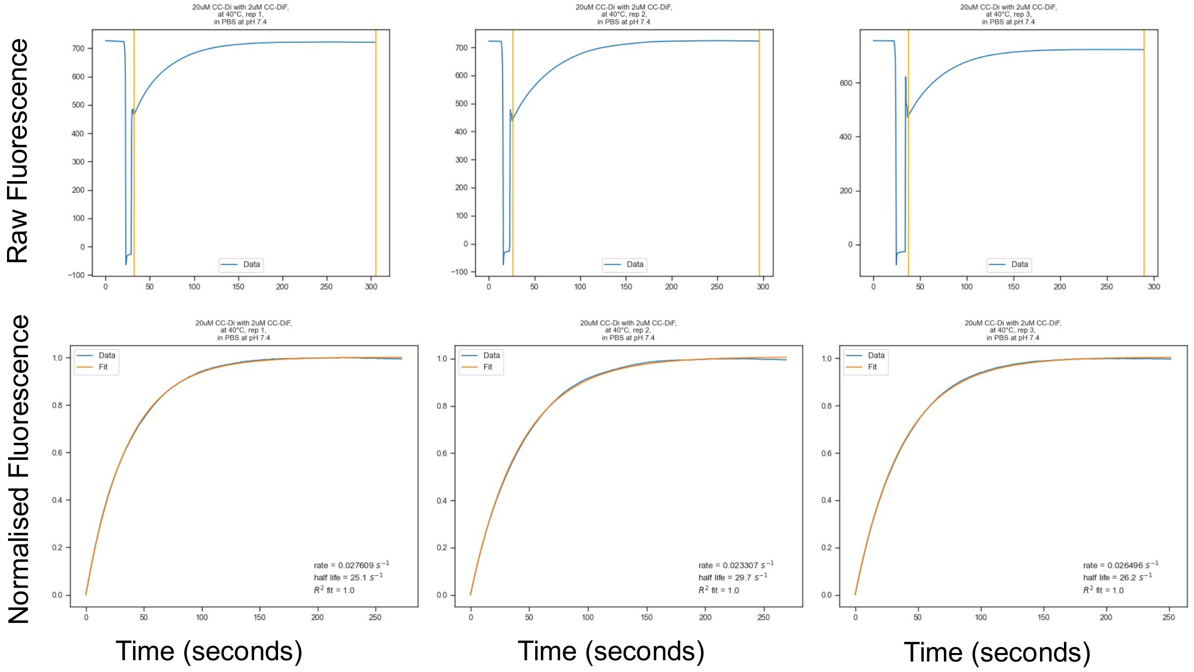


**Figure S16. Time course data and fits for the exchange of CC-Di (20 µM) and labeled CC-Di (2 µM), replicates 1-3 at 40 °C.** (Top) Raw fluorescence (A.U.) time course data for the exchange with orange lines showing where the raw time course was trimmed for the fit to the single exponent. (Bottom) Normalized fluorescence data (A.U.) (blue) monoexponential fits (orange).

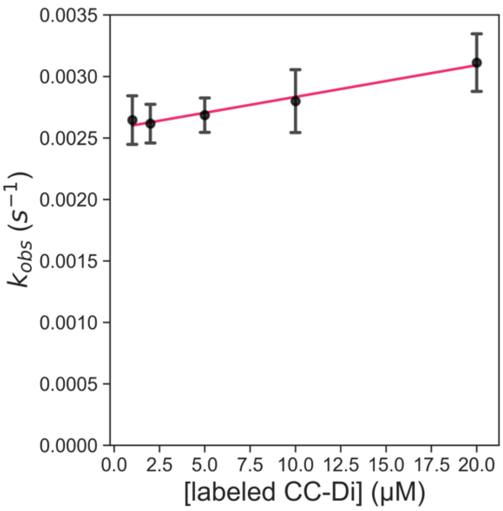


**Figure S17.** Plot of the rate ($k_{obs}$) of exchange of 200 μM unlabled CC-Di with variable concentrations of labeled CC-Di (1-20 μM**).** Data points are shown as the average of 3 independent replicates, error bars are shown to 1 standard deviation, and the line of best fit is also shown.

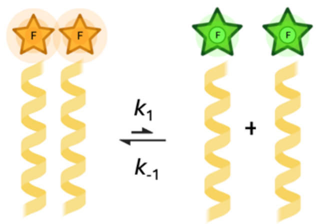


**Scheme S1.** Scheme of FAM-CC-Di dissociation.

If $[L]$ is defined as the concentration of dissociated, monomeric FAM-CC-Di and $[LL]$ is defined as the concentration of folded dimeric FAM-CC-Di at equilibrium and the overall concentration of FAM-CC-Di is 20 μM then:

At equilibrium, $K = \frac{[L]^2}{[LL]}$

$K$ = 6.67 x$10^{-11}$,[1] from which the degree of dissociation is deduced to be 0.00182 at 20 μM concentration, therefore $[L]$ = 0.0365 μM and $[LL]$ = 20.0 μM

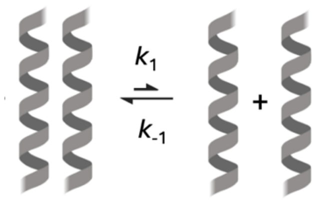


**Scheme S2.** Scheme of CC-Di dissociation.

If $[U]$ is defined as the concentration of dissociated, monomeric CC-Di and $[UU]$ is defined as the concentration of folded dimeric CC-Di at equilibrium, and the overall concentration of CC-Di is 200 μM then then:

At equilibrium, $K = \frac{[U]^2}{[UU]}$

$K$ = 6.67 x$10^{-11}$, from which the degree of dissociation is deduced to be 0.00058 at 200 μM concentration, therefore $[U]$ = 0.115 μM and $[UU]$ =200 μM

**Table S1.** Peptide concentration of dissociated monomeric ($[L]$, $[U]$) and folded dimeric ($[LL]$, $[UU]$) FAM-CC-Di and CC-Di at experimental conditions where the total concentration of unlabeled peptide was constant and the concentration of labeled peptide varied.

| **Total Concentration FAM-CC-Di (µM)** | 1 | 2 | 5 | 10 | 20 |
|---|---|---|---|---|---|
| **Total Concentration CC-Di (µM)** | 200 | 200 | 200 | 200 | 200 |
| $[L]$ (µM) | 8.13x10$^{-3}$ | 1.15x10$^{-2}$ | 1.82x10$^{-2}$ | 2.58x10$^{-2}$ | 3.65x10$^{-2}$ |
| $[LL]$ (µM) | 0.992 | 1.99 | 4.98 | 9.97 | 20.0 |
| $[U]$ (µM) | 1.15x10$^{-1}$ | 1.15x10$^{-1}$ | 1.15x10$^{-1}$ | 1.15x10$^{-1}$ | 1.15x10$^{-1}$ |
| $[UU]$ (µM) | 200 | 200 | 200 | 200 | 200 |

**Table S2.** Peptide concentration of dissociated monomeric ($[L]$, $[U]$) and folded dimeric ($[LL]$, $[UU]$) FAM-CC-Di and CC-Di at experimental conditions where the total concentration of labeled peptide was constant and the concentration of unlabeled peptide varied.

| **Total Concentration FAM-CC-Di (µM)** | 2 | 2 | 2 | 2 | 2 | 2 | 2 | 20 |
|---|---|---|---|---|---|---|---|---|
| **Total Concentration CC-Di (µM)** | 2 | 5 | 10 | 20 | 30 | 50 | 100 | 200 |
| $[L]$ (µM) | 1.15x10$^{-2}$ | 1.15x10$^{-2}$ | 1.15x10$^{-2}$ | 1.15x10$^{-2}$ | 1.15x10$^{-2}$ | 1.15x10$^{-2}$ | 1.15x10$^{-2}$ | 3.65x10$^{-2}$ |
| $[LL]$ (µM) | 1.99 | 1.99 | 1.99 | 1.99 | 1.99 | 1.99 | 1.988 | 20.0 |
| $[U]$ (µM) | 1.15x10$^{-2}$ | 1.82x10$^{-2}$ | 2.58x10$^{-2}$ | 3.65x10$^{-2}$ | 4.47x10$^{-2}$ | 5.77x10$^{-2}$ | 8.16x10$^{-2}$ | 1.15x10$^{-1}$ |
| $[UU]$ (µM) | 1.99 | 4.98 | 9.97 | 20.0 | 30.0 | 59.9 | 99.9 | 200 |

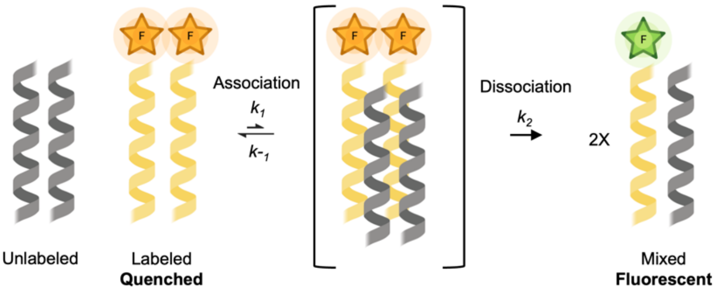


**Scheme S3.** Proposed mechanism 1 of exchange of FAM-CC-Di and CC-Di.

The rate of growth of fluorescence is the same as the rate of growth of dimers consisting of 1 copy of FAM-CC-Di and one copy of CC-Di ($LU$).

$$\text{Rate} = \frac{d[UL]}{dt} = 2k_2[UULL]$$

Applying the steady-state approximation to the tetramer:

$$\frac{d[UULL]}{dt} = k_1[UU][LL] - k_{-1}[UULL] - k_2[UULL] = 0$$

So

$$[UULL] = \frac{k_1}{k_{-1}+k_2}[UU][LL]$$

And

$$\text{Rate} = \frac{d[UL]}{dt} = \frac{2k_2k_1}{k_{-1}+k_2}[UU][LL]$$

If $[UU]$ is much greater than $[LL]$ then pseudo-first-order kinetics apply and we can write:

$$\text{Rate} = \frac{d[UL]}{dt} = k_{obs}[LL]$$

Where the pseudo-first-order rate constant $k_{\text{obs}}$ is:

$$k_{obs} = \frac{2k_2k_1}{k_{-1}+k_2}[UU]$$

Hence a plot of $k_{\text{obs}}$ vs $[UU]$ will be linear with gradient $\frac{2k_2k_1}{k_{-1}+k_2}$.

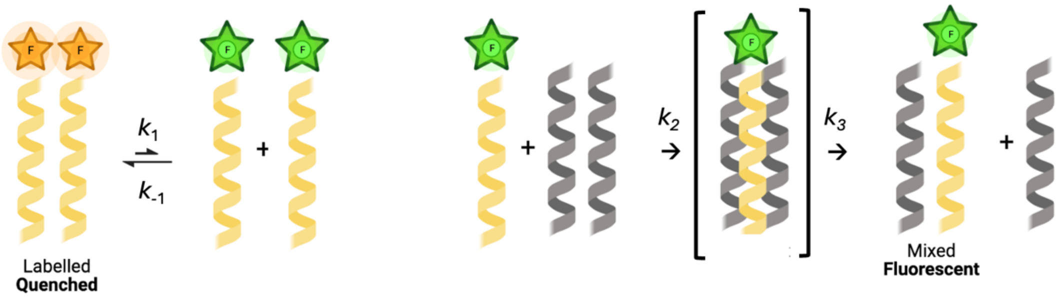


**Scheme S4.** Proposed mechanism 2 of exchange of FAM-CC-Di and CC-Di.

The rate of growth of fluorescence is the same as the rate of growth of dimers consisting of 1 copy of FAM-CC-Di and one copy of CC-Di ($LU$) if fluorescence from dissociated monomeric FAM-CC-Di and from the trimeric steady-state intermediate is negligible. This is likely accurate as the monomeric and trimeric species are unstable and only present at low concentrations at any given time.

$$\text{Rate} = \frac{d[LU]}{dt} = k_3[LUU]$$

We can also write

$$\frac{d[LUU]}{dt} = k_2[L][UU] - k_3[LUU]$$

and apply the steady state approximation to the intermediate so

$$\frac{d[LUU]}{dt} = 0$$

Hence $k_2[L][UU] = k_3[LUU]$ and

$$\text{Rate} = \frac{d[LU]}{dt} = k_2[L][UU]$$

Which is first order in $[UU]$ and $[L]$.

If we assume the dissociation of dimeric FAM-CC-Di is at equilibrium, then the equilibrium constant is $K = [L]^2/[LL]$ and hence $[L] = K^{1/2}\,[LL]^{1/2}$. Substituting to the rate expression gives:

$$\text{Rate} = \frac{d[LU]}{dt} = k_2\,K^{1/2}[LL]^{1/2}\,[UU]$$

Under experimental conditions in which $[UU] >> [LL]$, we can write:

$$\text{Rate} = \frac{d[LU]}{dt} = k_{obs}\,[LL]^{1/2}$$

Where $k_{\text{obs}} = k_2 K^{1/2}\,[UU]$. Hence a plot of $k_{\text{obs}}$ vs $[UU]$ should be linear (Figure 1C) with gradient $k_2 K^{1/2}$.

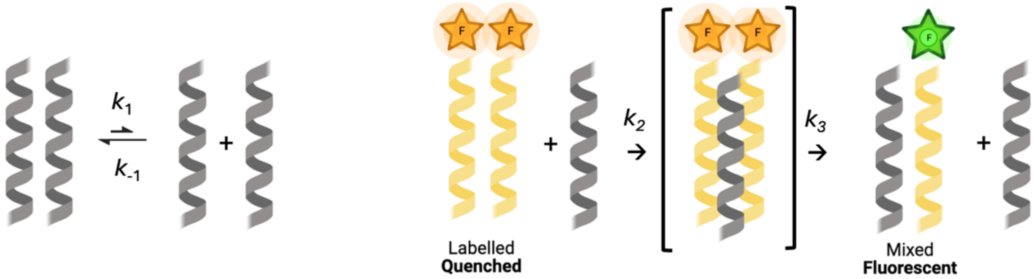


**Scheme S5.** Proposed mechanism 3 of exchange of FAM-CC-Di and CC-Di.

The rate of growth of fluorescence is the same as the rate of growth of dimers consisting of 1 copy of FAM-CC-Di and one copy of CC-Di ($LU$).

$$\text{Rate} = \frac{d[LU]}{dt} = k_3[LLU]$$

We can also write

$$\frac{d[LLU]}{dt} = k_2[LL][U] - k_3[LLU]$$

and apply the steady state approximation to the intermediate so

$$\frac{d[LLU]}{dt} = 0$$

Hence $k_2[LL][U] = k_3[LLU]$ and

$$\text{Rate} = \frac{d[LU]}{dt} = k_2[LL][U]$$

Which is first order in $[UU]$ and $[L]$.

If we assume the dissociation of dimeric CC-Di is at equilibrium, then the equilibrium constant is $K = [U]^2/[UU]$ and hence $[U] = K^{1/2}\,[UU]^{1/2}$. Substituting to the rate expression gives:

$$\text{Rate} = \frac{d[LU]}{dt} = k_2\,K^{1/2}[UU]^{1/2}\,[LL]$$

Under experimental conditions in which $[UU] >> [LL]$, we can write:

$$\text{Rate} = \frac{d[LU]}{dt} = k_{obs}\,[LL]$$

Where $k_{obs} = k_2 K^{1/2}\,[UU]^{1/2}$. Hence a plot of $k_{obs}$ vs $[UU]^{1/2}$ should be linear with gradient $k_2 K^{1/2}$.

**Table S3. Data for the exponential fits for the exchange of CC-Di**

| Temperature (°C) | [CC-Di] Unlabeled (μM) | [CC-Di] Labeled (μM) | Replicate # | Rate ($k_{obs}$, $s^{-1}$) | Halflife ($t_{1/2}$, s) | $R^2$ score of the fit |
|---|---|---|---|---|---|---|
| *Data for constant [CC-Di] Unlabelled* | | | | | | |
| 25 | 200 | 1 | 1 | 0.00244 | 284 | 0.999 |
| 25 | 200 | 1 | 2 | 0.00265 | 261 | 0.991 |
| 25 | 200 | 1 | 3 | 0.00284 | 244 | 0.999 |
| 25 | 200 | 2 | 1 | 0.00244 | 284 | 0.993 |
| 25 | 200 | 2 | 2 | 0.00266 | 260 | 0.993 |
| 25 | 200 | 2 | 3 | 0.00275 | 253 | 0.999 |
| 25 | 200 | 5 | 1 | 0.00252 | 275 | 0.997 |
| 25 | 200 | 5 | 2 | 0.00277 | 251 | 0.999 |
| 25 | 200 | 5 | 3 | 0.00277 | 250 | 0.999 |
| 25 | 200 | 10 | 1 | 0.00251 | 276 | 0.996 |
| 25 | 200 | 10 | 2 | 0.00298 | 233 | 0.999 |
| 25 | 200 | 10 | 3 | 0.00291 | 238 | 0.999 |
| 25 | 200 | 20 | 1 | 0.00329 | 211 | 0.997 |
| 25 | 200 | 20 | 2 | 0.00285 | 243 | 0.987 |
| 25 | 200 | 20 | 3 | 0.00320 | 217 | 0.999 |
| *Data for constant [CC-Di] Labelled* | | | | | | |
| 25 | 2 | 2 | 1 | 0.00118 | 589 | 0.952 |
| 25 | 2 | 2 | 2 | 0.000574 | 1207 | 0.995 |
| 25 | 2 | 2 | 3 | 0.000896 | 774 | 0.997 |
| 25 | 5 | 2 | 1 | 0.00114 | 606 | 0.984 |
| 25 | 5 | 2 | 2 | 0.000917 | 756 | 0.996 |
| 25 | 5 | 2 | 3 | 0.000670 | 1035 | 0.981 |
| 25 | 10 | 2 | 1 | 0.000967 | 717 | 0.989 |
| 25 | 10 | 2 | 2 | 0.000813 | 852 | 0.998 |
| 25 | 10 | 2 | 3 | 0.000785 | 883 | 0.999 |

| | | | | | | |
|---|---|---|---|---|---|---|
| 25 | 20 | 2 | 1 | 0.00104 | 664 | 0.998 |
| 25 | 20 | 2 | 2 | 0.00113 | 613 | 0.996 |
| 25 | 20 | 2 | 3 | 0.00108 | 644 | 0.995 |
| 25 | 30 | 2 | 1 | 0.00112 | 619 | 0.998 |
| 25 | 30 | 2 | 2 | 0.00105 | 660 | 0.996 |
| 25 | 30 | 2 | 3 | 0.00110 | 630 | 0.996 |
| 25 | 50 | 2 | 1 | 0.00141 | 492 | 0.998 |
| 25 | 50 | 2 | 2 | 0.00140 | 494 | 0.989 |
| 25 | 50 | 2 | 3 | 0.00154 | 451 | 0.999 |
| 25 | 100 | 2 | 1 | 0.00159 | 437 | 0.997 |
| 25 | 100 | 2 | 2 | 0.00197 | 352 | 0.989 |
| 25 | 100 | 2 | 3 | 0.00192 | 362 | 0.99 |
| 25 | 200 | 2 | 1 | 0.00244 | 284 | 0.993 |
| 25 | 200 | 2 | 2 | 0.00266 | 260 | 0.994 |
| 25 | 200 | 2 | 3 | 0.00275 | 253 | 0.999 |
| *Data for the Arhennius plot* | | | | | | |
| 25 | 20 | 2 | 1 | 0.00104 | 664 | 0.998 |
| 25 | 20 | 2 | 2 | 0.00113 | 613 | 0.996 |
| 25 | 20 | 2 | 3 | 0.00108 | 644 | 0.995 |
| 28 | 20 | 2 | 1 | 0.00248 | 280 | 0.999 |
| 28 | 20 | 2 | 2 | 0.00240 | 289 | 0.999 |
| 28 | 20 | 2 | 3 | 0.00251 | 276 | 0.999 |
| 32 | 20 | 2 | 1 | 0.00551 | 126 | 0.999 |
| 32 | 20 | 2 | 2 | 0.00500 | 139 | 0.996 |
| 32 | 20 | 2 | 3 | 0.00535 | 129 | 0.999 |
| 37 | 20 | 2 | 1 | 0.0158 | 43.9 | 0.999 |
| 37 | 20 | 2 | 2 | 0.0110 | 63.1 | 0.996 |
| 37 | 20 | 2 | 3 | 0.0117 | 59.3 | 0.997 |

| | | | | | | |
|---|---|---|---|---|---|---|
| 40 | 20 | 2 | 1 | 0.0276 | 25.1 | 0.999 |
| 40 | 20 | 2 | 2 | 0.0233 | 29.7 | 0.999 |
| 40 | 20 | 2 | 3 | 0.0265 | 26.2 | 0.999 |

**Table S4. Data for the fit to the Arrhenius equation and subsequent activation energy calculation.**

| **Temperature (°C)** | **Rate ($k_{obs}$, $s^{-1}$) (average of 3 replicates)** | **1/T ($K^{-1}$)** | **ln($k_{obs}$)** |
|---|---|---|---|
| 25 | 0.00108 | 0.00335 | -6.83 |
| 28 | 0.00246 | 0.00332 | -6.01 |
| 32 | 0.00529 | 0.00328 | -5.24 |
| 37 | 0.0128 | 0.00322 | -4.36 |
| 40 | 0.0258 | 0.00319 | -3.66 |
| Slope (Ea/R) | -19036 | | |
| Intercept (lnA) | 57.105 | | |
| R (molar gas constant) (J $K^{-1}$ $mol^{-1}$) | 8.3144598 | | |
| Activation Energy (J $mol^{-1}$) | 158274 | | |
| Activation Energy (kJ $mol^{-1}$) | 158.274 | | |
| Activation Energy (kcal $mol^{-1}$) | 37.8 | | |
| $R^2$ of linear regression | 0.996 | | |

**Table S5. CC Basis Set Sequences**

| ***g*-Register CC** | ***gabcdef gabcdef gabcdef gabcdef*** | **Monoistopic Mass** |
|---|---|---|
| CC-Di | Ac-G EIAALKQ EIAALKK ENAALKW EIAALKQ G-CONH2 | 3246.9 |
| FAM-CC-Di | FAM-GGGG EIAALKQ EIAALKK ENAALKW EIAALKQ G-CONH2 | 3737.0 |
| CC-Tri | Ac-G EIAAIKQ EIAAIKK EIAAIKW EIAAIKQ G-CONH2 | 3245.9 |
| FAM-CC Tri | FAM-GGGG EIAAIKQ EIAAIKK EIAAIKW EIAAIKQ G-CONH2 | 3735.0 |
| CC-Tet | Ac-G ELAAIKQ ELAAIKK ELAAIKW ELAAIKQ G-CONH2 | 3245.9 |
| FAM-CC-Tet | FAM-GGGG ELAAIKQ ELAAIKK ELAAIKW ELAAIKQ G-CONH2 | 3735.0 |
| ***c*-Register CC** | ***cdefgab cdefgab cdefgab cdefgab*** | |
| CC-Pent2 | Ac-G EIAQTLK EIAKTLK EIAWTLK EIAQTLK G-CONH2 | 3365.9 |
| FAM-CC-Pent2 | FAM-GGGG EIAQTLK EIAKTLK EIAWTLK EIAQTLK G-CONH2 | 3855.0 |
| CC-Hex2 | Ac-G EIAQSLK EIAKSLK EIAWSLK EIAQSLK G-CONH2 | 3309.9 |
| FAM-CC-Hex2 | FAM-GGGG EIAQSLK EIAKSLK EIAWSLK EIAQSLK G-CONH2 | 3800.0 |
| CC-Hept | Ac-G EIAQALK EIAKALK EIAWALK EIAQALK G-CONH2 | 3245.9 |
| FAM-CC-Hept | FAM-GGGG EIAQALK EIAKALK EIAWALK EIAQALK G-CONH2 | 3735.0 |

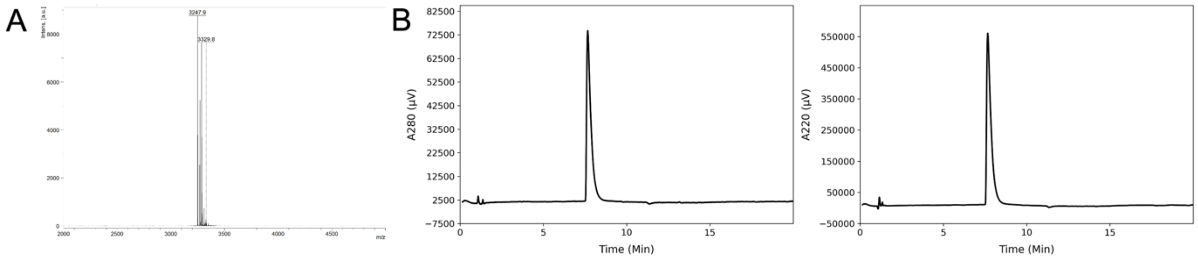


**Figure S18. CC-Di MALDI and AHPLC data.** (A) MALDI-TOF MS analysis of purified CC-Di. Monoisotopic [M+H] peaks were observed at 3247.9; calculated m/z is 3246.9. (B) Analytical HPLC trace of purified CC-Di. Chromatograms were monitored at 280 (left) and 220 (right) nm wavelengths.

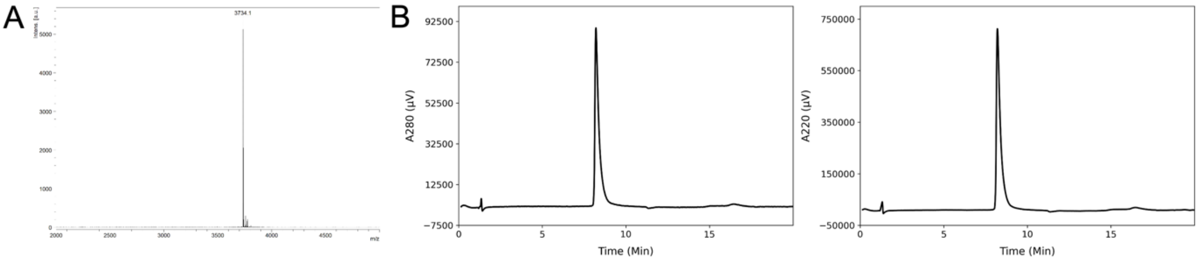


**Figure S19. FAM-CC-Di MALDI and AHPLC data.** (A) MALDI-TOF MS analysis of purified FAM-CC-Di. Monoisotopic [M+H] peaks were observed at 3734.1; calculated m/z is 3737.0. (B) Analytical HPLC trace of purified FAM-CC-Di. Chromatograms were monitored at 280 (left) and 220 (right) nm wavelengths.

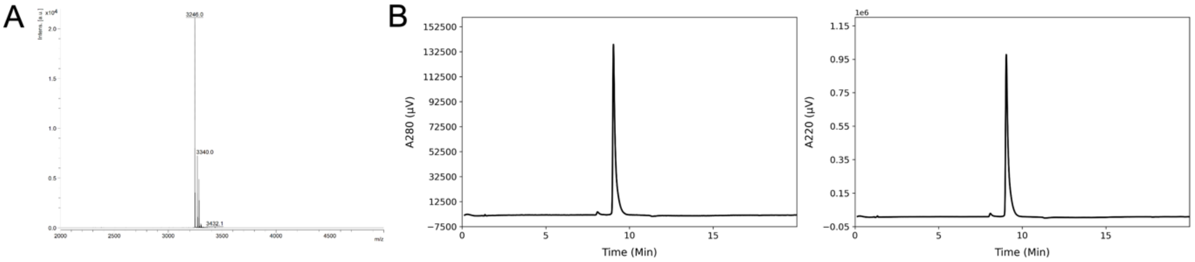


**Figure S20. CC-Tri MALDI and AHPLC data.** (A) MALDI-TOF MS analysis of purified CC-Tri. Monoisotopic [M+H] peaks were observed at 3246.0; calculated m/z is 3245.9. (B)

Analytical HPLC trace of purified CC-Tri. Chromatograms were monitored at 280 (left) and 220 (right) nm wavelengths

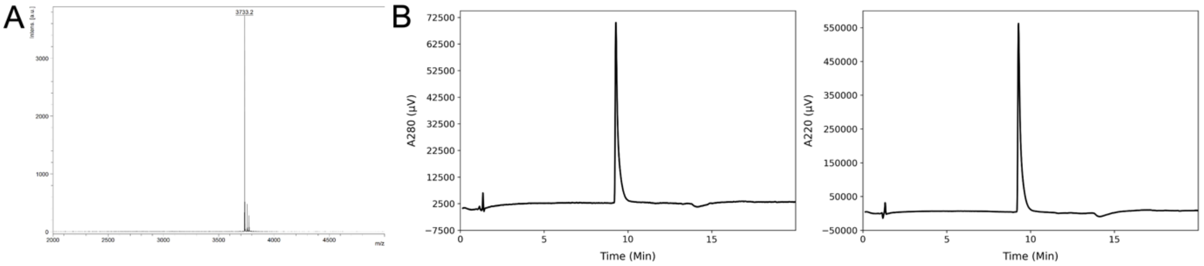


**Figure S21. FAM-CC-Tri MALDI and AHPLC data.** (A) MALDI-TOF MS analysis of purified FAM-CC-Tri. Monoisotopic [M+H] peaks were observed at 3733.2; calculated m/z is 3735.0. (B) Analytical HPLC trace of purified FAM-CC-Tri. Chromatograms were monitored at 280 (left) and 220 (right) nm wavelengths.

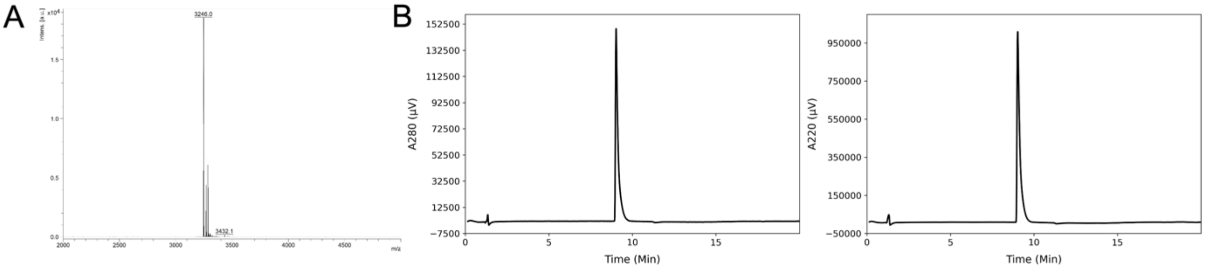


**Figure 22. CC-Tet MALDI and AHPLC data.** (A) MALDI-TOF MS analysis of purified CC-Tet. Monoisotopic [M+H] peaks were observed at 3246.0; calculated m/z is 3245.9. (B) Analytical HPLC trace of purified CC-Tet. Chromatograms were monitored at 280 (left) and 220 (right) nm wavelengths.

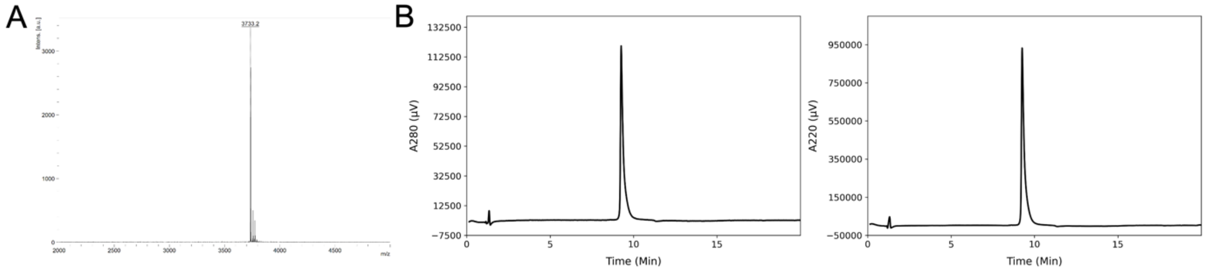


**Figure S23. FAM-CC-Tet MALDI and AHPLC data.** (A) MALDI-TOF MS analysis of purified FAM-CC-Tet. Monoisotopic [M+H] peaks were observed at 3733.2; calculated m/z is 3735.0. (B) Analytical HPLC trace of purified FAM-CC-Tet. Chromatograms were monitored at 280 (left) and 220 (right) nm wavelengths.

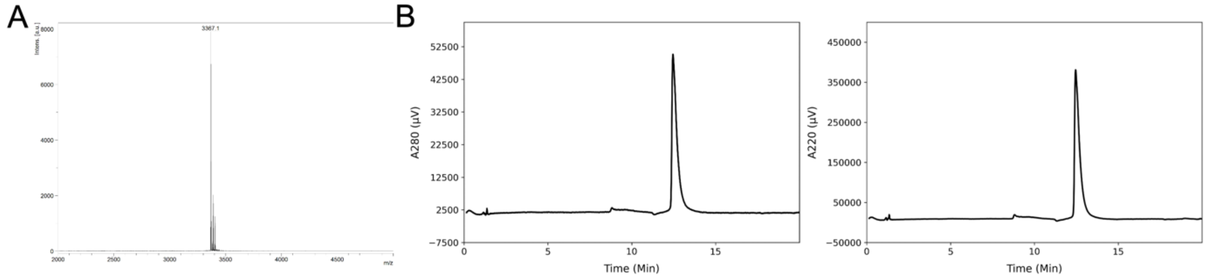


**Figure S24. CC-Pent2 MALDI and AHPLC data.** (A) MALDI-TOF MS analysis of purified CC-Pent2. Monoisotopic [M+H] peaks were observed at 3367.1; calculated m/z is 3365.9. (B)

Analytical HPLC trace of purified CC-Pent2. Chromatograms were monitored at 280 (left) and 220 (right) nm wavelengths.

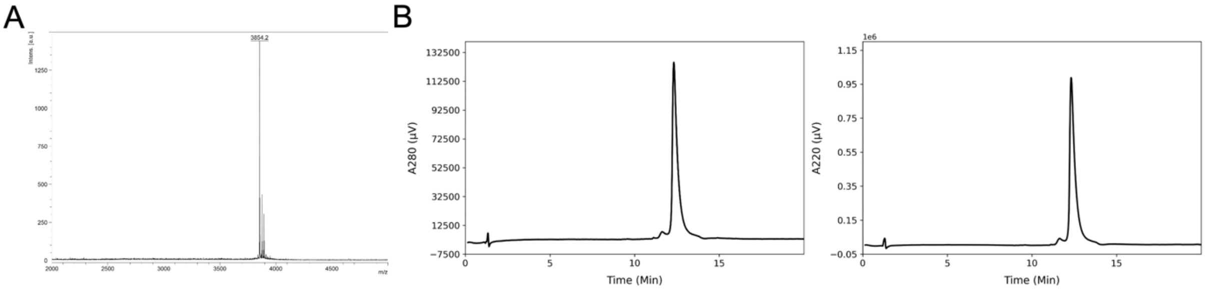


**Figure S25. FAM-CC-Pent2 MALDI and AHPLC data.** (A) MALDI-TOF MS analysis of purified FAM-CC-Pent2. Monoisotopic [M+H] peaks were observed at 3854.2; calculated m/z is 3855.0. (B) Analytical HPLC trace of purified FAM-CC-Pent2. Chromatograms were monitored at 280 (left) and 220 (right) nm wavelengths.

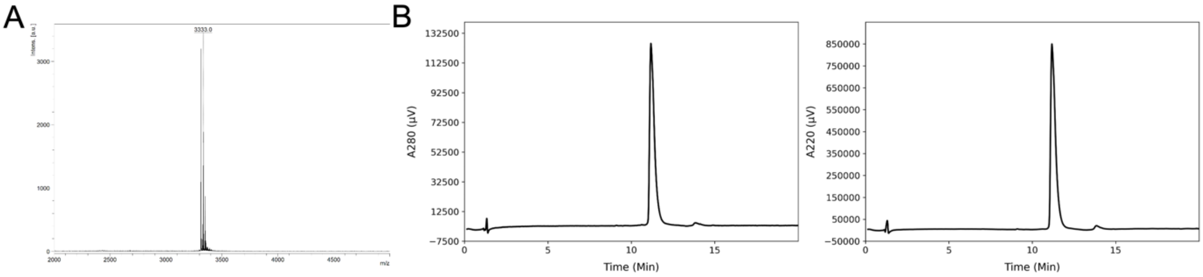


**Figure S26. CC-Hex2 MALDI and AHPLC data.** (A) MALDI-TOF MS analysis of purified CC-Hex2. Sodium adduct [M+Na] peaks were observed at 3333.0; calculated m/z is 3332.9. (B) Analytical HPLC trace of purified CC-Hex2. Chromatograms were monitored at 280 (left) and 220 (right) nm wavelengths.

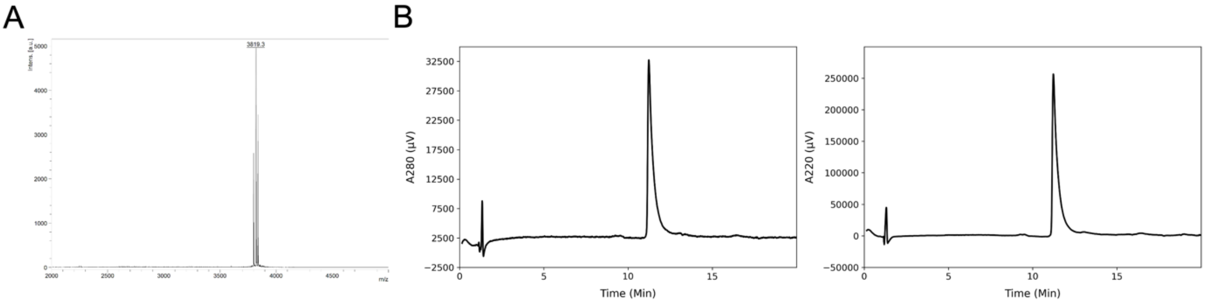


**Figure S27. FAM-CC-Hex2 MALDI and AHPLC data.** (A) MALDI-TOF MS analysis of purified CC-Hex2. Sodium adduct [M+Na] peaks were observed at 3819.3; calculated m/z is 3823.0. (B) Analytical HPLC trace of purified CC-Hex2. Chromatograms were monitored at 280 (left) and 220 (right) nm wavelengths.

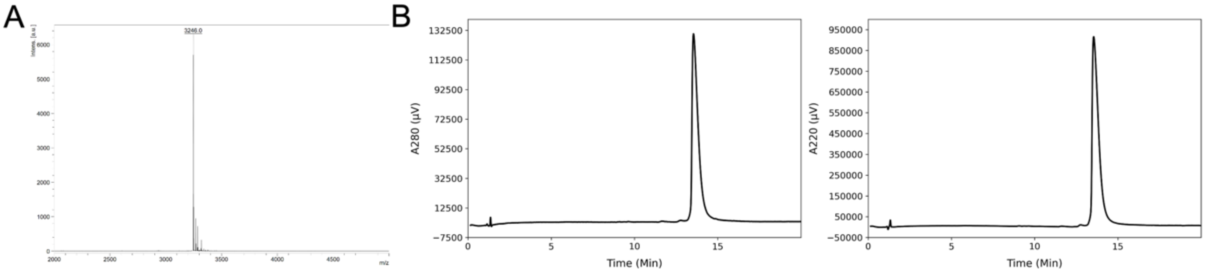


**Figure S28. CC-Hept MALDI and AHPLC data.** (A) MALDI-TOF MS analysis of purified CC-Hept. Monoisotopic [M+H] peaks were observed at 3246.0; calculated m/z is 3245.9. (B)

Analytical HPLC trace of purified CC-Hept. Chromatograms were monitored at 280 (left) and 220 (right) nm wavelengths.

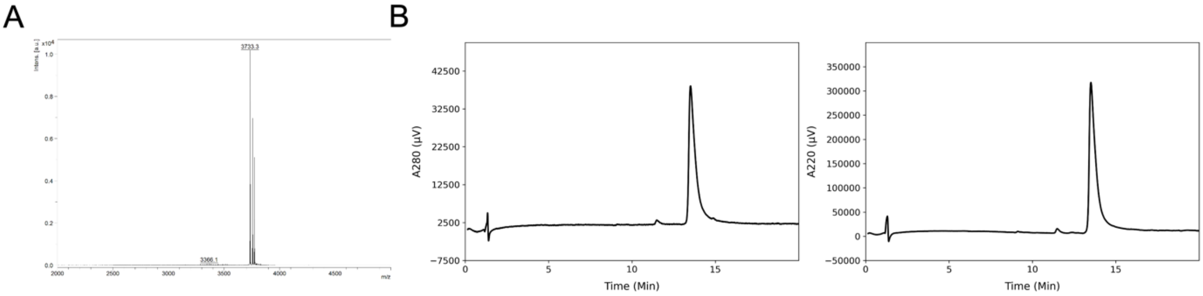


**Figure S29. FAM-CC-Hept MALDI and AHPLC data.** (A) MALDI-TOF MS analysis of purified FAM-CC-Hept. Monoisotopic [M+H] peaks were observed at 3733.3; calculated m/z is 3735.0. (B) Analytical HPLC trace of purified FAM-CC-Hept. Chromatograms were monitored at 280 (left) and 220 (right) nm wavelengths.

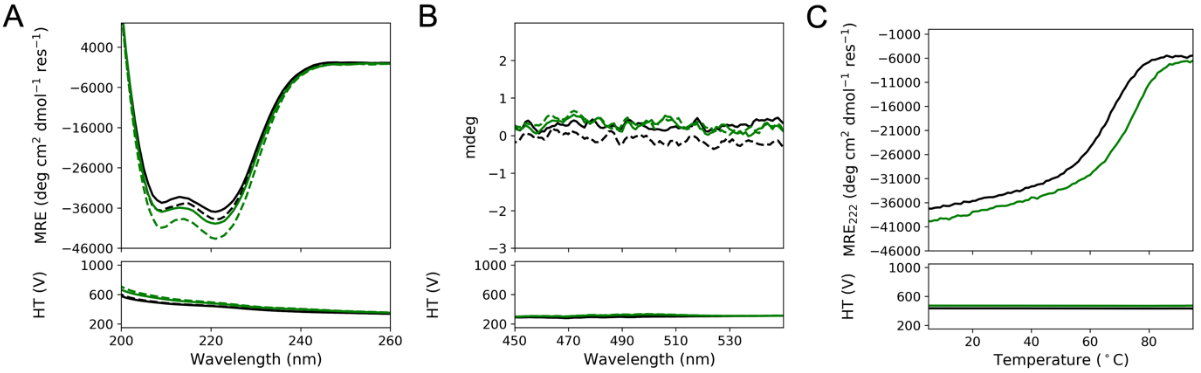


**Figure S30. CD Characterization of CC-Di and FAM-CC-Di.** CD spectra of 50 µM CC-Di (black) and FAM-CC-Di (green) in PBS at 5 °C are shown in (A) and (B). Samples were analyzed prior to melting (solid lines) and post-melting (dotted lines). (A) shows the spectra between 260-200 nm and (B) shows the spectra between 550-450 nm. (C) shows the thermal melt of the two peptides from 5-95 °C. There is not a noticeable difference in the CD characterization of the unlabeled and labeled variants of CC-Di.

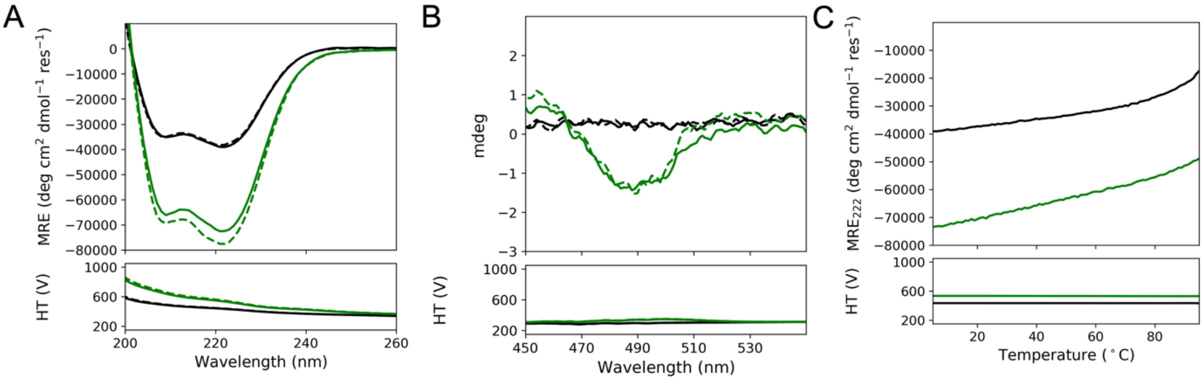


**Figure S31. CD Characterization of CC-Tri and FAM-CC-Tri.** CD spectra of 50 µM CC-Tri (black) and FAM-CC-Tri (green) in PBS at 5 °C are shown in (A) and (B). Samples were analyzed prior to melting (solid lines) and post-melting (dotted lines). (A) shows the spectra between 260-200 nm and (B) shows the spectra between 550-450 nm. (C) shows the thermal melt of the two peptides from 5-9 5°C. The labeled peptide shows an increase in CD signal (A) as well as evidence of the Cotton effect (B). However, the thermal melts are consistent between the samples, indicative that the FAM-labeled peptide has the same thermodynamic properties as the unlabeled peptide.

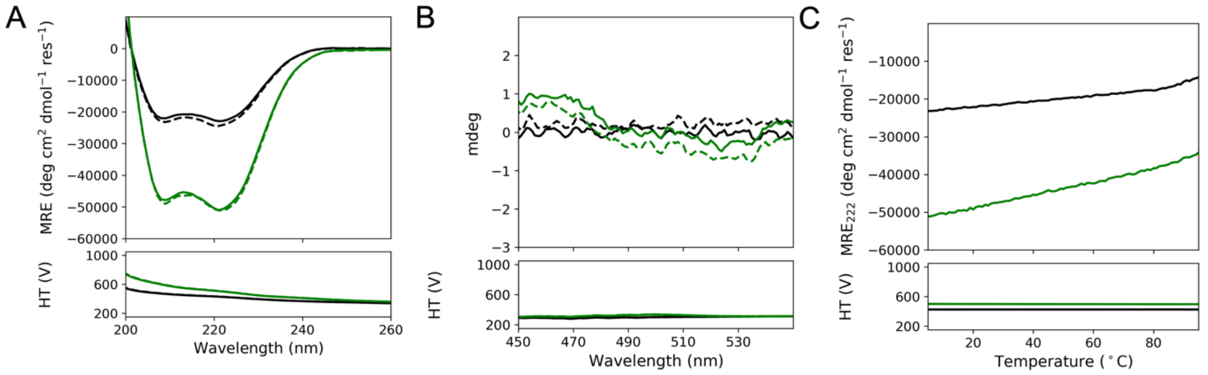


**Figure S32. CD Characterization of CC-Tet and FAM-CC-Tet.** CD spectra of 50 µM CC-Tet (black) and FAM-CC-Tet (green) in PBS at 5 °C are shown in (A) and (B). Samples were analyzed prior to melting (solid lines) and post-melting (dotted lines). (A) shows the spectra between 260-200 nm and (B) shows the spectra between 550-450 nm. (C) shows the thermal melt of the two peptides from 5-95 °C. The labeled peptide shows an increase in CD signal (A). However, the thermal melts are consistent between the samples, indicative that the FAM-labeled peptide has the same thermodynamic properties as the unlabeled peptide.

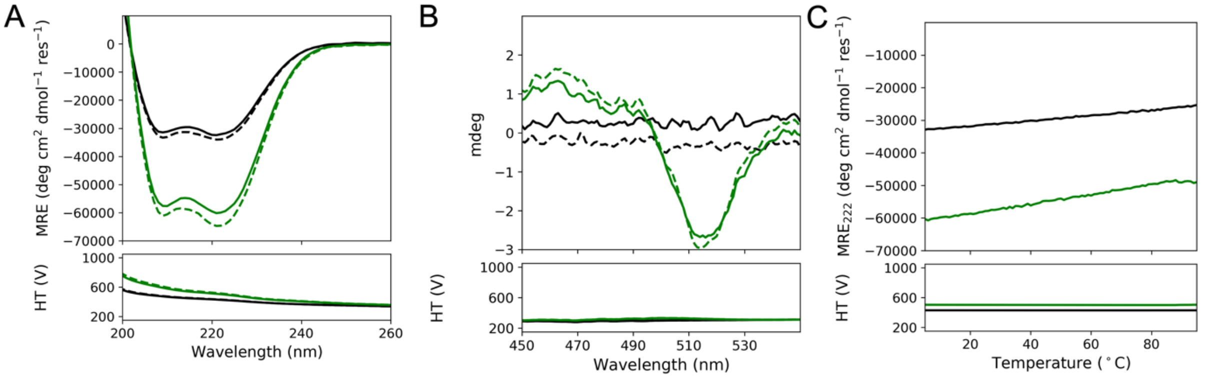


**Figure S33. CD Characterization of CC-Pent2 and FAM-CC-Pent2.** CD spectra of 50 µM CC-Pent2 (black) and FAM-CC-Pent2 (green) in PBS at 5 °C are shown in (A) and (B). Samples were analyzed prior to melting (solid lines) and post-melting (dotted lines). (A) shows the spectra between 260-200 nm and (B) shows the spectra between 550-450 nm. (C) shows the thermal melt of the two peptides from 5-95 °C. The labeled peptide shows an increase in CD signal (A) as well as evidence of Cotton effect (B). However, the thermal melts are consistent between the samples, indicative that the FAM-labeled peptide has the same thermodynamic properties as the unlabeled peptide.

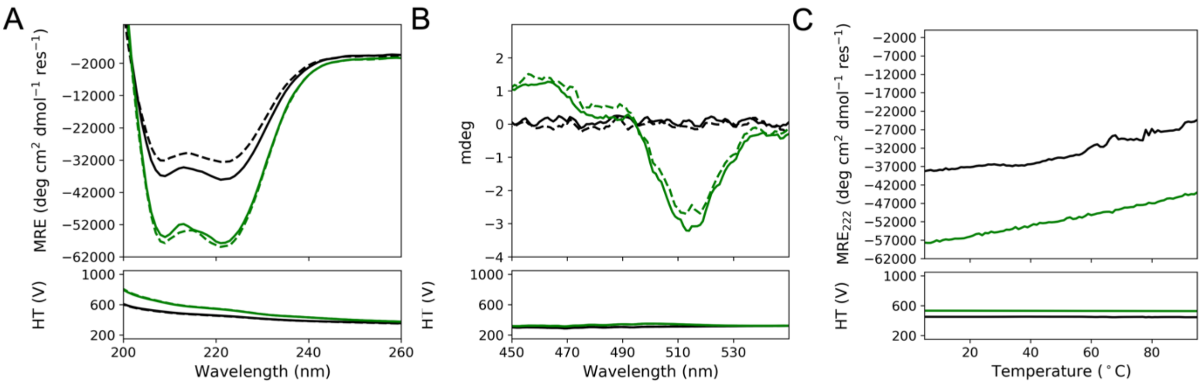


**Figure S34. CD Characterization of CC-Hex2 and FAM-CC-Hex2.** CD spectra of 50 µM CC-Hex2 (black) and FAM-CC-Hex2 (green) in PBS at 5 °C are shown in (A) and (B). Samples were analyzed prior to melting (solid lines) and post-melting (dotted lines). (A) shows the spectra between 260-200 nm and (B) shows the spectra between 550-450 nm. (C) shows the thermal melt of the two peptides from 5-95 °C. The labeled peptide shows an increase in CD signal (A) as well as evidence of Cotton effect (B). However, the thermal melts are consistent between the samples, indicative that the FAM-labeled peptide has the same thermodynamic properties as the unlabeled peptide.

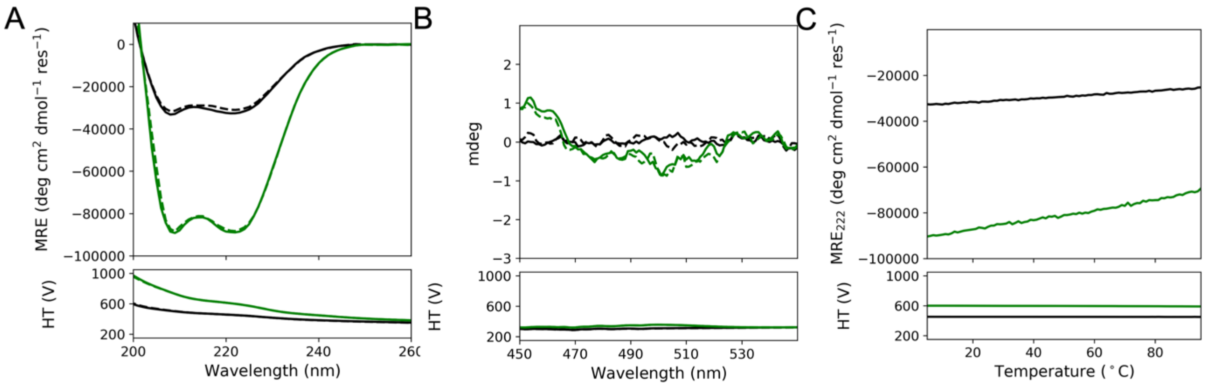


**Figure S35. CD Characterization of CC-Hept and FAM-CC-Hept.** CD spectra of 50 µM CC-Hept (black) and FAM-CC-Hept (green) in PBS at 5 °C are shown in (A) and (B). Samples were analyzed prior to melting (solid lines) and post-melting (dotted lines). (A) shows the spectra between 260-200 nm and (B) shows the spectra between 550-450 nm. (C) shows the thermal melt of the two peptides from 5-95 °C. The labeled peptide shows an increase in CD signal (A). However, the thermal melts are consistent between the samples, indicative that the FAM-labeled peptide has the same thermodynamic properties as the unlabeled peptide.

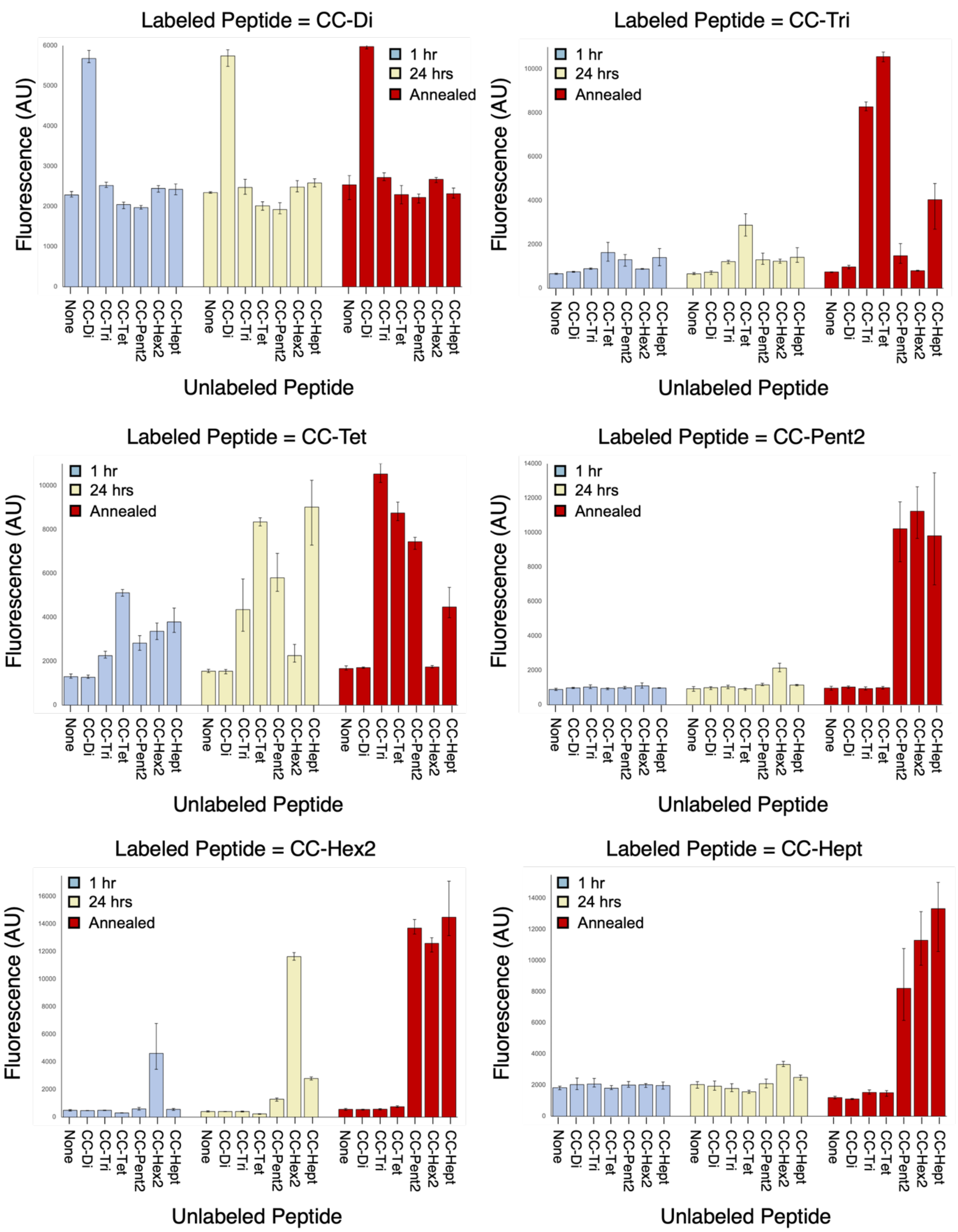


**Figure S36. Histogram representation of CC Basis Set fluorescence exchange data.** The raw fluorescence measurements of the Basis Set exchange studies are shown. The plotted fluorescence is the averaged value between three measurements. The top and bottom of the error bars are the maximum and minimum fluorescence values observed for each mixture respectively.

**Table S6. Raw and Normalized CC Basis Set Fluorescence Data**

| FAM-labelled Peptide | Unlabelled Peptide | 1 hr-raw | | | 1 hr-normalized | | | Avg | Std Dev |
|---|---|---|---|---|---|---|---|---|---|
| FAM-CC-Di | - | 2235 | 2372 | 2256 | -0.087 | -0.048 | -0.081 | -0.072 | 0.021 |
| FAM-CC-Di | CC-Di | 5577 | 5880 | 5582 | 0.884 | 0.972 | 0.885 | 0.914 | 0.050 |
| FAM-CC-Di | CC-Tri | 2470 | 2488 | 2603 | -0.087 | -0.048 | -0.081 | -0.072 | 0.021 |
| FAM-CC-Di | CC-Tet | 2106 | 2082 | 1944 | -0.060 | -0.069 | -0.122 | -0.084 | 0.033 |
| FAM-CC-Di | CC-Pent2 | 2028 | 1959 | 1933 | -0.090 | -0.116 | -0.126 | -0.111 | 0.019 |
| FAM-CC-Di | CC-Hex2 | 2354 | 2520 | 2471 | -0.053 | -0.005 | -0.019 | -0.025 | 0.025 |
| FAM-CC-Di | CC-Hept | 2420 | 2562 | 2289 | 0.060 | 0.114 | 0.010 | 0.061 | 0.052 |
| FAM-CC-Tri | - | 631 | 685 | 671 | -0.015 | -0.007 | -0.009 | -0.010 | 0.004 |
| FAM-CC-Tri | CC-Di | 758 | 728 | 769 | 0.002 | -0.002 | 0.004 | 0.001 | 0.003 |
| FAM-CC-Tri | CC-Tri | 867 | 928 | 879 | 0.017 | 0.025 | 0.018 | 0.020 | 0.004 |
| FAM-CC-Tri | CC-Tet | 1231 | 1574 | 2094 | 0.027 | 0.068 | 0.131 | 0.075 | 0.052 |
| FAM-CC-Tri | CC-Pent2 | 1008 | 1379 | 1536 | 0.000 | 0.045 | 0.064 | 0.036 | 0.033 |
| FAM-CC-Tri | CC-Hex2 | 871 | 897 | 861 | 0.017 | 0.021 | 0.016 | 0.018 | 0.002 |
| FAM-CC-Tri | CC-Hept | 1031 | 1358 | 1812 | 0.003 | 0.042 | 0.097 | 0.047 | 0.047 |
| FAM-CC-Tet | - | 1242 | 1229 | 1420 | -0.060 | -0.062 | -0.035 | -0.053 | 0.015 |
| FAM-CC-Tet | CC-Di | 1221 | 1220 | 1366 | -0.063 | -0.063 | -0.043 | -0.057 | 0.012 |
| FAM-CC-Tet | CC-Tri | 2456 | 2141 | 2173 | 0.111 | 0.067 | 0.071 | 0.083 | 0.024 |
| FAM-CC-Tet | CC-Tet | 5114 | 5269 | 4964 | 0.486 | 0.508 | 0.465 | 0.487 | 0.022 |
| FAM-CC-Tet | CC-Pent2 | 3162 | 2818 | 2500 | 0.211 | 0.162 | 0.117 | 0.163 | 0.047 |
| FAM-CC-Tet | CC-Hex2 | 3742 | 3367 | 2991 | 0.293 | 0.240 | 0.187 | 0.240 | 0.053 |
| FAM-CC-Tet | CC-Hept | 4420 | 3623 | 3316 | 0.388 | 0.276 | 0.233 | 0.299 | 0.080 |
| FAM-CC-Pent2 | - | 880 | 833 | 958 | -0.008 | -0.012 | 0.000 | -0.007 | 0.006 |
| FAM-CC-Pent2 | CC-Di | 938 | 1014 | 992 | -0.002 | 0.006 | 0.004 | 0.003 | 0.004 |
| FAM-CC-Pent2 | CC-Tri | 940 | 1150 | 997 | -0.002 | 0.019 | 0.004 | 0.007 | 0.011 |
| FAM-CC-Pent2 | CC-Tet | 976 | 877 | 985 | 0.002 | -0.008 | 0.003 | -0.001 | 0.006 |
| FAM-CC-Pent2 | CC-Pent2 | 916 | 982 | 1067 | -0.004 | 0.003 | 0.011 | 0.003 | 0.008 |

| | | | | | | | | | |
|---|---|---|---|---|---|---|---|---|---|
| FAM-CC-Pent2 | CC-Hex2 | 966 | 1059 | 1269 | 0.001 | 0.010 | 0.031 | 0.014 | 0.016 |
| FAM-CC-Pent2 | CC-Hept | 991 | 961 | 956 | 0.003 | 0.000 | 0.000 | 0.001 | 0.002 |
| FAM-CC-Hex2 | - | 540 | 485 | 442 | -0.002 | -0.006 | -0.010 | -0.006 | 0.004 |
| FAM-CC-Hex2 | CC-Di | 459 | 481 | 470 | -0.008 | -0.006 | -0.007 | -0.007 | 0.001 |
| FAM-CC-Hex2 | CC-Tri | 503 | 492 | 477 | -0.005 | -0.006 | -0.007 | -0.006 | 0.001 |
| FAM-CC-Hex2 | CC-Tet | 297 | 317 | 294 | -0.037 | -0.036 | -0.038 | -0.037 | 0.001 |
| FAM-CC-Hex2 | CC-Pent2 | 510 | 692 | 572 | -0.019 | -0.004 | -0.014 | -0.013 | 0.008 |
| FAM-CC-Hex2 | CC-Hex2 | 3468 | 6785 | 3593 | 0.242 | 0.518 | 0.252 | 0.337 | 0.156 |
| FAM-CC-Hex2 | CC-Hept | 478 | 631 | 577 | -0.022 | -0.009 | -0.014 | -0.015 | 0.007 |
| FAM-CC-Hept | - | 1705 | 1859 | 1931 | 0.041 | 0.054 | 0.060 | 0.052 | 0.010 |
| FAM-CC-Hept | CC-Di | 2452 | 1929 | 1710 | 0.103 | 0.060 | 0.042 | 0.068 | 0.031 |
| FAM-CC-Hept | CC-Tri | 2426 | 1895 | 1878 | 0.101 | 0.057 | 0.056 | 0.071 | 0.026 |
| FAM-CC-Hept | CC-Tet | 1969 | 1721 | 1706 | 0.063 | 0.043 | 0.042 | 0.049 | 0.012 |
| FAM-CC-Hept | CC-Pent2 | 1891 | 2231 | 1864 | 0.057 | 0.085 | 0.055 | 0.065 | 0.017 |
| FAM-CC-Hept | CC-Hex2 | 2103 | 2099 | 1855 | 0.074 | 0.074 | 0.054 | 0.067 | 0.012 |
| FAM-CC-Hept | CC-Hept | 2201 | 1963 | 1739 | 0.082 | 0.063 | 0.044 | 0.063 | 0.019 |

| **FAM-labelled Peptide** | **Unlabelled Peptide** | **24 hr-raw** | | | **24 hr-normalized** | | | **Avg** | **Std Dev** |
|---|---|---|---|---|---|---|---|---|---|
| FAM-CC-Di | - | 2366 | 2328 | 2333 | -0.049 | -0.060 | -0.059 | -0.056 | 0.006 |
| FAM-CC-Di | CC-Di | 5900 | 5485 | 5847 | 0.978 | 0.857 | 0.962 | 0.932 | 0.066 |
| FAM-CC-Di | CC-Tri | 2445 | 2300 | 2675 | -0.026 | -0.069 | 0.040 | -0.018 | 0.055 |
| FAM-CC-Di | CC-Tet | 2116 | 1907 | 2022 | -0.056 | -0.136 | -0.092 | -0.095 | 0.040 |
| FAM-CC-Di | CC-Pent2 | 2093 | 1818 | 1859 | -0.065 | -0.170 | -0.155 | -0.130 | 0.057 |
| FAM-CC-Di | CC-Hex2 | 2441 | 2363 | 2644 | -0.028 | -0.050 | 0.031 | -0.015 | 0.042 |
| FAM-CC-Di | CC-Hept | 2687 | 2488 | 2574 | 0.162 | 0.086 | 0.119 | 0.122 | 0.038 |
| FAM-CC-Tri | - | 646 | 620 | 712 | -0.013 | -0.016 | -0.004 | -0.011 | 0.006 |
| FAM-CC-Tri | CC-Di | 767 | 629 | 794 | 0.003 | -0.015 | 0.007 | -0.001 | 0.012 |
| FAM-CC-Tri | CC-Tri | 1279 | 1118 | 1233 | 0.071 | 0.050 | 0.065 | 0.062 | 0.011 |

| | | | | | | | | | |
|---|---|---|---|---|---|---|---|---|---|
| FAM-CC-Tri | CC-Tet | 2849 | 3399 | 2378 | 0.222 | 0.288 | 0.165 | 0.225 | 0.062 |
| FAM-CC-Tri | CC-Pent2 | 1194 | 1606 | 1095 | 0.022 | 0.072 | 0.010 | 0.035 | 0.033 |
| FAM-CC-Tri | CC-Hex2 | 1232 | 1144 | 1329 | 0.065 | 0.054 | 0.078 | 0.066 | 0.012 |
| FAM-CC-Tri | CC-Hept | 1201 | 1852 | 1187 | 0.023 | 0.102 | 0.021 | 0.049 | 0.046 |
| FAM-CC-Tet | - | 1535 | 1483 | 1630 | -0.019 | -0.026 | -0.006 | -0.017 | 0.011 |
| FAM-CC-Tet | CC-Di | 1627 | 1425 | 1579 | -0.006 | -0.035 | -0.013 | -0.018 | 0.015 |
| FAM-CC-Tet | CC-Tri | 5748 | 3365 | 3939 | 0.576 | 0.239 | 0.320 | 0.379 | 0.176 |
| FAM-CC-Tet | CC-Tet | 8363 | 8161 | 8530 | 0.945 | 0.917 | 0.969 | 0.944 | 0.026 |
| FAM-CC-Tet | CC-Pent2 | 6915 | 5299 | 5179 | 0.741 | 0.513 | 0.496 | 0.583 | 0.137 |
| FAM-CC-Tet | CC-Hex2 | 2768 | 1957 | 2052 | 0.155 | 0.041 | 0.054 | 0.083 | 0.063 |
| FAM-CC-Tet | CC-Hept | 10242 | 7291 | 9528 | 1.210 | 0.794 | 1.110 | 1.038 | 0.217 |
| FAM-CC-Pent2 | - | 939 | 782 | 1058 | -0.002 | -0.018 | 0.000 | -0.006 | 0.010 |
| FAM-CC-Pent2 | CC-Di | 886 | 1042 | 1044 | -0.007 | 0.009 | 0.004 | 0.002 | 0.008 |
| FAM-CC-Pent2 | CC-Tri | 931 | 1137 | 1040 | -0.003 | 0.018 | 0.004 | 0.007 | 0.011 |
| FAM-CC-Pent2 | CC-Tet | 958 | 847 | 968 | 0.000 | -0.011 | 0.003 | -0.003 | 0.007 |
| FAM-CC-Pent2 | CC-Pent2 | 1116 | 1113 | 1248 | 0.016 | 0.016 | 0.029 | 0.020 | 0.008 |
| FAM-CC-Pent2 | CC-Hex2 | 2082 | 1912 | 2409 | 0.113 | 0.096 | 0.146 | 0.119 | 0.025 |
| FAM-CC-Pent2 | CC-Hept | 1185 | 1099 | 1157 | 0.023 | 0.014 | 0.020 | 0.019 | 0.004 |
| FAM-CC-Hex2 | - | 460 | 391 | 373 | -0.008 | -0.014 | -0.015 | -0.013 | 0.004 |
| FAM-CC-Hex2 | CC-Di | 416 | 396 | 415 | -0.012 | -0.014 | -0.012 | -0.012 | 0.001 |
| FAM-CC-Hex2 | CC-Tri | 441 | 372 | 397 | -0.010 | -0.016 | -0.013 | -0.013 | 0.003 |
| FAM-CC-Hex2 | CC-Tet | 197 | 239 | 251 | -0.046 | -0.042 | -0.038 | -0.042 | 0.004 |
| FAM-CC-Hex2 | CC-Pent2 | 1381 | 1371 | 1148 | 0.054 | 0.053 | 0.034 | 0.047 | 0.011 |
| FAM-CC-Hex2 | CC-Hex2 | 11610 | 11929 | 11367 | 0.919 | 0.945 | 0.899 | 0.921 | 0.023 |
| FAM-CC-Hex2 | CC-Hept | 2681 | 2910 | 2798 | 0.164 | 0.183 | 0.174 | 0.174 | 0.010 |
| FAM-CC-Hept | - | 2225 | 1805 | 2066 | 0.084 | 0.050 | 0.071 | 0.068 | 0.017 |
| FAM-CC-Hept | CC-Di | 2264 | 1855 | 1671 | 0.088 | 0.054 | 0.039 | 0.060 | 0.025 |
| FAM-CC-Hept | CC-Tri | 2095 | 1656 | 1585 | 0.074 | 0.037 | 0.032 | 0.048 | 0.023 |
| FAM-CC-Hept | CC-Tet | 1675 | 1571 | 1474 | 0.039 | 0.030 | 0.022 | 0.031 | 0.008 |

| | | | | | | | | | |
|---|---|---|---|---|---|---|---|---|---|
| FAM-CC-Hept | CC-Pent2 | 1820 | 2386 | 2075 | 0.051 | 0.098 | 0.072 | 0.074 | 0.023 |
| FAM-CC-Hept | CC-Hex2 | 3257 | 3517 | 3190 | 0.169 | 0.191 | 0.164 | 0.175 | 0.014 |
| FAM-CC-Hept | CC-Hept | 2521 | 2637 | 2324 | 0.109 | 0.118 | 0.093 | 0.107 | 0.013 |

| **FAM-labelled Peptide** | **Unlabelled Peptide** | **annealed-raw** | | | **annealed-normalized** | | | **Avg** | **Std Dev** |
|---|---|---|---|---|---|---|---|---|---|
| FAM-CC-Di | - | 2170 | 2669 | 2769 | -0.106 | 0.039 | 0.068 | 0.000 | 0.093 |
| FAM-CC-Di | CC-Di | 5943 | 6068 | 5920 | 0.990 | 1.026 | 0.983 | 1.000 | 0.023 |
| FAM-CC-Di | CC-Tri | 2837 | 2642 | 2676 | 0.087 | 0.031 | 0.041 | 0.053 | 0.030 |
| FAM-CC-Di | CC-Tet | 2297 | 2524 | 2064 | 0.013 | 0.100 | -0.076 | 0.012 | 0.088 |
| FAM-CC-Di | CC-Pent2 | 2272 | 2311 | 2084 | 0.003 | 0.018 | -0.069 | -0.016 | 0.046 |
| FAM-CC-Di | CC-Hex2 | 2720 | 2703 | 2593 | 0.053 | 0.049 | 0.017 | 0.040 | 0.020 |
| FAM-CC-Di | CC-Hept | 2287 | 2455 | 2208 | 0.009 | 0.073 | -0.021 | 0.020 | 0.048 |
| FAM-CC-Tri | - | 751 | 717 | 754 | 0.001 | -0.003 | 0.002 | 0.000 | 0.003 |
| FAM-CC-Tri | CC-Di | 1046 | 888 | 975 | 0.041 | 0.020 | 0.031 | 0.030 | 0.010 |
| FAM-CC-Tri | CC-Tri | 8243 | 8496 | 8093 | 0.995 | 1.029 | 0.976 | 1.000 | 0.027 |
| FAM-CC-Tri | CC-Tet | 10332 | 10578 | 10772 | 1.125 | 1.155 | 1.178 | 1.153 | 0.027 |
| FAM-CC-Tri | CC-Pent2 | 1133 | 1286 | 2033 | 0.015 | 0.033 | 0.124 | 0.057 | 0.058 |
| FAM-CC-Tri | CC-Hex2 | 781 | 794 | 822 | 0.005 | 0.007 | 0.011 | 0.008 | 0.003 |
| FAM-CC-Tri | CC-Hept | 2700 | 4778 | 4634 | 0.204 | 0.455 | 0.438 | 0.366 | 0.140 |
| FAM-CC-Tet | - | 1599 | 1618 | 1791 | -0.010 | -0.007 | 0.017 | 0.000 | 0.015 |
| FAM-CC-Tet | CC-Di | 1723 | 1671 | 1740 | 0.008 | 0.000 | 0.010 | 0.006 | 0.005 |
| FAM-CC-Tet | CC-Tri | 11003 | 10434 | 10148 | 1.318 | 1.238 | 1.197 | 1.251 | 0.061 |
| FAM-CC-Tet | CC-Tet | 8405 | 9248 | 8601 | 0.951 | 1.070 | 0.979 | 1.000 | 0.062 |
| FAM-CC-Tet | CC-Pent2 | 7101 | 7645 | 7586 | 0.767 | 0.844 | 0.835 | 0.815 | 0.042 |
| FAM-CC-Tet | CC-Hex2 | 1708 | 1697 | 1800 | 0.005 | 0.004 | 0.018 | 0.009 | 0.008 |
| FAM-CC-Tet | CC-Hept | 5367 | 4066 | 3985 | 0.522 | 0.338 | 0.327 | 0.396 | 0.110 |
| FAM-CC-Pent2 | - | 972 | 821 | 1076 | 0.002 | -0.014 | 0.012 | 0.000 | 0.013 |
| FAM-CC-Pent2 | CC-Di | 947 | 1045 | 1093 | -0.001 | 0.009 | 0.014 | 0.007 | 0.007 |

| | | | | | | | | | |
|---|---|---|---|---|---|---|---|---|---|
| FAM-CC-Pent2 | CC-Tri | 915 | 1039 | 852 | -0.004 | 0.008 | -0.010 | -0.002 | 0.010 |
| FAM-CC-Pent2 | CC-Tet | 1000 | 913 | 1063 | 0.004 | -0.004 | 0.011 | 0.004 | 0.008 |
| FAM-CC-Pent2 | CC-Pent2 | 11785 | 10334 | 10570 | 1.089 | 0.943 | 0.967 | 1.000 | 0.078 |
| FAM-CC-Pent2 | CC-Hex2 | 12662 | 9654 | 11397 | 1.178 | 0.875 | 1.050 | 1.034 | 0.152 |
| FAM-CC-Pent2 | CC-Hept | 13474 | 10814 | 8992 | 1.259 | 0.992 | 0.808 | 1.020 | 0.227 |
| FAM-CC-Hex2 | - | 632 | 499 | 546 | 0.006 | -0.005 | -0.001 | 0.000 | 0.006 |
| FAM-CC-Hex2 | CC-Di | 567 | 508 | 572 | 0.001 | -0.004 | 0.001 | -0.001 | 0.003 |
| FAM-CC-Hex2 | CC-Tri | 615 | 519 | 592 | 0.005 | -0.003 | 0.003 | 0.001 | 0.004 |
| FAM-CC-Hex2 | CC-Tet | 698 | 822 | 759 | -0.004 | 0.007 | 0.002 | 0.002 | 0.005 |
| FAM-CC-Hex2 | CC-Pent2 | 14317 | 13273 | 13498 | 1.148 | 1.059 | 1.078 | 1.095 | 0.046 |
| FAM-CC-Hex2 | CC-Hex2 | 13004 | 11976 | 12783 | 1.035 | 0.949 | 1.016 | 1.000 | 0.045 |
| FAM-CC-Hex2 | CC-Hept | 17102 | 13215 | 13161 | 1.383 | 1.055 | 1.050 | 1.163 | 0.191 |
| FAM-CC-Hept | - | 2225 | 1805 | 2066 | 0.006 | -0.007 | 0.001 | 0.000 | 0.007 |
| FAM-CC-Hept | CC-Di | 2264 | 1855 | 1671 | -0.005 | -0.007 | -0.011 | -0.008 | 0.003 |
| FAM-CC-Hept | CC-Tri | 2095 | 1656 | 1585 | 0.018 | 0.024 | 0.040 | 0.027 | 0.011 |
| FAM-CC-Hept | CC-Tet | 1675 | 1571 | 1474 | 0.006 | 0.034 | 0.037 | 0.026 | 0.017 |
| FAM-CC-Hept | CC-Pent2 | 1820 | 2386 | 2075 | 0.789 | 0.408 | 0.537 | 0.578 | 0.194 |
| FAM-CC-Hept | CC-Hex2 | 3257 | 3517 | 3190 | 0.984 | 0.701 | 0.812 | 0.832 | 0.143 |
| FAM-CC-Hept | CC-Hept | 2521 | 2637 | 2324 | 1.088 | 0.773 | 1.139 | 1.000 | 0.198 |

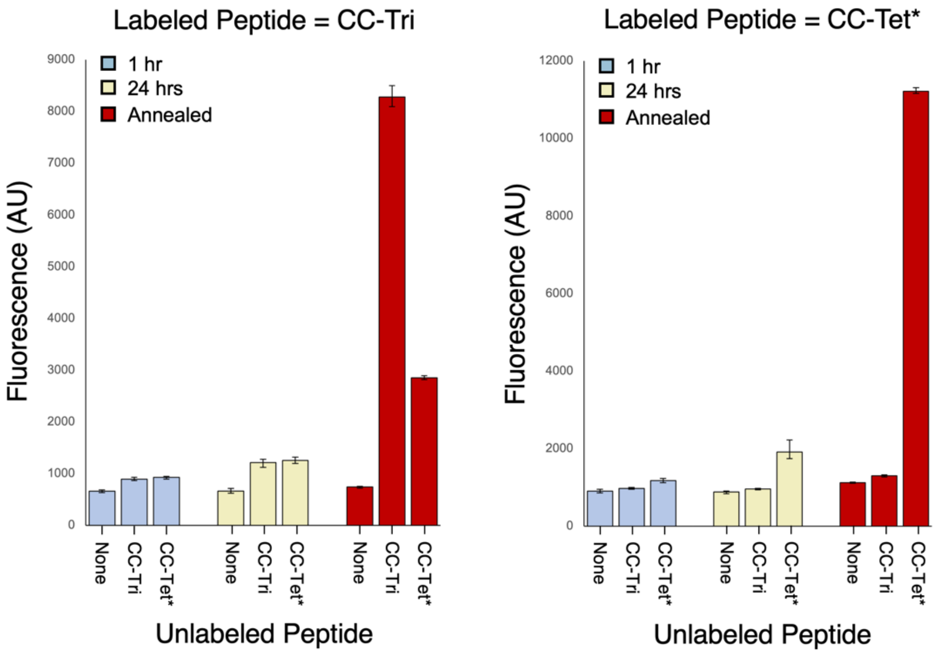


**Figure S37.** Histogram representation of CC-Tri and CC-Tet* fluorescence exchange data. The raw fluorescence measurements of the CC-Tri and CC-Tet* exchange studies are shown. The plotted fluorescence is the averaged value between three measurements. The top and bottom of the error bars are the maximum and minimum fluorescence values observed for each mixture respectively.

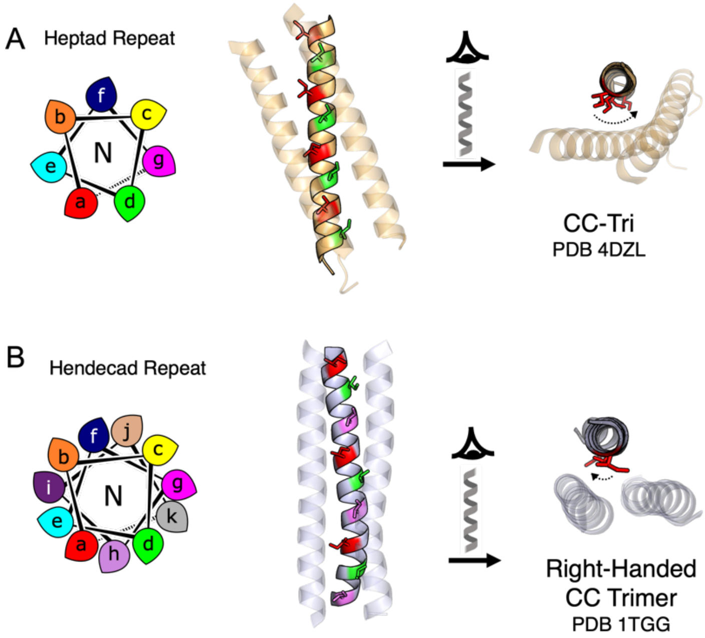


**Figure S38.** A comparison of the canonical CC heptad repeat and noncanonical hendecad repeat. A) Helical wheel representation of a heptad repeat where positions ***a*** and ***d*** (red and green, respectively) contribute to the coiled-coil interface. This is illustrated by CC-Tri,[1] which forms a left-handed coiled coil. (B) A similar helical wheel for a hendecad repeat where positions ***a***, ***d*** and ***h*** (colored red, green, and lilac, respectively) define the coiled-coil interface. This is illustrated by the PDB entry 1tgg, which forms a straightened coiled-coil trimer.[21]

**Table S7. Orthogonal Set Sequences**

| *g*-Register CC | ***gabcdef gabcdef gabcdef gabcdef*** | Monoistopic Mass |
|---|---|---|
| CC-Di | Ac-G EIAALKQ EIAALKK ENAALKW EIAALKQ G-CONH2 | 3246.9 |
| FAM-CC-Di | FAM-GGGG EIAALKQ EIAALKK ENAALKW EIAALKQ G-CONH2 | 3737.0 |
| CC-Tri | Ac-G EIAAIKQ EIAAIKK EIAAIKW EIAAIKQ G-CONH2 | 3245.9 |
| FAM-CC Tri | FAM-GGGG EIAAIKQ EIAAIKK EIAAIKW EIAAIKQ G-CONH2 | 3735.0 |
| ***c*-Register CC** | ***cdefgab cdefgab cdefgab cdefgab*** | |
| CC-Tet* | Ac-G EIQKQLK EIQKQLK EIQWQLK EIQKQLK G-CONH2 | 3703.1 |
| FAM-CC-Tet* | FAM-GGGG EIQKQLK EIQKQLK EIQWQLK EIQKQLK G-CONH2 | 4192.3 |
| CC-Hex2 | Ac-G EIAQSLK EIAKSLK EIAWSLK EIAQSLK G-CONH2 | 3309.9 |
| FAM-CC-Hex2 | FAM-GGGG EIAQSLK EIAKSLK EIAWSLK EIAQSLK G-CONH2 | 3800.0 |
| | ***cdefgab cdefgab cdefghijkab cdefgab*** | |
| CC-Pent2-hen3 | Ac-G EIAQALK EIAKALK EIAWALAAALK EIAQALK G-CONH2 | 3692.1 |
| FAM-CC-Pent2-hen3 | FAM-GGGG EIAQALK EIAKALK EIAWALAAALK EIAQALK G-CONH2 | 4182.3 |
| | ***cdefgab cdefghijkab cdefgab cdefgab*** | |
| CC-Hept-IV-hen2 | Ac-G EVAQAIK EVAKAVAAAIK EVAWAIK EVAQAIK G-CONH2 | 3502.0 |
| FAM-CC-Hept-IV-hen2 | FAM-GGGG EVAQAIK EVAKAVAAAIK EVAWAIK EVAQAIK G-CONH2 | 3992.1 |

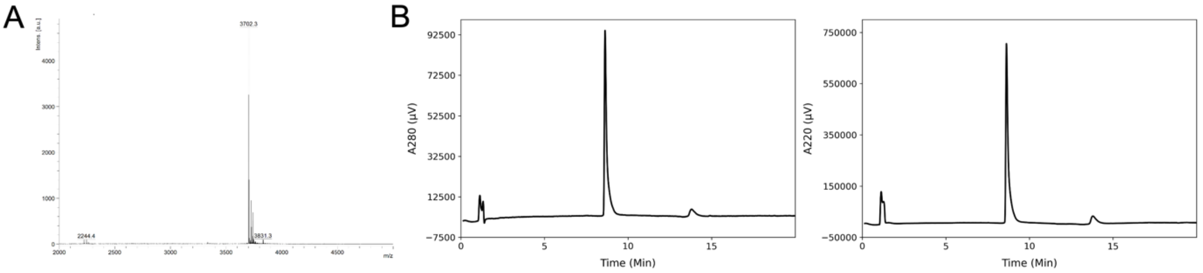


**Figure S39. MALDI and AHPLC of CC-Tet*.** (A) MALDI-TOF MS analysis of purified CC-Tet*. Monoisotopic [M+H] peaks were observed at 3702.3; calculated m/z is 3703.1. (B) Analytical HPLC trace of purified CC-Tet*. Chromatograms were monitored at 280 (left) and 220 (right) nm wavelengths.

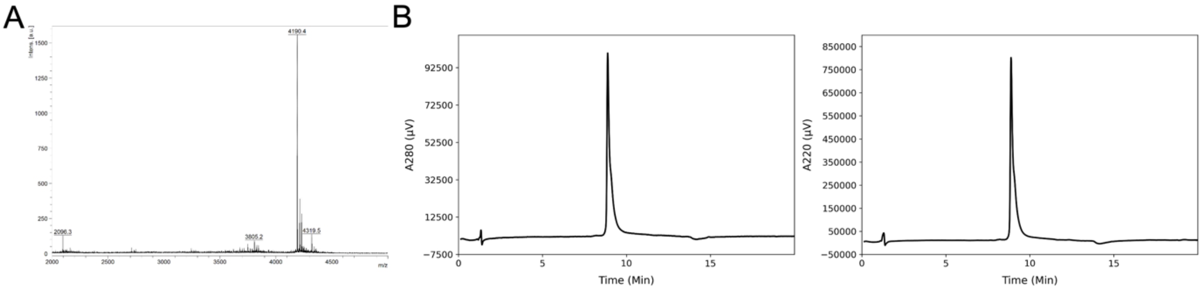


**Figure S40. MALDI and AHPLC of FAM-CC-Tet*.** (A) MALDI-TOF MS analysis of purified FAM-CC-Tet*. Monoisotopic [M+H] peaks were observed at 4190.4; calculated m/z is 4192.3. (B) Analytical HPLC trace of purified FAM-CC-Tet*. Chromatograms were monitored at 280 (left) and 220 (right) nm wavelengths.

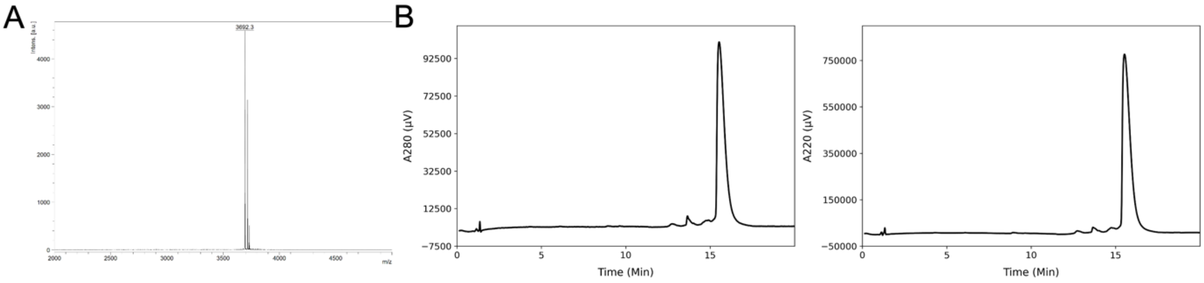


**Figure S41. MALDI and AHPLC of CC-Pent2-hen3.** (A) MALDI-TOF MS analysis of purified CC-Pent2-hen3. Monoisotopic [M+H] peaks were observed at 3692.3; calculated m/z is

3692.1. (B) Analytical HPLC trace of purified CC-Pent2-hen3. Chromatograms were monitored at 280 (left) and 220 (right) nm wavelengths.

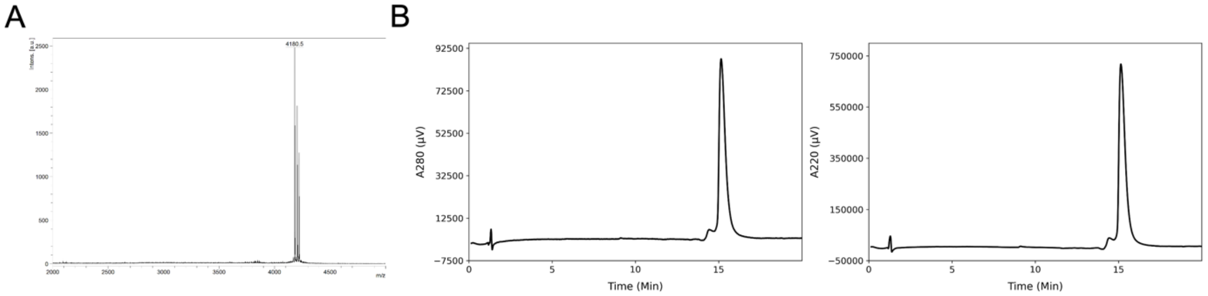


**Figure S42. MALDI and AHPLC of FAM-CC-Pent2-hen3.** (A) MALDI-TOF MS analysis of purified FAM-CC-Pent2-hen3. Monoisotopic [M+H] peaks were observed at 4180.5; calculated m/z is 4182.3. (B) Analytical HPLC trace of purified FAM-CC-Pent2-hen3. Chromatograms were monitored at 280 (left) and 220 (right) nm wavelengths.

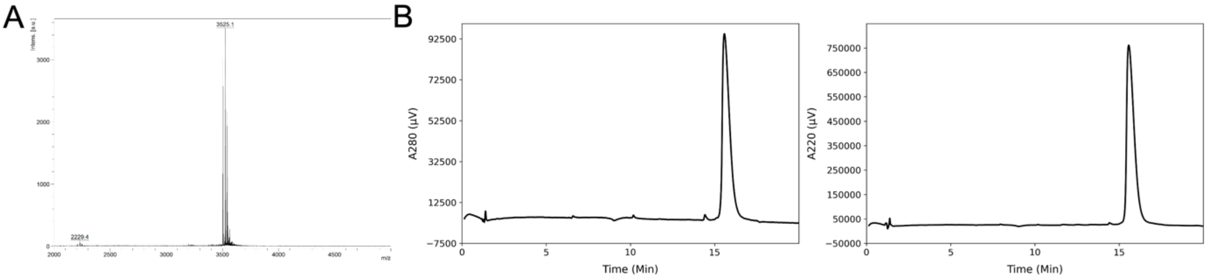


**Figure S43. MALDI and AHPLC of CC-Hept-IV-hen2.** (A) MALDI-TOF MS analysis of purified CC-Hept-IV-hen2. Sodium adduct [M+Na] peaks were observed at 3525.1; calculated m/z is 3525.0. (B) Analytical HPLC trace of purified CC-Hept-IV-hen2. Chromatograms were monitored at 280 (left) and 220 (right) nm wavelengths.

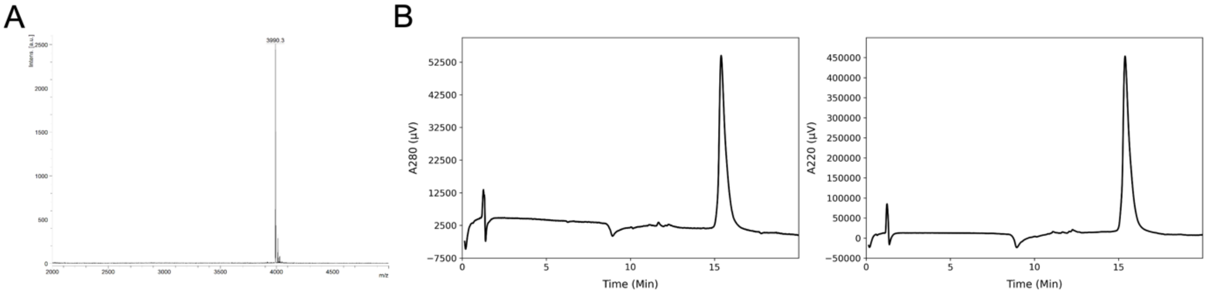


**Figure S44. MALDI and AHPLC of FAM-CC-Hept-IV-hen2.** (A) MALDI-TOF MS analysis of purified CC-Hept-IV-hen2. Sodium adduct [M+Na] peaks were observed at 3990.3; calculated m/z is 3992.1. (B) Analytical HPLC trace of purified CC-Hept-IV-hen2. Chromatograms were monitored at 280 (left) and 220 (right) nm wavelengths.

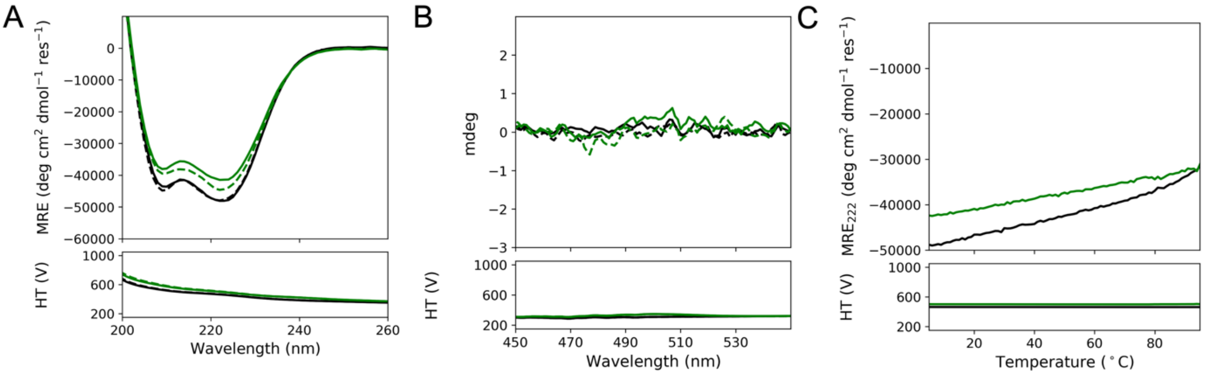


**Figure S45. CD Characterization of CC-Tet* and FAM-CC-Tet*.** CD spectra of 50 µM CC-Tet* (black) and FAM-CC-Tet* (green) in PBS at 5 °C are shown in (A) and (B). Samples were analyzed prior to melting (solid lines) and post-melting (dotted lines). (A) shows the spectra between 260-200 nm and (B) shows the spectra between 550-450 nm. (C) shows the thermal melt of the two peptides from 5-95 °C. There is not a noticeable difference in the CD characterization of the unlabeled and labeled variants of CC-Tet*.

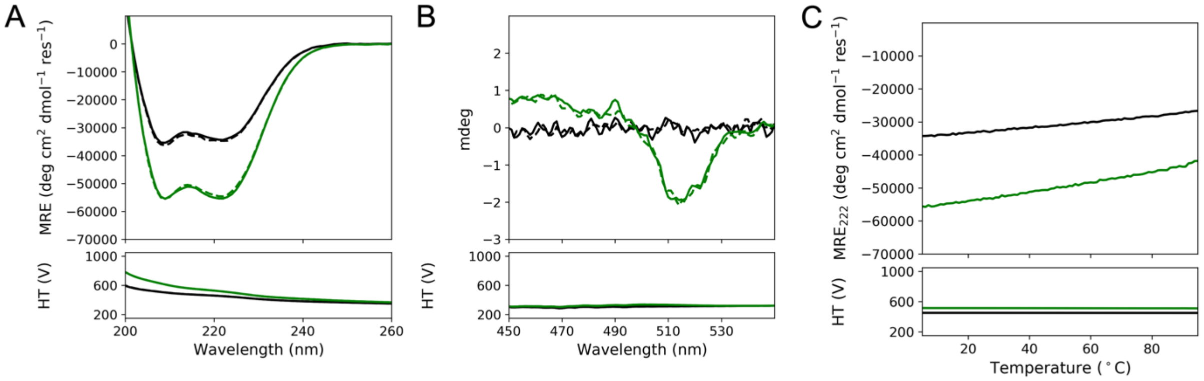


**Figure S46. CD Characterization of CC-Pent2-hen3 and FAM- Pent2-hen3.** CD spectra of 50 µM CC-Pent2-hen3 (black) and FAM-CC-Pent2-hen3 (green) in PBS at 5 °C are shown in (A) and (B). Samples were analyzed prior to melting (solid lines) and post-melting (dotted lines). (A) shows the spectra between 260-200 nm and (B) shows the spectra between 550-450 nm. (C) shows the thermal melt of the two peptides from 5-95 °C. The labeled peptide shows an increase in CD signal (A) as well as evidence of the Cotton effect (B). However, the thermal melts are consistent between the samples, indicative that the FAM-labeled peptide has the same thermodynamic properties as the unlabeled peptide.

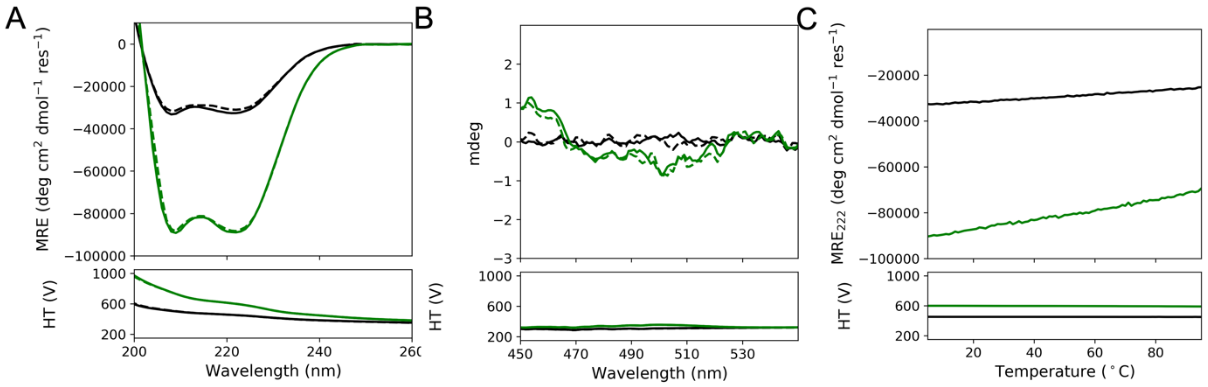


**Figure S47. CD Characterization of CC-Hept-IV-hen2 and FAM-Hept-IV-hen2.** CD spectra of 50 µM CC-Hept-IV-hen2 (black) and 25 µM FAM-CC-Hept-IV-hen2 (green) in PBS at 5 °C are shown in (A) and (B). Samples were analyzed prior to melting (solid lines) and post-melting (dotted lines). (A) shows the spectra between 260-200 nm and (B) shows the spectra between 550-450 nm. (C) shows the thermal melt of the two peptides from 5-95 °C. The labeled peptide shows an increase in CD signal (A). However, the thermal melts are consistent between the samples, indicative that the FAM-labeled peptide has the same thermodynamic properties as the unlabeled peptide.

**Table S8.** Table 1 of CC-Hex2-hen2 with Ala at ***h*****,** CC-Hept-hen2 with Ala at ***h**,* and CC-Hept-IV-hen2 stuctures.

| | **CC-Hex2-hen2, Ala at *h* (PDB 9gf2)** | **CC-Hept-hen2, Ala at *h* (PDB 9gf3)** | **CC-Hept-IV-hen2 (PDB 9gf4)** |
|---|---|---|---|
| Wavelength (Å) | 0.9795 | 0.9795 | 0.9795 |
| Resolution Range | 48.56 - 2.10 (2.31 - 2.1) | 47.51-1.65 (1.67-1.65) | 19.67-1.49 (1.52-1.49) |
| Space Group | P 2 21 21 | P 21 21 21 | P 2 21 21 |
| Unit Cell | 48.563 58.157 59.258 90 90 90 | 60.267 61.754 154.441 90 90 90 | 38.546 47.919 153.02 90 90 90 |
| Unique Reflections | 10246 (2510) | 70117 (2831) | 47044 (2595) |
| Completeness (%) | 99.59 (99.84) | 99.83 (100.00) | 99.57 (95.65) |
| Wilson B-factor | 33.40 | 21.79 | 20.26 |
| Reflections used in refinement | 10246 (2510) | 70117 (2831) | 47044 (2595) |
| Reflections used for R-free | 532(114) | 3455 (135) | 2431 (148) |
| R-work | 0.2252 (0.1904) | 0.1939 (0.3003) | 0.1715 (0.2311) |
| R-free | 0.2220 (0.1944) | 0.2148 (0.3255) | 0.1972 (0.2912) |
| Number of non-hydrogen atoms | 1426 | 4046 | 1999 |
| macromolecules | 1372 | 3474 | 1716 |
| ligands | 16 | 173 | 90 |
| solvent | 38 | 399 | 193 |
| Protein Residues | 190 | 476 | 238 |
| RMS(bonds) | 0.022 | 0.059 | 0.064 |
| RMS(angles) | 2.52 | 1.49 | 1.27 |
| Ramachandran favored(%) | 96.63 | 100.00 | 100.00 |
| Ramachandran allowed(%) | 1.69 | 0.00 | 0.00 |
| Ramachandran outliers(%) | 1.69 | 0.00 | 0.00 |
| Rotamer outliers(%) | 7.81 | 0.00 | 0.00 |
| Clashscore | 7.22 | 2.88 | 1.33 |
| Average B-factor | 52.62 | 29.07 | 25.71 |
| macromolecules | 53.04 | 26.47 | 22.82 |
| ligands | 43.57 | 55.53 | 51.11 |
| solvent | 41.15 | 40.22 | 39.54 |

**Table S9.** Oligomeric states of CC-Pent2-hen3 and CC-Hept-IV-hen2 measured by AUC (SV and SE) and in crystal structures.

| Peptide | Oligomeric state based on SV data | Oligomeric state based on SV data | Oligomeric state based on crystallography data |
|---|---|---|---|
| CC-Pent2-hen3 | 5.3 | 4.9 | ND |
| CC-Hept-IV-hen2 | 6.1 | 6.0 | 7 |

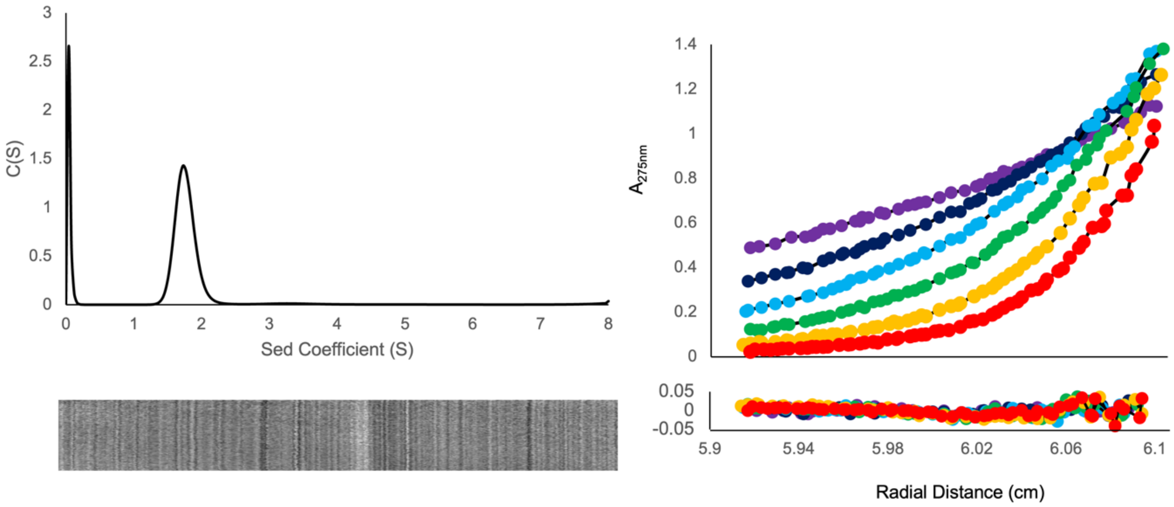


**Figure S48**. AUC data for CC-Pent2-hen3. Left shows the sedimentation velocity (SV) data acquired for CC-Pent2-hen3. The molecular weight calculated from the data is 19558 Da = 5.3 x the monomeric mass of CC-Pent2-hen3 (3693.4 Da) ($f/f_0$ = 1.225, s = 1.757, sw(20,w) = 1.798 S). Right shows the sedimentation equilibrium data acquired for CC-Pent2-hen3. The molecular weight derived from the data is 18098 Da = 4.9 x the monomeric mass of CC-Pent2-hen3 ($\chi^2$ = 4.98).

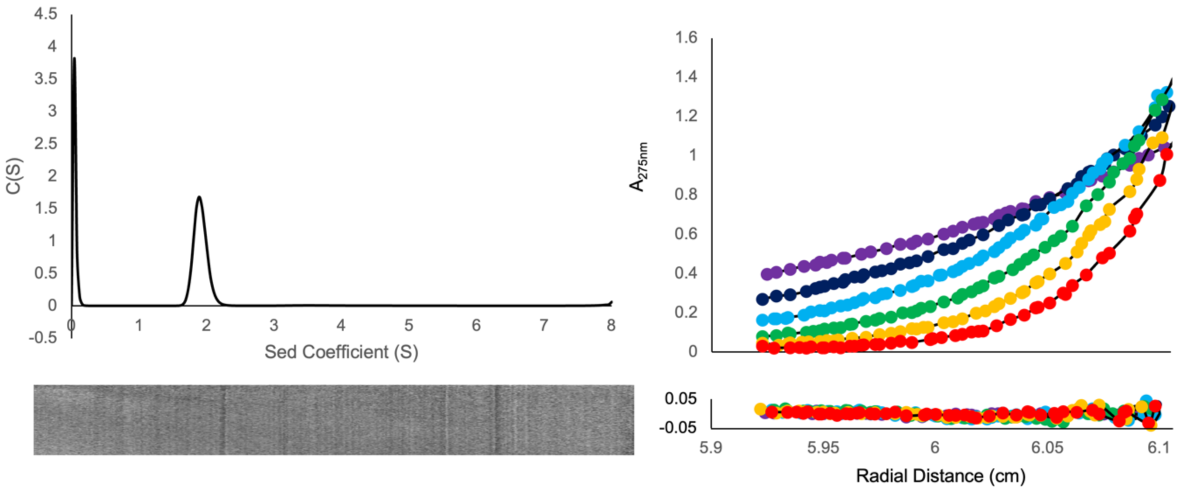


**Figure S49**. AUC data for CC-Hept-IV-hen2. Left shows the sedimentation velocity (SV) data acquired for CC-Hept-IV-hen2. The molecular weight calculated from the data is 21185 Da = 6.1 x the monomeric mass of CC-Hept-IV-hen2 (3503.1 Da) ($f/f_0$ = 1.204, s = 1.909, sw(20,w) = 1.953 S). Right shows the sedimentation equilibrium data acquired for CC-Hept-IV-hen2. The molecular weight derived from the data is 20904 Da = 6.0 x the monomeric mass of CC-Hept-IV-hen2 ($\chi^2$ = 3.94).

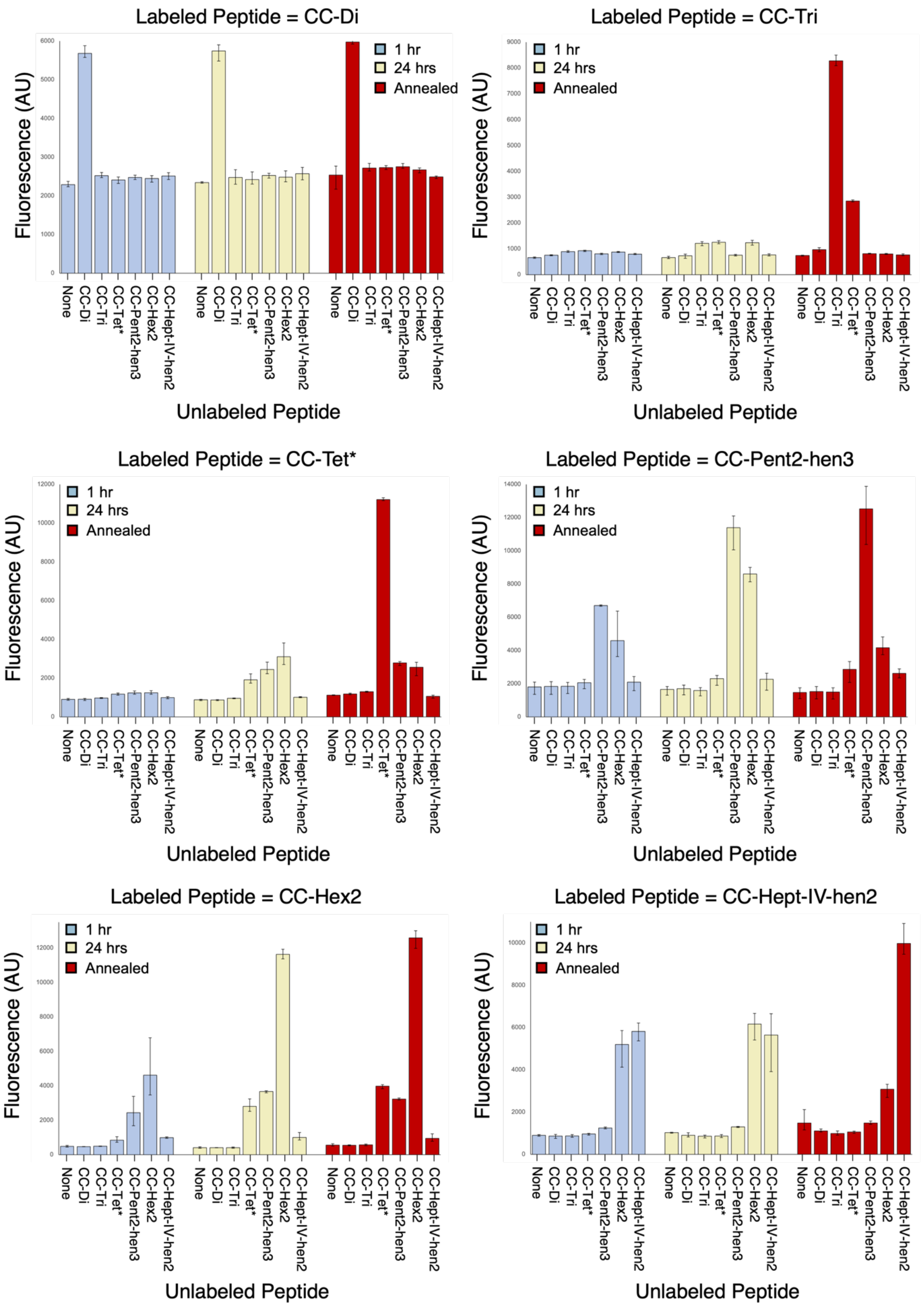


**Figure S50. Histogram representation of Orthogonal CC Set fluorescence exchange data.** The raw fluorescence measurements of the Orthogonal CC Set exchange studies are shown. The plotted fluorescence is the averaged value between three measurements. The top and bottom of the error bars are the maximum and minimum fluorescence values observed for each mixture respectively.

**Table S10. Orthogonal CC Set Fluorescence Data**

| FAM-labelled Peptide | Unlabelled Peptide | 1 hr-raw | | | 1 hr-normalized | | | Avg | Std Dev |
|---|---|---|---|---|---|---|---|---|---|
| FAM-CC-Di | - | 2235 | 2372 | 2256 | -0.087 | -0.048 | -0.081 | -0.072 | 0.021 |
| FAM-CC-Di | CC-Di | 5577 | 5880 | 5582 | 0.884 | 0.972 | 0.885 | 0.914 | 0.050 |
| FAM-CC-Di | CC-Tri | 2470 | 2488 | 2603 | -0.019 | -0.014 | 0.019 | -0.005 | 0.021 |
| FAM-CC-Di | CC-Tet* | 2319 | 2486 | 2418 | -0.063 | -0.015 | -0.034 | -0.037 | 0.024 |
| FAM-CC-Di | CC-Pent2-hen3 | 2467 | 2533 | 2419 | -0.020 | -0.001 | -0.034 | -0.018 | 0.017 |
| FAM-CC-Di | CC-Hex2 | 2354 | 2520 | 2471 | -0.053 | -0.005 | -0.019 | -0.025 | 0.025 |
| FAM-CC-Di | CC-Hept-IV-hen2 | 2421 | 2598 | 2523 | -0.033 | 0.018 | -0.004 | -0.006 | 0.026 |
| FAM-CC-Tri | - | 631 | 685 | 671 | -0.015 | -0.007 | -0.009 | -0.010 | 0.004 |
| FAM-CC-Tri | CC-Di | 758 | 728 | 769 | 0.002 | -0.002 | 0.004 | 0.001 | 0.003 |
| FAM-CC-Tri | CC-Tri | 867 | 928 | 879 | 0.017 | 0.025 | 0.018 | 0.020 | 0.004 |
| FAM-CC-Tri | CC-Tet* | 942 | 948 | 891 | 0.027 | 0.028 | 0.020 | 0.025 | 0.004 |
| FAM-CC-Tri | CC-Pent2-hen3 | 778 | 829 | 804 | 0.005 | 0.012 | 0.008 | 0.008 | 0.003 |
| FAM-CC-Tri | CC-Hex2 | 871 | 897 | 861 | 0.017 | 0.021 | 0.016 | 0.018 | 0.002 |
| FAM-CC-Tri | CC-Hept-IV-hen2 | 823 | 776 | 783 | 0.011 | 0.005 | 0.006 | 0.007 | 0.003 |
| FAM-CC-Tet* | - | 860 | 947 | 908 | -0.026 | -0.017 | -0.021 | -0.021 | 0.004 |
| FAM-CC-Tet* | CC-Di | 843 | 943 | 953 | -0.027 | -0.017 | -0.016 | -0.020 | 0.006 |
| FAM-CC-Tet* | CC-Tri | 954 | 1000 | 960 | -0.016 | -0.012 | -0.016 | -0.015 | 0.002 |
| FAM-CC-Tet* | CC-Tet* | 1125 | 1227 | 1173 | 0.001 | 0.011 | 0.005 | 0.006 | 0.005 |
| FAM-CC-Tet* | CC-Pent2-hen3 | 1209 | 1340 | 1167 | 0.009 | 0.022 | 0.005 | 0.012 | 0.009 |
| FAM-CC-Tet* | CC-Hex2 | 1217 | 1351 | 1171 | 0.010 | 0.023 | 0.005 | 0.013 | 0.009 |
| FAM-CC-Tet* | CC-Hept-IV-hen2 | 1025 | 1043 | 932 | -0.009 | -0.008 | -0.019 | -0.012 | 0.006 |
| FAM-CC-Pent2-hen3 | - | 2096 | 1962 | 1353 | 0.056 | 0.044 | -0.011 | 0.030 | 0.036 |
| FAM-CC-Pent2-hen3 | CC-Di | 2028 | 2120 | 1356 | 0.050 | 0.058 | -0.011 | 0.033 | 0.038 |
| FAM-CC-Pent2-hen3 | CC-Tri | 2074 | 2053 | 1404 | 0.054 | 0.052 | -0.006 | 0.033 | 0.034 |
| FAM-CC-Pent2-hen3 | CC-Tet* | 2226 | 2257 | 1696 | 0.068 | 0.071 | 0.020 | 0.053 | 0.029 |
| FAM-CC-Pent2-hen3 | CC-Pent2-hen3 | 6715 | 6660 | 6742 | 0.474 | 0.469 | 0.477 | 0.474 | 0.004 |

| | | | | | | | | | |
|---|---|---|---|---|---|---|---|---|---|
| FAM-CC-Pent2-hen3 | CC-Hex2 | 3753 | 6373 | 3633 | 0.206 | 0.443 | 0.195 | 0.282 | 0.140 |
| FAM-CC-Pent2-hen3 | CC-Hept-IV-hen2 | 2430 | 2282 | 1581 | 0.086 | 0.073 | 0.010 | 0.056 | 0.041 |
| FAM-CC-Hex2 | - | 937 | 916 | 854 | -0.064 | -0.066 | -0.074 | -0.068 | 0.005 |
| FAM-CC-Hex2 | CC-Di | 904 | 946 | 749 | -0.068 | -0.063 | -0.086 | -0.072 | 0.012 |
| FAM-CC-Hex2 | CC-Tri | 875 | 935 | 805 | -0.071 | -0.064 | -0.079 | -0.072 | 0.008 |
| FAM-CC-Hex2 | CC-Tet* | 989 | 985 | 909 | -0.058 | -0.058 | -0.067 | -0.061 | 0.005 |
| FAM-CC-Hex2 | CC-Pent2-hen3 | 1292 | 1208 | 1249 | -0.022 | -0.032 | -0.027 | -0.027 | 0.005 |
| FAM-CC-Hex2 | CC-Hex2 | 4124 | 5615 | 5855 | 0.311 | 0.487 | 0.515 | 0.438 | 0.110 |
| FAM-CC-Hex2 | CC-Hept-IV-hen2 | 5873 | 6211 | 5365 | 0.517 | 0.557 | 0.458 | 0.511 | 0.050 |
| FAM-CC-Hept-IV-hen2 | - | 1705 | 1859 | 1931 | 0.041 | 0.054 | 0.060 | 0.052 | 0.010 |
| FAM-CC-Hept-IV-hen2 | CC-Di | 2452 | 1929 | 1710 | 0.103 | 0.060 | 0.042 | 0.068 | 0.031 |
| FAM-CC-Hept-IV-hen2 | CC-Tri | 2426 | 1895 | 1878 | 0.101 | 0.057 | 0.056 | 0.071 | 0.026 |
| FAM-CC-Hept-IV-hen2 | CC-Tet* | 1969 | 1721 | 1706 | 0.063 | 0.043 | 0.042 | 0.049 | 0.012 |
| FAM-CC-Hept-IV-hen2 | CC-Pent2-hen3 | 1891 | 2231 | 1864 | 0.057 | 0.085 | 0.055 | 0.065 | 0.017 |
| FAM-CC-Hept-IV-hen2 | CC-Hex2 | 2103 | 2099 | 1855 | 0.074 | 0.074 | 0.054 | 0.067 | 0.012 |
| FAM-CC-Hept-IV-hen2 | CC-Hept-IV-hen2 | 2201 | 1963 | 1739 | 0.082 | 0.063 | 0.044 | 0.063 | 0.019 |

| **FAM-labelled Peptide** | **Unlabelled Peptide** | **24 hr-raw** | | | **24 hr-normalized** | | | **Avg** | **Std Dev** |
|---|---|---|---|---|---|---|---|---|---|
| FAM-CC-Di | - | 2366 | 2328 | 2333 | -0.049 | -0.060 | -0.059 | -0.056 | 0.006 |
| FAM-CC-Di | CC-Di | 5900 | 5485 | 5847 | 0.978 | 0.857 | 0.962 | 0.932 | 0.066 |
| FAM-CC-Di | CC-Tri | 2445 | 2300 | 2675 | -0.026 | -0.069 | 0.040 | -0.018 | 0.055 |
| FAM-CC-Di | CC-Tet* | 2304 | 2337 | 2617 | -0.067 | -0.058 | 0.024 | -0.034 | 0.050 |
| FAM-CC-Di | CC-Pent2-hen3 | 2583 | 2460 | 2533 | 0.014 | -0.022 | -0.001 | -0.003 | 0.018 |
| FAM-CC-Di | CC-Hex2 | 2441 | 2363 | 2644 | -0.028 | -0.050 | 0.031 | -0.015 | 0.042 |
| FAM-CC-Di | CC-Hept-IV-hen2 | 2571 | 2412 | 2732 | 0.010 | -0.036 | 0.057 | 0.010 | 0.046 |
| FAM-CC-Tri | - | 646 | 620 | 712 | -0.013 | -0.016 | -0.004 | -0.011 | 0.006 |
| FAM-CC-Tri | CC-Di | 767 | 629 | 794 | 0.003 | -0.015 | 0.007 | -0.001 | 0.012 |
| FAM-CC-Tri | CC-Tri | 1279 | 1118 | 1233 | 0.071 | 0.050 | 0.065 | 0.062 | 0.011 |

| | | | | | | | | | |
|---|---|---|---|---|---|---|---|---|---|
| FAM-CC-Tri | CC-Tet* | 1251 | 1194 | 1316 | 0.068 | 0.060 | 0.076 | 0.068 | 0.008 |
| FAM-CC-Tri | CC-Pent2-hen3 | 755 | 727 | 785 | 0.002 | -0.002 | 0.006 | 0.002 | 0.004 |
| FAM-CC-Tri | CC-Hex2 | 1232 | 1144 | 1329 | 0.065 | 0.054 | 0.078 | 0.066 | 0.012 |
| FAM-CC-Tri | CC-Hept-IV-hen2 | 808 | 725 | 765 | 0.009 | -0.002 | 0.003 | 0.003 | 0.006 |
| FAM-CC-Tet* | - | 903 | 836 | 894 | -0.021 | -0.028 | -0.022 | -0.024 | 0.004 |
| FAM-CC-Tet* | CC-Di | 836 | 888 | 890 | -0.028 | -0.023 | -0.023 | -0.025 | 0.003 |
| FAM-CC-Tet* | CC-Tri | 939 | 977 | 950 | -0.018 | -0.014 | -0.017 | -0.016 | 0.002 |
| FAM-CC-Tet* | CC-Tet* | 1743 | 2224 | 1775 | 0.062 | 0.109 | 0.065 | 0.079 | 0.027 |
| FAM-CC-Tet* | CC-Pent2-hen3 | 2275 | 2827 | 2231 | 0.114 | 0.169 | 0.110 | 0.131 | 0.033 |
| FAM-CC-Tet* | CC-Hex2 | 2807 | 3815 | 2698 | 0.167 | 0.267 | 0.156 | 0.197 | 0.061 |
| FAM-CC-Tet* | CC-Hept-IV-hen2 | 987 | 1006 | 1042 | -0.013 | -0.011 | -0.008 | -0.011 | 0.003 |
| FAM-CC-Pent2-hen3 | - | 1827 | 1818 | 1312 | 0.032 | 0.031 | -0.015 | 0.016 | 0.027 |
| FAM-CC-Pent2-hen3 | CC-Di | 1857 | 1915 | 1325 | 0.035 | 0.040 | -0.014 | 0.020 | 0.029 |
| FAM-CC-Pent2-hen3 | CC-Tri | 1749 | 1761 | 1262 | 0.025 | 0.026 | -0.019 | 0.011 | 0.026 |
| FAM-CC-Pent2-hen3 | CC-Tet* | 2496 | 2494 | 1906 | 0.092 | 0.092 | 0.039 | 0.075 | 0.031 |
| FAM-CC-Pent2-hen3 | CC-Pent2-hen3 | 12100 | 12029 | 10059 | 0.962 | 0.956 | 0.777 | 0.898 | 0.105 |
| FAM-CC-Pent2-hen3 | CC-Hex2 | 9002 | 8133 | 8658 | 0.681 | 0.603 | 0.650 | 0.645 | 0.040 |
| FAM-CC-Pent2-hen3 | CC-Hept-IV-hen2 | 2624 | 2577 | 1610 | 0.104 | 0.100 | 0.012 | 0.072 | 0.052 |
| FAM-CC-Hex2 | - | 460 | 391 | 373 | -0.008 | -0.014 | -0.015 | -0.013 | 0.004 |
| FAM-CC-Hex2 | CC-Di | 416 | 396 | 415 | -0.012 | -0.014 | -0.012 | -0.012 | 0.001 |
| FAM-CC-Hex2 | CC-Tri | 441 | 372 | 397 | -0.010 | -0.016 | -0.013 | -0.013 | 0.003 |
| FAM-CC-Hex2 | CC-Tet* | 2668 | 3237 | 2515 | 0.175 | 0.223 | 0.163 | 0.187 | 0.032 |
| FAM-CC-Hex2 | CC-Pent2-hen3 | 3710 | 3699 | 3584 | 0.262 | 0.261 | 0.251 | 0.258 | 0.006 |
| FAM-CC-Hex2 | CC-Hex2 | 11610 | 11929 | 11367 | 0.919 | 0.945 | 0.899 | 0.921 | 0.023 |
| FAM-CC-Hex2 | CC-Hept-IV-hen2 | 857 | 1287 | 850 | 0.025 | 0.061 | 0.024 | 0.036 | 0.021 |
| FAM-CC-Hept-IV-hen2 | - | 999 | 1042 | 1033 | -0.057 | -0.051 | -0.053 | -0.054 | 0.003 |
| FAM-CC-Hept-IV-hen2 | CC-Di | 862 | 1020 | 825 | -0.073 | -0.054 | -0.077 | -0.068 | 0.012 |
| FAM-CC-Hept-IV-hen2 | CC-Tri | 886 | 909 | 772 | -0.070 | -0.067 | -0.083 | -0.073 | 0.009 |
| FAM-CC-Hept-IV-hen2 | CC-Tet* | 869 | 939 | 796 | -0.072 | -0.064 | -0.080 | -0.072 | 0.008 |

| | | | | | | | | | |
|---|---|---|---|---|---|---|---|---|---|
| FAM-CC-Hept-IV-hen2 | CC-Pent2-hen3 | 1293 | 1271 | 1320 | -0.022 | -0.025 | -0.019 | -0.022 | 0.003 |
| FAM-CC-Hept-IV-hen2 | CC-Hex2 | 6400 | 5410 | 6664 | 0.579 | 0.463 | 0.611 | 0.551 | 0.078 |
| FAM-CC-Hept-IV-hen2 | CC-Hept-IV-hen2 | 6363 | 6653 | 3908 | 0.575 | 0.609 | 0.286 | 0.490 | 0.178 |

| FAM-labelled Peptide | Unlabelled Peptide | annealed-raw | | | annealed-normalized | | | Avg | Std Dev |
|---|---|---|---|---|---|---|---|---|---|
| FAM-CC-Di | - | 2170 | 2669 | 2769 | -0.106 | 0.039 | 0.068 | 0.000 | 0.093 |
| FAM-CC-Di | CC-Di | 5943 | 6068 | 5920 | 0.990 | 1.026 | 0.983 | 1.000 | 0.023 |
| FAM-CC-Di | CC-Tri | 2837 | 2642 | 2676 | 0.087 | 0.031 | 0.041 | 0.053 | 0.030 |
| FAM-CC-Di | CC-Tet* | 2679 | 2730 | 2779 | 0.042 | 0.056 | 0.071 | 0.056 | 0.015 |
| FAM-CC-Di | CC-Pent2-hen3 | 2835 | 2713 | 2711 | 0.087 | 0.051 | 0.051 | 0.063 | 0.021 |
| FAM-CC-Di | CC-Hex2 | 2720 | 2703 | 2593 | 0.053 | 0.049 | 0.017 | 0.040 | 0.020 |
| FAM-CC-Di | CC-Hept-IV-hen2 | 2511 | 2510 | 2448 | -0.007 | -0.008 | -0.026 | -0.013 | 0.010 |
| FAM-CC-Tri | - | 751 | 717 | 754 | 0.001 | -0.003 | 0.002 | 0.000 | 0.003 |
| FAM-CC-Tri | CC-Di | 1046 | 888 | 975 | 0.041 | 0.020 | 0.031 | 0.030 | 0.010 |
| FAM-CC-Tri | CC-Tri | 8243 | 8496 | 8093 | 0.995 | 1.029 | 0.976 | 1.000 | 0.027 |
| FAM-CC-Tri | CC-Tet* | 2890 | 2818 | 2853 | 0.285 | 0.276 | 0.280 | 0.280 | 0.005 |
| FAM-CC-Tri | CC-Pent2-hen3 | 828 | 805 | 800 | 0.012 | 0.009 | 0.008 | 0.009 | 0.002 |
| FAM-CC-Tri | CC-Hex2 | 781 | 794 | 822 | 0.005 | 0.007 | 0.011 | 0.008 | 0.003 |
| FAM-CC-Tri | CC-Hept-IV-hen2 | 804 | 738 | 746 | 0.008 | 0.000 | 0.001 | 0.003 | 0.005 |
| FAM-CC-Tet* | - | 1104 | 1118 | 1136 | -0.002 | 0.000 | 0.002 | 0.000 | 0.002 |
| FAM-CC-Tet* | CC-Di | 1145 | 1224 | 1180 | 0.003 | 0.010 | 0.006 | 0.006 | 0.004 |
| FAM-CC-Tet* | CC-Tri | 1269 | 1319 | 1299 | 0.015 | 0.020 | 0.018 | 0.017 | 0.002 |
| FAM-CC-Tet* | CC-Tet* | 11307 | 11171 | 11160 | 1.009 | 0.996 | 0.995 | 1.000 | 0.008 |
| FAM-CC-Tet* | CC-Pent2-hen3 | 2657 | 2853 | 2843 | 0.152 | 0.172 | 0.171 | 0.165 | 0.011 |
| FAM-CC-Tet* | CC-Hex2 | 2129 | 2748 | 2815 | 0.100 | 0.161 | 0.168 | 0.143 | 0.037 |
| FAM-CC-Tet* | CC-Hept-IV-hen2 | 1086 | 1121 | 969 | -0.003 | 0.000 | -0.015 | -0.006 | 0.008 |
| FAM-CC-Pent2-hen3 | - | 1579 | 1748 | 1097 | 0.009 | 0.025 | -0.034 | 0.000 | 0.031 |
| FAM-CC-Pent2-hen3 | CC-Di | 1658 | 1821 | 1084 | 0.017 | 0.031 | -0.035 | 0.004 | 0.035 |

| | | | | | | | | | |
|---|---|---|---|---|---|---|---|---|---|
| FAM-CC-Pent2-hen3 | CC-Tri | 1664 | 1748 | 1071 | 0.017 | 0.025 | -0.037 | 0.002 | 0.033 |
| FAM-CC-Pent2-hen3 | CC-Tet* | 3167 | 3332 | 2078 | 0.153 | 0.168 | 0.055 | 0.125 | 0.062 |
| FAM-CC-Pent2-hen3 | CC-Pent2-hen3 | 13302 | 13881 | 10377 | 1.071 | 1.123 | 0.806 | 1.000 | 0.170 |
| FAM-CC-Pent2-hen3 | CC-Hex2 | 4814 | 3926 | 3748 | 0.302 | 0.222 | 0.206 | 0.243 | 0.052 |
| FAM-CC-Pent2-hen3 | CC-Hept-IV-hen2 | 2886 | 2623 | 2349 | 0.128 | 0.104 | 0.079 | 0.104 | 0.024 |
| FAM-CC-Hex2 | - | 632 | 499 | 546 | 0.006 | -0.005 | -0.001 | 0.000 | 0.006 |
| FAM-CC-Hex2 | CC-Di | 567 | 508 | 572 | 0.001 | -0.004 | 0.001 | -0.001 | 0.003 |
| FAM-CC-Hex2 | CC-Tri | 615 | 519 | 592 | 0.005 | -0.003 | 0.003 | 0.001 | 0.004 |
| FAM-CC-Hex2 | CC-Tet* | 4064 | 3828 | 4041 | 0.291 | 0.272 | 0.289 | 0.284 | 0.011 |
| FAM-CC-Hex2 | CC-Pent2-hen3 | 3295 | 3210 | 3175 | 0.227 | 0.220 | 0.217 | 0.222 | 0.005 |
| FAM-CC-Hex2 | CC-Hex2 | 13004 | 11976 | 12783 | 1.035 | 0.949 | 1.016 | 1.000 | 0.045 |
| FAM-CC-Hex2 | CC-Hept-IV-hen2 | 784 | 1213 | 859 | 0.019 | 0.054 | 0.025 | 0.033 | 0.019 |
| FAM-CC-Hept-IV-hen2 | - | 1154 | 1172 | 2112 | -0.038 | -0.036 | 0.074 | 0.000 | 0.065 |
| FAM-CC-Hept-IV-hen2 | CC-Di | 1056 | 1201 | 1053 | -0.050 | -0.033 | -0.050 | -0.044 | 0.010 |
| FAM-CC-Hept-IV-hen2 | CC-Tri | 1103 | 907 | 937 | -0.044 | -0.067 | -0.064 | -0.059 | 0.012 |
| FAM-CC-Hept-IV-hen2 | CC-Tet* | 1101 | 1086 | 991 | -0.045 | -0.046 | -0.058 | -0.049 | 0.007 |
| FAM-CC-Hept-IV-hen2 | CC-Pent2-hen3 | 1575 | 1419 | 1447 | 0.011 | -0.007 | -0.004 | 0.000 | 0.010 |
| FAM-CC-Hept-IV-hen2 | CC-Hex2 | 3246 | 2687 | 3311 | 0.208 | 0.142 | 0.216 | 0.189 | 0.040 |
| FAM-CC-Hept-IV-hen2 | CC-Hept-IV-hen2 | 9536 | 9460 | 10919 | 0.949 | 0.940 | 1.112 | 1.000 | 0.097 |
